# Retracing Human Genetic Histories and Natural Selection Using Precise Local Ancestry Inference

**DOI:** 10.1101/2023.09.11.557177

**Authors:** Jon Lerga-Jaso, Biljana Novković, Deepu Unnikrishnan, Varuna Bamunusinghe, Marcelinus R. Hatorangan, Charlie Manson, Haley Pedersen, Alex Osama, Andrew Terpolovsky, Sandra Bohn, Adriano De Marino, Abdallah A. Mahmoud, Karatuğ O. Bircan, Umar Khan, Manfred G. Grabherr, Puya G. Yazdi

## Abstract

In an increasingly diverse world, including admixed individuals in genomic studies is imperative for equity and portability. A crucial first step is precise local ancestry inference (LAI). We have developed Orchestra, a LAI model with unprecedented accuracy, and trained on over 10,000 single-origin individuals from 35 worldwide populations. We employed Orchestra to delve into genetic relationships and demographic histories, with a focus on Latin Americans, a prime example of admixture, and the Ashkenazi Jewish, whose origins have long been debated. Finally, Orchestra enabled us to map signatures of selection, notably identifying trace Scandinavian ancestry in British samples and unveiling an immune-rich region linked to respiratory infections. Our work advances the field of LAI and holds promise for improvements in future applications for admixed populations.

**One-Sentence Summary:** Orchestra unveils Latino and Ashkenazi ancestral roots and a candidate Viking locus under selection in the British population

## Introduction

Despite vast differences in phenotypes, languages, and culture, any two humans living today share over 98% of their DNA (*1*). The remainder, of which only 0.1% is due to SNPs, can tell us the part of the world from which an individual, or their ancestors, originated. For decades, population geneticists have used DNA to infer the history of humankind (*2-4*). Mating between individuals in geographical proximity, coupled with genetic drift and divergent demographic histories, helped shape our modern human genetic landscape, allowing us to reference any single individual to well differentiated reference populations (*5, 6*).

However, as the movement of people across the globe has intensified in recent centuries, humans have become more admixed, and an increasing number of people cannot be traced back to a single reference population (*7*). While global ancestry inference (GAI) allows us to infer an individual’s overall admixture proportions, it fails to provide information about the fine-scale patterns across the genome. Despite similar global admixture proportions, two individuals may have very different ancestry compositions at any location within the genome (*8*). That is why in admixed populations, local ancestry inference (LAI) becomes indispensable for various downstream applications. Several LAI methods that infer the ancestry of different segments on each chromosome have been developed over the years (*9-11*). LAI applied to admixed populations has been used to boost power and resolution in GWAS (*12*), improve GWAS and expression quantitative trait loci (eQTL) colocalization (*13*), and detect gene-gene and gene-environment interactions (*14*). In addition, LAI models have been leveraged to improve polygenic risk scores (PRS) specifically for admixed individuals (*15, 16*). However, these efforts have mainly applied cross-continental resolution (e.g., European *vs*. African or East Asian).

Many human populations, despite being geographically close, are genetically heterogeneous. Such regional variation can impact the genetic architecture of complex phenotypes. In Africa alone, the genomic diversity is vast, showing extreme allele frequency divergence in many medically relevant variants (*17*). Similarly, genomic variability is extensive among Asians, who comprise nearly 60% of the total world population, with unequal genetic disorder burden and pharmacological susceptibility (*18*). Subtle genetic clines can be observed even for Europeans (*19*). In fact, the European North-South gradient in height is one of the best-documented examples of how selective adaptation has shaped complex traits (*20*). Therefore, various genomic disciplines may have a lot to gain from broadening the scope of LAI to include within-continent diversity.

## Results

### Local Ancestry Deconvolution with Orchestra

Here, we present Orchestra (**O**ptimal [**r**e]**c**ombination of **h**aplotypes to **e**stablish **s**egmentation of a **t**arget from **r**eference **a**ncestries), a novel LAI algorithm, and demonstrate its superiority to other state-of-the-art LAI algorithms. We apply Orchestra to retrace the genetic history of Latin Americans, as a prime example of admixture. We next explore the relationship between 35 worldwide populations and show that Orchestra can be used to estimate genetic closeness between populations and shed light on their demographic history. Finally, we use Orchestra to detect natural selection signatures.

Orchestra consists of a two-stage pipeline: a base layer and a smoothing module (Fig. 1A). The base layer classifies genomic windows of predetermined size by generating a distance measure between the target genome and each of the reference populations. This measure, *recombination distance*, is the minimum number of segments needed to reconstruct a target sequence from the sequences present in each reference population. It approximates the number of crossover events needed to reconstruct a given sequence. The base layer uses a greedy approach in which a similarity matrix is calculated by an element-to-element comparison per position and per sample, to obtain a vector of recombination distances across all reference populations. The smoothing module is a deep learning model with convolutional and attention-based elements. The convolutional element processes the base layer insights generated for each window using the information from surrounding windows. The attention-based component provides a weak link to global ancestry. This is reflective of real world genomes, since the presence of a certain ancestry in one place of the genome increases the likelihood of finding that same ancestry in other genomic regions. Combining the recombination distance base layer with a deep learning smoothing module synergistically leads to a novel, state-of-the-art technique for accurate ancestry deconvolution.

**Fig. 1.**
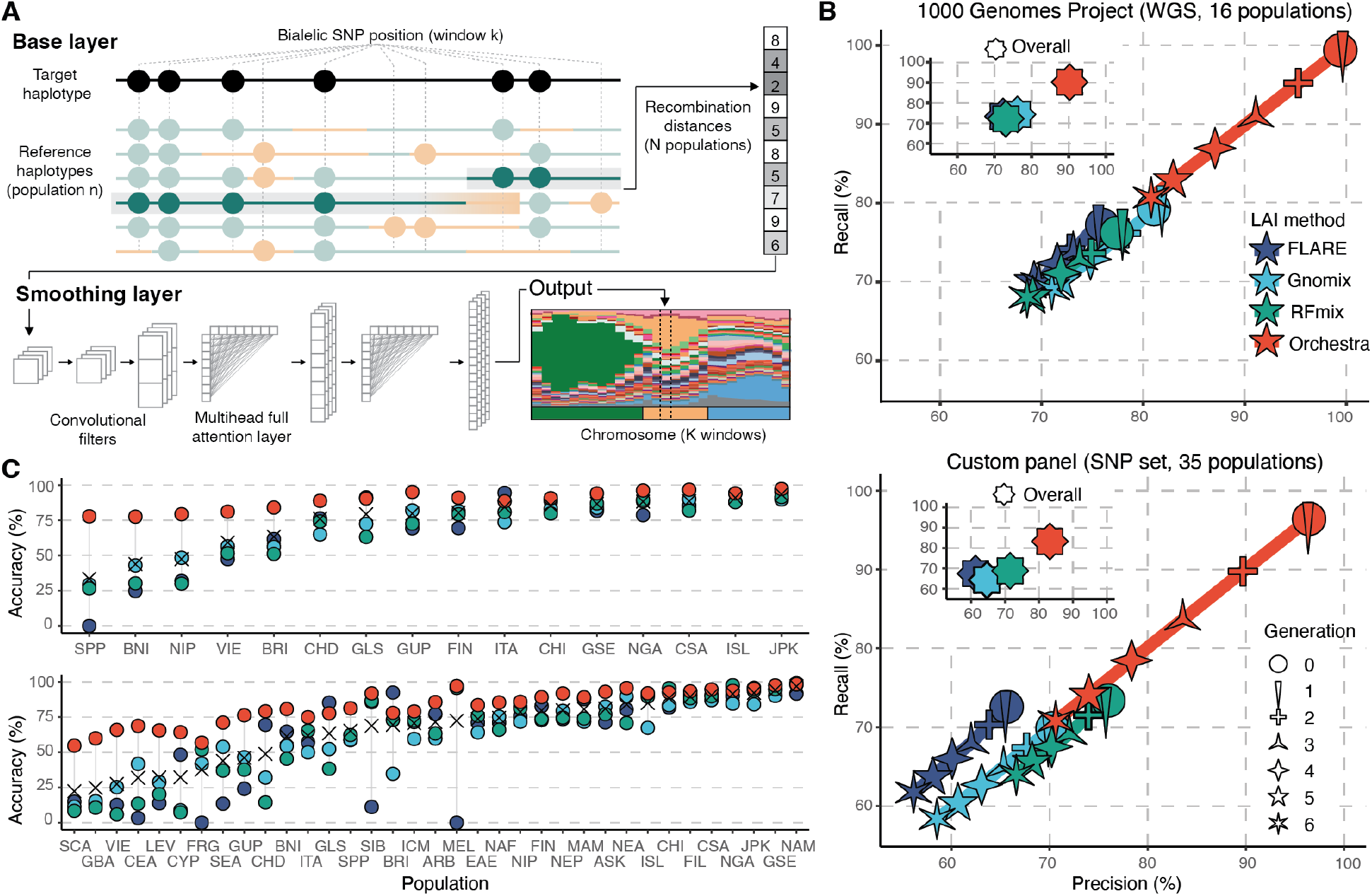
Performance of Orchestra and other LAI methods. (**A**) Orchestra schematics. The input consists of a target genome whose ancestry is unknown and a reference set of single-origin individuals grouped by population. For each genomic window, the base layer generates the *recombination distance* as the minimum number of segments needed to reconstruct the target from the sequences of each reference population *n* (filled circles represent alternative alleles; green circles show correspondence between the reference genomes and the target). The smoothing layer then processes these distance measures with a series of convolutional and attention layers using information from surrounding windows and other regions of the genome, yielding the final probabilities for the *N* reference ancestries. This procedure is repeated for all windows in the genome. (**B**) Recall and precision of Orchestra, RFmix, Gnomix and FLARE across 6 generations. Star shapes refer to the number of generations of simulated admixture (the more points the star has, the higher the generation). (**C**) Accuracy (%) per population for the 16 populations in the 1KGP dataset and the 35 populations in the larger custom dataset. Populations are ordered by mean accuracy across all methods (cross). Accuracy is shown as synonymous with recall.

The accuracy of any ancestry model greatly depends on the quality of the reference panel. We assembled a set of reference populations by merging data from more than 30 published studies, combining both whole genome sequencing and array-based genotyping (table S1). A significant fraction of the total samples comes from non-UK ancestries captured by the UK Biobank (UKBB). With much shorter migratory distances just a few decades ago, we found that tracing ancestral origins by birth-place and self-reported ethnicity of UKBB participants was a sufficiently reliable proxy for ancestry (figs. S1-3). All retrieved samples underwent a series of quality filtering steps. We kept a composite set of directly genotyped variants obtained by combining all SNPs from array-based studies and filtered by a minor allele frequency (MAF) ≥ 5% to minimize imputation-related biases (see Methods). Next we conducted two GWASs to check if each SNP was associated with a genotyping platform or ancestry, and filtered out those that ranked in the top high and low end, respectively, to minimize batch effects and retain meaningful ancestry informative differences. We then used two separate dimensionality reduction techniques to characterize relationships between samples and remove any samples that showed a disagreement between reported ancestry and inferred genetic origin: 1) Principal component analysis (PCA) followed by uniform manifold approximation and projection (UMAP) (*21*) and 2) t-distributed stochastic neighbor embedding (t-SNE) (*22*) used on genealogical nearest neighbor (GNN) statistics estimated with tsinfer (*5*). This resulted in a high-quality reference panel of 10,169 non-admixed individuals from 35 world regions, which we used as our reference populations (fig. S4; see table S2 for three-letter population abbreviations; see Methods for more details).

We benchmarked Orchestra against other leading LAI algorithms, including RFmix (*9*), Gnomix (*10*) and FLARE (*11*), using two reference panels: 1) 1KGP-16pops, a high-coverage WGS set of non-admixed and unrelated samples collected by the 1000 Genomes Project (1KGP) with 16 populations and 2) custom-35pop, our larger, more diverse curated panel with 35 populations. Both panels were split into test and training sets (20% and 80% of samples) and used to simulate 6 generations of random admixture using SLiM (*23*). Precision and recall were reported as performance estimates on all chromosomes per generation and per population.

Orchestra substantially outperformed other LAI methods (Fig. 1B). When using the 1KGP-16pops reference panel, Orchestra’s average recall and precision across generations was 90.17% and 90.22%, respectively; an improvement of +15.89% and +14.03% compared to the second best model, Gnomix. For the custom-35pops panel, the average recall and precision was 79.54% and 80.54%, respectively, an improvement of +15.04% and +13.99% compared to the next best model, RFmix. Orchestra was the most accurate across 6 generations of admixture. As expected, the accuracy decreased with an increasing number of generations. However Orchestra’s performance in the most admixed samples equaled or exceeded the best performance in the non-admixed generations by other LAI methods.

Orchestra retained high accuracy regardless of the reference population, with an ability to distinguish between closely related ancestries. Orchestra achieved accuracy greater than 75% for all populations within the 1KGP-16pops panel (Fig. 1C). For the custom-35pops panel, Orchestra achieved an accuracy of over 50% for all populations, and over 75% for 26 out of 35 populations. The other three LAI models struggled with a third of the populations, with accuracy below 50% (Fig. 1C). Orchestra’s accuracy was superior at both region-wide and continental levels, the recall exceeding 93.43 and 98.90% for 1KGP-16pops and 87.73% and 94.03% for custom-35pops (figs. S5-8).

In addition to our two panels, we applied all LAI models to over 10,000 UK biobank samples that were not included in the custom-35pops panel (fig. S9). Orchestra outperformed the other LAI methods for 91% of the 103 evaluated countries.

### Retracing Genetic Histories

Latin Americans are a prime example of admixture, as their DNA can be traced to three broad sources, European, Sub-Saharan African and Amerindian. However, there is a wide fluctuation in the proportion of these ancestries throughout the continent. In addition, the genetic makeup of Latinos shows regional heterogeneity. For example, Colombians are more likely to have Senegalese, Gambian or Guinean African ancestry, while Brazilians are more likely to have ancestry from Angola and Congo. Similarly, while a lot of Latin Americans get their European ancestry from Spain or Portugal, many Argentinians also have Italian roots (*24*).

To assess the accuracy of our LAI model in these populations, we simulated Latino individuals from Southern (SPP and ITA) and Northern (FRG and BRI) Europeans, Western (GSE, GLS and NGA) and Central and Southern (CSA) Africans and artificially-reconstructed Native Americans (NAM). Simulations were performed by emulating genetic intermixing for 12 generations using SLiM (*23*). Native American genomes were created *in silico* using the Latino samples from the 1KGP, keeping only the genomic segments identified as East Asian as a proxy for indigenous ancestry (fig. S10). Simulations were adapted to the genetic makeup that can be found today in three broad regions within Latin America: the Antilles, comprised of 55% European, 40% African (specifically NGA) and 5% Native American ancestry (NAM); Mexico and Central America, made up of 50% European, 10% African (GLS and GSE) and 40% Native American ancestry (NAM); and South America, composed of 65%, 15% and 20% of European, African (GLS, GSE and CSA) and Native American (NAM), respectively (*24*; fig. S11). For benchmarking purposes, we compared our results against FLARE, Gnomix and RFmix (Fig. 2A). Orchestra achieved an overall precision and recall of 77.17% and 76.73%, respectively, outperforming the other three LAI models in all three aforementioned regions.

**Fig. 2.**
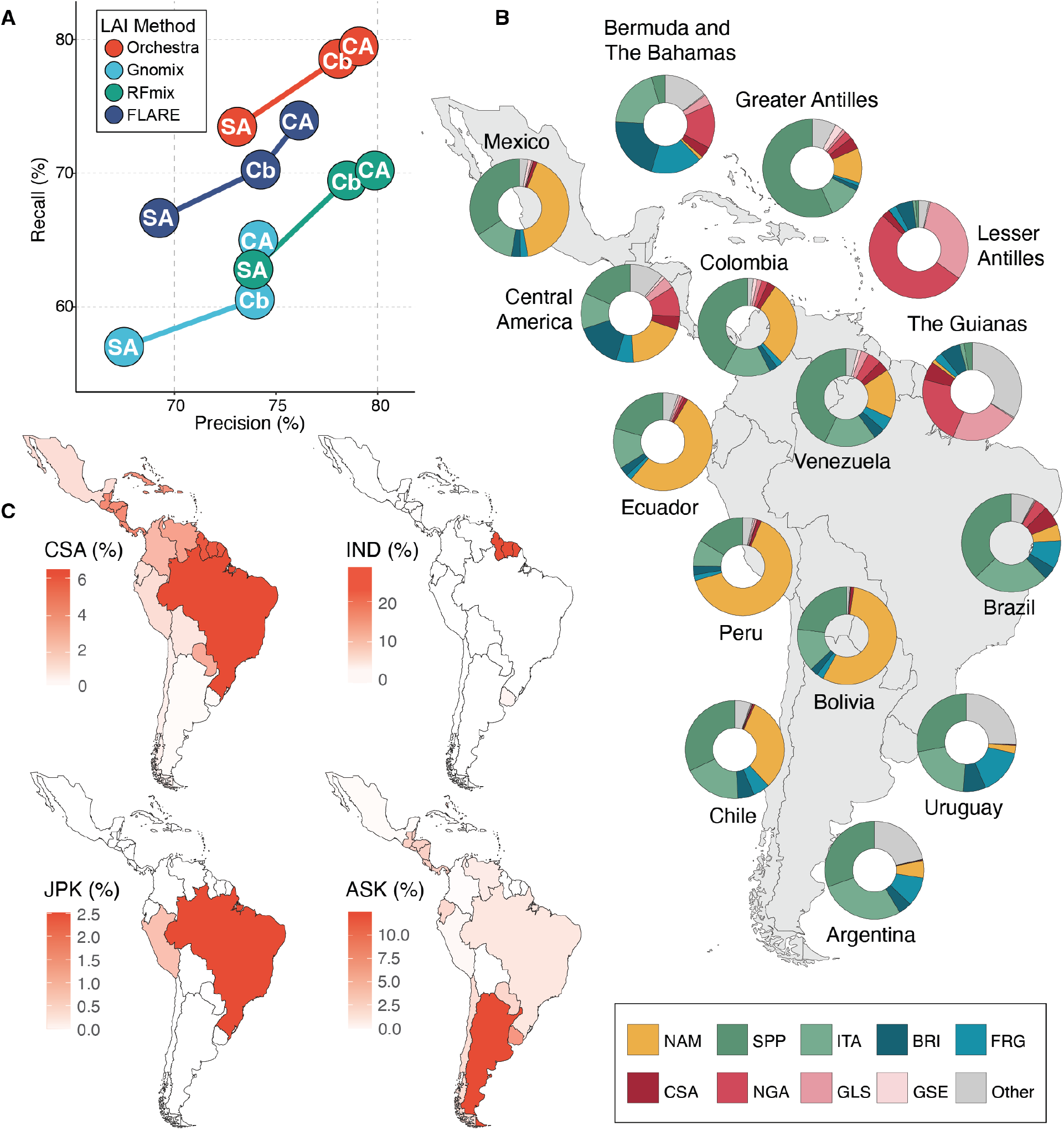
Orchestra’s performance in Latino American individuals. (**A**) Ancestral composition of 1KGP admixed American populations and UKBB participants that were born in the Americas revealed by Orchestra. Proportions of Native American (yellow), Southern European (green), Northern European (blue) and African (red) ancestries are shown. (**B**) Orchestra was also able to detect trace ancestries that reflect known historical population displacements and immigration events. Clockwise: Greater percentage of Central, South & Southeast African (CSA) ancestry in Brazil, A relatively large percentage of various Indian ancestries in the Guianas (IND), greater percentage of Ashkenazi Jewish (ASK) in Argentina, and Japanese & Korean (JPK) ancestry in Brazil and Peru. (**C**) Percent recall and precision for ancestry deconvolution by FLARE (navy), Gnomix (light blue), RFmix (green), and Orchestra (red) on Latinos simulations (equivalent to 12 generations of admixture; we adjusted the simulations to mimic the actual genetic composition of different regions within the continent: CA = Central America, Cb = Caribbean, SA = South America).

This gave us confidence to apply our model to real life Latin American samples from the 1KGP and UKBB datasets (Fig. 2B, fig. S12), where Orchestra was able to successfully retrace major patterns in the genetic history of the Latin Americas. For example, the highest percentage of Native American ancestry (NAM) was found in Bolivia, Peru, Ecuador and Mexico, matching demographic and genetic reports from this region (*24-26*). The majority of African ancestry in the Caribbeans was assigned to Nigerian (NGA) and next Ghanaian, Ivorian, Liberian & Sierra Leonean (GLS) ancestry. In contrast, a larger portion of the African ancestry in Brazil was assigned to Central, South & Southeast African (CSA) ancestry, which captures populations of Bantu origin on the African continent. This is in agreement with historical records of Africans being transported to Brazil primarily from Angola, a former Portuguese colony (*24*).

Orchestra captured a higher percentage of Spanish & Portuguese (SPP) ancestry in Mexico, the Greater Antilles, Columbia and Venezuela. British & Irish ancestry (BRI) was more prevalent in Bermuda and the Bahamas, Lesser Antilles and the Guianas. Italian (ITA) ancestry was more prominent in Argentina, Brazil and Uruguay. These findings match known demographic and historic evidence (*27*).

Interestingly, Orchestra was able to detect several notable trace ancestries (Fig. 2C). In addition to the aforementioned CSA in Brazil, these also include a high percentage of Indian ancestries (IND = BNI, GUP, ISL, NIP) in the Guianas, reflecting the indenture system used in former British colonies (*28*), a high percentage of Ashkenazi Jewish (ASK) ancestry in Argentina, which hosts the largest Ashkenazi Jewish community in South America, as well as the Japanese ancestry detected in Brazil and Peru (JPK), which experienced a large wave of Japanese immigration over the first half of the 20th century (*29*).

We also applied this method to all UKBB samples not used in our reference panel (fig. S13) and to samples belonging to ethnicities found in various datasets that were not included in the custom-35pops panel (figs. S14-25).

### Ancestral Mapping

Seeing that we were able to identify 35 different populations with unprecedented accuracy (Fig. 1), next we explored the relationships among these populations. We created 35 distinct reference panels for each target population, where that target population was omitted from its own reference panel. Orchestra was run to obtain admixture proportions, which were then converted into a matrix of distances that were projected onto two-dimensional space using the SMACOF algorithm. The resulting network (Fig. 3A) largely reflects geographical proximity and replicates known relationships between various populations (*19, 30*).

**Fig. 3.**
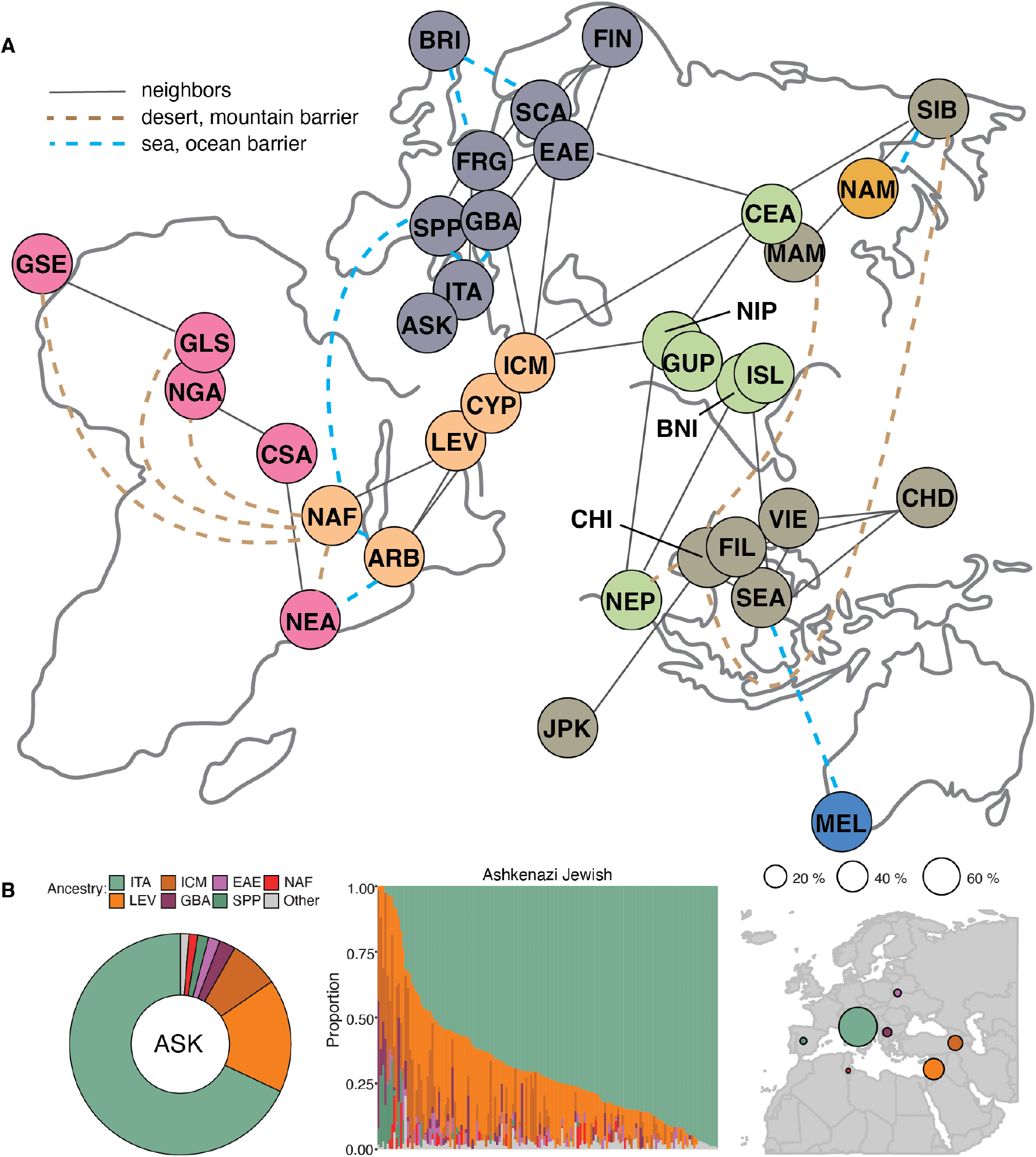
Ancestral mapping. (**A**) Ancestral map of 35 populations projected onto two-dimensional space using the SMACOF algorithm applied to a matrix of distances derived from the proportions obtained in the analysis. Solid lines connect neighboring populations. Dashed lines indicate populations separated by geographical barriers to dispersion. Continent contours were illustrated by hand. (**B**) Inferred ancestry of the Ashkenazi Jewish (ASK) population according to Orchestra when ASK was omitted from the custom-35-pops training set. ARB: Arab, ASK: Ashkenazi Jewish, BRI: British & Irish, BNI: Bengali & East Indian, CEA: Central Asian, CHD: Chinese Dai, CHI: Han Chinese, CSA: Central, South & Southeast African, CYP: Cypriot, EAE: Eastern European, FIL: Filipino, FIN: Finish, FRG: French & German, GBA: Greek & Balkan, GLS: Ghanaian, Ivorian, Liberian & Sierra Leonean, GSE: Gambian & Senegalese, GUP: Gujarati Patels, ICM: Turkish, Iraqi, Iranian & Caucasian, ISL: Southern Indian & Sri Lankan, ITA: Italian, JPK: Japanese & Korean, LEV: Levantine, MAM: Manchurian & Mongolian, MEL: Melanesian & Aboriginal Australian, NAF: North African, NAM: Native American, NEA: Northeast African, NEP: Nepalese, NGA: Nigerian, NIP: North Indian & Pakistani, SCA: Scandinavian, SEA: Southeast Asian, SIB: Siberian, SPP: Spanish & Portuguese, VIE: Vietnamese.

Ancestral mapping results for individual populations are shown in figures S26-60. For example, when we removed our French & German (FRG) population from the reference panel, the FRG samples were mapped as mostly British and Irish (BRI, 58.9 %), Scandinavian (SCA, 13.1 %), Italian (ITA, 9.5 %) and Eastern European (EAE, 9.4 %), with the ITA ancestry more prevalent in the French and the Swiss, while EAE ancestry was more common in Austrians and Germans (fig. S32). Our Turkish, Iraqi, Iranian & Caucasian population (ICM) was reconstructed as mostly Levantine (LEV, 65.2 %), North Indian & Pakistani (NIP, 11.9 %), Cypriot (CYP, 9.6 %), Central Asian (CEA, 6.4 %) and Greek & Balkan (GBA, 3.8 %). NIP ancestry was more common in the East, in Iraqis and Iranians, while GBA was more present in the West, especially in the Turks (fig. S44). In these and many other populations, we observed a genetic cline, indicating there is genetic heterogeneity within most of our 35 populations.

The ancestry and origin of the Askenazi Jewish have been subject to heated debate over the last two decades. Here we mapped our Ashkenazi Jewish as primarily Italian (ITA, 68 %), followed by Levantine (LEV, 16.6 %), Turkish, Iraqi, Iranian or Caucasian (ICM, 7.2 %), Greek and Balkan (GBA, 2.4 %) and Eastern European (EAE, 1.7 %) (Fig. 3B). This largely agrees with several reports based on both modern and medieval Ashkenazi Jewish DNA (*31-33*).

### Detecting Signatures of Natural Selection

To check if we could leverage our LAI model to detect signatures of natural selection, we first aimed to replicate previously identified signals. We followed the methods described in Cuadros-Espinosa *et al*. (2022) (*34*), where combined statistics based on admixture proportions (Fadm and LAD) was used to scan genomes of admixed populations for selection signals. Out of the seven admixed populations tested, we were able to completely replicate signals in four populations and partially in one population (table S3, fig. S61). Some of the discrepancies may be due to methodology. We used a reference panel with 35 different populations to detect admixture, *vs*. only the populations involved in the admixture in the original study. However, the fact that many of the signals were detectable, even when we used different datasets with fewer samples suggests that these signals are robust to discovery with different methods. This also suggests that Orchestra may be used to recover signals of natural selection at a local level.

We then proceeded to apply Orchestra to British samples (N = 415,859) in the UK biobank dataset. Figure 4A shows the distribution of Scandinavian (SCA) ancestry in this population. SCA ancestry was particularly enriched in the East of England and East Midlands, where we also found the highest density of former tentative Viking settlements, inferred as settlement names ending in -by, -thorpe or -toft, confirming previous reports of Scandinavian hotspots in Eastern England (*35, 36*). Next we aimed to identify potential adaptive signals using the Fadm and LAD framework. We found a significant enrichment of SCA ancestry on chromosome 10, region 10q11.21-22 (Fig. 4B-C, fig. S62). We identified significant variants in this region that have been functionally linked to several immune-related genes and potential targets for natural selection, including *MAPK8, WASHC2C* and *MARCH8*. Interestingly, both *MAPK8* and *WASHC2C* have been linked to smallpox virus infection and replication rates (*37, 38*), which is of note considering that the Vikings were reported to be carriers of smallpox-like viruses (*39*). Apart from smallpox, these genes have also been linked to influenza, bacterial pneumonia, tuberculosis (*40*), and other infectious diseases prominent in Middle Age Britain (*41*). Furthermore, the region displays an enrichment of GWAS hits where SCA ancestry is associated with elevated erythrocyte and hemoglobin levels (fig. S63), and there is a higher prevalence of SCA ancestry among UKBB participants reporting lower incidences of “respiratory infection” and “influenza with pneumonia” (fig. S64).

**Fig. 4.**
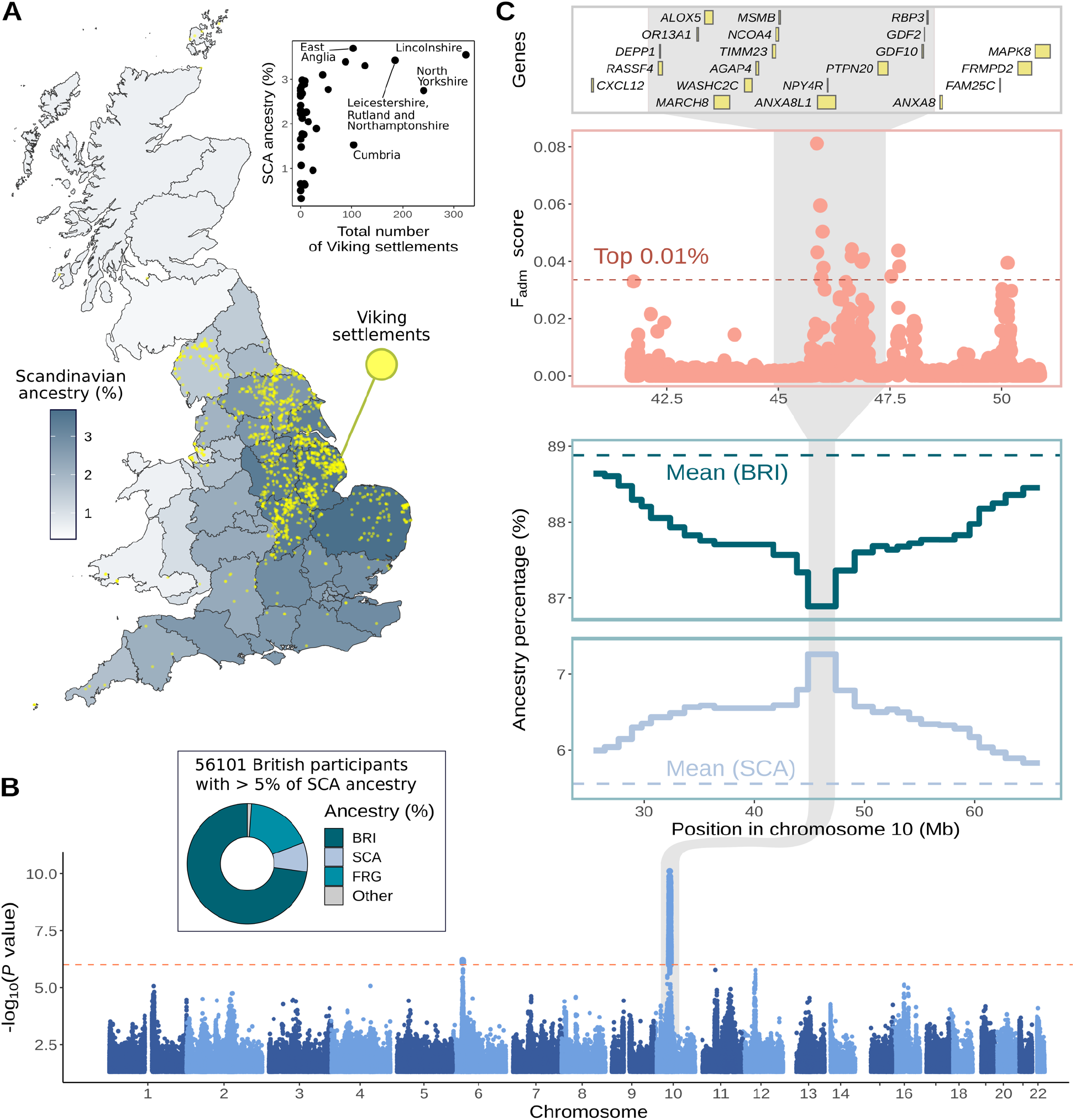
Signatures of Scandinavian ancestry and natural selection in British UK biobank participants. (**A**) Scandinavian (SCA) ancestry percentage in UK and presumed Viking settlements, inferred from place name endings: -by, -thorpe, or -toft. Inset shows the correlation between the total places with Viking-derived names and average SCA ancestry per county. (**B**) Genome-wide adaptive admixture signals in 56,101 British individuals with over 5% SCA ancestry. Larger points indicate variants surpassing the significance threshold (denoted by a horizontal dotted line). The chr10 signal detected is marked by a gray-shaded area. Inset, estimated ancestral composition for this sample set. (**C**) Local percentages of BRI and SCA ancestries on chromosome 10 windows (average percentage levels depicted by horizontal lines), alongside the Fadm score per variant and genes within the region.

## Discussion

While our world is becoming increasingly admixed, genomic studies have largely focused on non-admixed populations with a pronounced European bias (*8, 41, 45*). This is an issue when we consider that genomic models developed and trained in one population have poor portability outside that population (*45-47*). Recently, strides have been made to address this gap, but most have been limited to cross-continental resolution, due to limitations in LAI accuracy (*12-16*).

To address this, we have developed Orchestra, a novel LAI model that can account for the genetic heterogeneity in admixed populations. Orchestra can work with reference panels made of many combined datasets, is able to accurately retrace demographic histories of complex admixed populations, such as Latin Americans, and does so on a fine-grained regional scale. Further, Orchestra can be used to elucidate relationships between different populations. We weigh in on the ongoing debate about the origins of the Ashkenazi Jewish, supporting strong genetic ties to the Italian Peninsula (*31-33*).

Finally, Orchestra’s local aspect enables us to apply it to downstream applications, such as detecting signatures of selection. We trace Scandinavian ancestry in British UKBB samples, which allows us to detect a potential immune-related signal on chromosome 10. This possibly Viking-derived region may have provided an edge against respiratory infections, such as smallpox, influenza or pneumonia. This region is, to this day, linked to a lower rate of respiratory infections and influenza in UKBB participants.

There is potential for improving the accuracy of Orchestra by creating more sophisticated reference panels. Some of the issues we had to overcome in this study were batch effects due to diverse sequencing technologies and insufficient coverage in more dated datasets. No reference panel can be perfect, as we are by definition breaking up genomic continuums into discrete populations.We expect that improving the reference panel will generate further finer-scale insights into recent admixture around the globe.

A limitation of Orchestra in its current iteration is its use of windows to infer local ancestry. With admixture, genomic segments become increasingly smaller with each generation due to cross-over and recombination. And while Orchestra is modeled on recombination to reconstruct ancestry within a window, this results in a trade-off between accuracy and the ability to detect signals from further back in time. Orchestra is relatively accurate at least up to around 12 generations of admixture and can detect trace ancestries from further back in time. This means it is suited to reconstruct events of relatively recent admixture, within Modern and potentially Medieval history. However, for reconstruction of Ancient history, other non-window based LAI models would have a clear advantage (*48, 49*).

Orchestra advances the field of LAI, enabling accurate detection of chromosomal segments at regional levels. It also takes an important step towards a more equitable genomics, promising to improve a range of downstream applications, such as long-range phasing and GWAS, and by extent genomic and personalized medicine in admixed populations.

## Acknowledgments

We thank the team and our colleagues at Omics Edge and Genius Labs. This research was conducted by using the UK Biobank Resource under application number 84038.

## Funding

All work was funded by a commercial source, Omics Edge, a subsidiary of Genius Labs Company. Omics Edge provided only funding for the study, but had no additional role in study design, data collection and analysis, decision to publish or preparation of the manuscript beyond the funding of the contributors’ salaries.

## Author contributions

Conceptualization: J.L.J., B.N. and P.G.Y. Data curation: J.L.J., B.N., V.B., M.R.H., H.P., S.B., A.D.M., A.A.M. and K.O.B. Formal Analysis: J.L.J., B.N., D.U., V.B., M.R.H., C.M., H.P., A.O., A.T. and M.G.G. Methodology: J.L.J., B.N., D.U., V.B., C.M., U.K., M.G.G. and P.G.Y. Investigation: J.L.J., B.N., D.U., V.B., M.R.H., C.M., H.P., A.O. and M.G.G. Software: D.U., V.B., C.M., A.O., A.T., S.B., A.D.M., A.A.M., K.O.B., U.K., M.G.G. Visualization: J.L.J., B.N. Funding acquisition: P.G.Y. Project administration: P.G.Y. Resources: P.G.Y. Supervision: P.G.Y. Writing – original draft: J.L.J. and B.N. Writing – review & editing: J.L.J., B.N. and P.G.Y.

## Competing interests

All authors are either employed by and/or hold stock or stock options in Genius Labs. In addition, P.G.Y. has equity in Systomic Health LLC and Ethobiotics LLC. This work has been used to file a provisional patent application. There are no other relevant activities or financial relationships which have influenced this work.

## Data and materials availability

All publicly available datasets used in this paper are available from their original publications and via application to UK biobank and dbGAP. Orchestra is available for non-commercial use via the corresponding author.

## References and Notes

1. A. W. Pang et al., Towards a comprehensive structural variation map of an individual human genome. Genome Biol. 11, R52 (2010). doi: 10.1186/gb-2010-11-5-r52; pmid: 20482838

2. R. L. Cann, M. Stoneking, A. C. Wilson, Mitochondrial DNA and human evolution. Nature 325, 31–36 (1987). doi: 10.1038/325031a0; pmid: 3025745

3. N. A. Rosenberg et al., Genetic structure of human populations. Science. 298, 2381–2385 (2002). doi: 10.1126/science.1078311; pmid: 12493913

4. R. Nielsen et al., Tracing the peopling of the world through genomics. Nature. 541, 302–310 (2017). doi: 10.1038/nature21347; pmid: 28102248

5. J. Kelleher et al., Inferring whole-genome histories in large population datasets. Nat. Genet. 51, 1330–1338 (2019). doi: 10.1038/s41588-019-0483-y; pmid: 31477934

6. A. Bergström et al., Insights into human genetic variation and population history from 929 diverse genomes. Science 367, eaay5012 (2020). doi: 10.1126/science.aay5012; pmid: 32193295

7. A. L. Atkin, N. K. Christophe, G. L. Stein, A. K. Gabriel, R. M. Lee, Race terminology in the field of psychology: Acknowledging the growing multiracial population in the U.S. Am. Psychol. 77, 381–393 (2022). doi: 10.1037/amp0000975; pmid: 35254853

8. T. Tan, E. G. Atkinson. Strategies for the Genomic Analysis of Admixed Populations. Annu. Rev. Biomed. Data Sci. 6, 105–127 (2023). doi: 10.1146/annurev-biodatasci-020722-014310; pmid: 37127050

9. B. K. Maples, S. Gravel, E. E. Kenny, C. D. Bustamante, RFMix: a discriminative modeling approach for rapid and robust local-ancestry inference. Am. J. Hum. Genet. 93, 278–288 (2013). doi: 10.1016/j.ajhg.2013.06.020; pmid: 23910464

10. H. Hilmarsson et al., High Resolution Ancestry Deconvolution for Next Generation Genomic Data. bioRxiv 2021.09.19.460980 [Preprint] (2021). 10.1101/2021.09.19.460980 [bioRxiv]

11. S. R. Browning, R. K. Waples, B. L. Browning, Fast, accurate local ancestry inference with FLARE. Am. J. Hum. Genet. 110, 326 –335 (2023). doi: 10.1016/j.ajhg.2022.12.010; pmid: 36610402

12. E. G. Atkinson et al., Tractor uses local ancestry to enable the inclusion of admixed individuals in GWAS and to boost power. Nat. Genet. 53, 195–204 (2021). doi: 10.1038/s41588-020-00766-y; pmid: 33462486

13. N. R. Gay et al., Impact of admixture and ancestry on eQTL analysis and GWAS colocalization in GTEx. Genome Biol. 21: 233 (2020). doi: 10.1186/s13059-020-02113-0; pmid: 32912333

14. R. A. Patel et al., Genetic interactions drive heterogeneity in causal variant effect sizes for gene expression and complex traits. Am. J. Hum. Genet. 109, 1286–1297 (2022). doi: 10.1016/j.ajhg.2022.05.014; pmid: 35716666

15. Q. Sun et al., Improving polygenic risk prediction in admixed populations by explicitly modeling ancestral-specific effects via GAUDI. bioRxiv 2022.10.06.511219 [Preprint] (2021). 10.1101/2022.10.06.511219 [bioRxiv]

16. D. Marnetto et al., Ancestry deconvolution and partial polygenic score can improve susceptibility predictions in recently admixed individuals. Nat. Commun. 11, 1628 (2020). doi: 10.1038/s41467-020-15464-w; pmid: 32242022

17. A. Choudhury et al., High-depth African genomes inform human migration and health. Nature 586, 741–748 (2020). doi: 10.1038/s41586-020-2859-7; pmid: 33846614

18. S. H. Chan et al., Analysis of clinically relevant variants from ancestrally diverse Asian genomes. Nat. Commun. 13, 6694 (2022). doi: 10.1038/s41467-022-34116-9; pmid: 36335097

19. J. Novembre et al., Genes mirror geography within Europe. Nature 456, 98–101 (2008). doi: 10.1038/nature07331; pmid: 18758442

20. I. Mathieson et al., Genome-wide patterns of selection in 230 ancient Eurasians. Nature 528, 499–503 (2015). doi: 10.1038/nature07331; pmid: 26595274

21. L. McInnes, J. Healy, J. Melville, Uniform Manifold Approximation and Projection for dimension reduction. arxiv.org/abs/1802.03426 (2018). [arXiv]

22. L. van der Maaten, G. Hinton, Visualizing data using t-sne. J. Mach. Learn. Res. 9, 2579–2605 (2008). https://www.jmlr.org/papers/v9/vandermaaten08a.html

23. B. C. Haller, P. W. Messer, SLiM 3: Forward Genetic Simulations Beyond the Wright-Fisher Model. Mol. Biol. Evol. 36, 632–637 (2019). doi: 10.1093/molbev/msy228; pmid: 30517680

24. L. Ongaro et al., The Genomic Impact of European Colonization of the Americas. Curr. Biol. 29, 3974–3986.e4 (2019). doi: 10.1016/j.cub.2019.09.076; pmid: 31735679

25. J. C. Chacón-Duque et al., Latin Americans show wide-spread Converso ancestry and imprint of local Native ancestry on physical appearance. Nat. Commun. 9, 5388 (2018). doi: 10.1038/s41467-018-07748-z; pmid: 30568240

26. A. Ruiz-Linares et al., Admixture in Latin America: geographic structure, phenotypic diversity and self-perception of ancestry based on 7,342 individuals. PLoS Genet. 10:e1004572 (2014). doi: 10.1371/journal.pgen.1004572; pmid: 25254375

27. J. R. Homburger et al., Genomic Insights into the Ancestry and Demographic History of South America. PLoS Genet. 11, e1005602 (2015). doi: 10.1371/journal.pgen.1005602; pmid: 26636962

28. L. Roopnarine, East Indian indentured emigration to the Caribbean: Beyond the push and pull model. Caribbean studies 31, 97–134 (2003). http://www.jstor.org/stable/25613409

29. J. Durand, D. S. Massey, New World Orders: Continuities and Changes in Latin American Migration. Ann. Am. Acad. Pol. Soc. Sci. 630, 20–52 (2010). doi: 10.1177/0002716210368102; pmid: 20814591

30. B. M. Peter, D. Petkova, J. Novembre, Genetic Landscapes Reveal How Human Genetic Diversity Aligns with Geography. Mol. Biol. Evol. 37, 943–951 (2020). doi: 10.1093/molbev/msz280; pmid: 31778174

31. M. D. Costa et al., A substantial prehistoric European ancestry amongst Ashkenazi maternal lineages. Nat. Commun. 4, 2543 (2013). doi: 10.1038/ncomms3543; pmid: 24104924

32. S. Waldman et al., Genome-wide data from medieval German Jews show that the Ashkenazi founder event pre-dated the 14^th^ century. Cell 185, 4703–4716.e16.(2022). doi: 10.1016/j.cell.2022.11.002; pmid: 36455558

33. J. Xue, T. Lencz, A. Darvasi, I. Pe’er, S. Carmi, The time and place of European admixture in Ashkenazi Jewish history. PLoS Genet. 13, e1006644 (2017). doi: 10.1371/journal.pgen.1006644; pmid: 28376121

34. S. Cuadros-Espinoza, G. Laval, L. Quintana-Murci, E. Patin, The genomic signatures of natural selection in admixed human populations. Am. J. Hum. Genet. 109, 710–726 (2022). doi: 10.1016/j.ajhg.2022.02.011; pmid: 35259336

35. J. Gretzinger et al., The Anglo-Saxon migration and the formation of the early English gene pool. Nature 610, 112–119 (2022). doi: 10.1038/s41586-022-05247-2; pmid: 36131019

36. S. Leslie et al., The fine-scale genetic structure of the British population. Nature 519, 309–314 (2015). doi: 10.1038/nature14230; pmid: 25788095

37. J. Chen et al., The Roles of c-Jun N-Terminal Kinase (JNK) in Infectious Diseases. Int. J. Mol. Sci. 22, 9640 (2021). doi: 10.3390/ijms22179640; pmid: 34502556

38. C. Y. Huang et al., A novel cellular protein, VPEF, facilitates vaccinia virus penetration into HeLa cells through fluid phase endocytosis. J. Virol. 82, 7988–7999 (2008). doi: 10.1128/JVI.00894-08; pmid: 18550675

39. B. Mühlemann et al., Diverse variola virus (smallpox) strains were widespread in northern Europe in the Viking Age. Science 369, eaaw8977 (2020). doi: 10.1126/science.aaw8977; pmid: 32703849

40. J. Robb, C. Cessford, J. Dittmar, S. A. Inskip, P. D. Mitchell, The greatest health problem of the Middle Ages? Estimating the burden of disease in medieval England. Int. J. Paleopathol. 34, 101–112 (2021). doi: 10.1016/j.ijpp.2021.06.011; pmid: 34237609

41. K. L. Korunes, A. Goldberg, Human genetic admixture. PLoS Genet. 17, e1009374 (2021). doi: 10.1371/journal.pgen.1009374; pmid: 33705374

42. J. Mendoza-Revilla et al., Disentangling Signatures of Selection Before and After European Colonization in Latin Americans. Mol. Biol. Evol. 39, msac076 (2022). doi: 10.1093/molbev/msac076; pmid: 35460423

43. J. C. Mychaleckyj et al., Genome-Wide Analysis in Brazilians Reveals Highly Differentiated Native American Genome Regions. Mol. Biol. Evol. 34, 559–574 (2017). doi: 10.1093/molbev/msw249. pmid: 28100790

44. J. Yu Cheng, A. J. Stern, F. Racimo, R. Nielsen, Detecting Selection in Multiple Populations by Modeling Ancestral Admixture Components. Mol. Biol. Evol. 39, msab294 (2022). doi: 10.1093/molbev/msab294. pmid: 34626111

45. A. R. Martin et al., Clinical use of current polygenic risk scores may exacerbate health disparities. Nat. Genet. 51, 584–591 (2019). doi: 10.1038/s41588-019-0379-x; pmid: 30926966

46. G. M. Belbin et al., Toward a fine-scale population health monitoring system. Cell 184, 2068–2083.e11(2021). doi: 10.1016/j.cell.2021.03.034; pmid: 33861964

47. F. Privé et al., Portability of 245 polygenic scores when derived from the UK Biobank and applied to 9 ancestry groups from the same cohort. Am. J. Hum. Genet. 109, 12–23 (2022). doi: 10.1016/j.ajhg.2021.11.008; pmid: 34995502

48. M. Chintalapati, N. Patterson, P. Moorjani, The spatiotemporal patterns of major human admixture events during the European Holocene. Elife 11, e77625 (2022). doi: 10.7554/eLife.77625; pmid: 35635751

49. P. Wangkumhang, M. Greenfield, G. Hellenthal, An efficient method to identify, date, and describe admixture events using haplotype information. Genome Res. 32, 1553–1564 (2022). doi: 10.1101/gr.275994.121; pmid: 35794007

50. A. Krizhevsky, I. Sutskever, G. Hinton, Imagenet classification with deep convolutional neural networks. Commun ACM. 60, 84–90 (2017). doi: 10.1145/3065386

51. A. Vaswani et al., Attention is all you need. Adv. Neural Inf. Process. Syst., 30, 5998–6008 (2017).

52. S. Purcell et al., PLINK: a tool set for whole-genome association and population-based linkage analyses. Am. J. Hum. Genet. 81, 559–575 (2007). doi: 10.1086/519795; pmid: 17701901

53. Picard v.2.26.7 v.3.0.0 (The Broad Institute Picard, 2020 & 2021). http://broadinstitute.github.io/picard/

54. B. L. Browning, Y. Zhou, S. R. Browning, A One-Penny Imputed Genome from Next-Generation Reference Panels. Am. J. Hum. Genet. 103, 338–348 (2018). doi: 10.1016/j.ajhg.2018.07.015; pmid: 30100085

55. M. Byrska-Bishop et al., High-coverage whole-genome sequencing of the expanded 1000 Genomes Project cohort including 602 trios. Cell 185, 3426–3440.e19 (2022). doi: 10.1016/j.cell.2022.08.004; pmid: 36055201

56. D. M. Behar et al., The genome-wide structure of the Jewish people. Nature 466, 238–242 (2010). doi: 10.1038/nature09103; pmid: 20531471

57. K. Tambets et al., Genes reveal traces of common recent demographic history for most of the Uralic-speaking populations. Genome Biol. 19, 139 (2018). doi: 10.1186/s13059-018-1522-1; pmid: 30241495

58. A. Manichaikul et al., Robust relationship inference in genome-wide association studies. Bioinformatics 26, 2867–2873 (2010). doi: 10.1093/bioinformatics/btq559; pmid: 20926424

59. 1000 Genomes Project Consortium, A global reference for human genetic variation. Nature 526, 68–74 (2015). doi: 10.1038/nature15393; pmid: 26432245

60. A. De Marino et al., A comparative analysis of current phasing and imputation software. PLoS One. 17, e0260177 (2022). doi: 10.1371/journal.pone.0260177; pmid: 36260643

61. E. Gilbert, A. Shanmugam, G. L. Cavalleri, Revealing the recent demographic history of Europe via haplotype sharing in the UK Biobank. Proc. Natl. Acad. Sci. U. S. A. 119, e2119281119 (2022). doi: 10.1073/pnas.2119281119; pmid: 35696575

62. S. McCarthy et al., A reference panel of 64,976 haplotypes for genotype imputation. Nat. Genet. 48, 1279–1283 (2016). doi: 10.1038/ng.3643; pmid: 27548312

63. J. Huang et al., Improved imputation of low-frequency and rare variants using the UK10K haplotype reference panel. Nat. Commun. 6, 8111 (2015). doi: 10.1038/ncomms9111; pmid: 26368830

64. C. Bycroft et al., The UK Biobank resource with deep phenotyping and genomic data. Nature 562, 203–209 (2018). doi: 10.1038/s41586-018-0579-z; pmid: 30305743

65. J. T. Leek et al., Tackling the widespread and critical impact of batch effects in high-throughput data. Nat. Rev. Genet. 11, 733–739 (2010). doi: 10.1038/nrg2825; pmid: 20838408

66. C. C. Laurie et al., Quality control and quality assurance in genotypic data for genome-wide association studies. Genet. Epidemiol. 34, 591–602 (2010). doi: 10.1002/gepi.20516; pmid: 20718045

67. A. Diaz-Papkovich, L. Anderson-Trocmé, C. Ben-Eghan, S. Gravel, UMAP reveals cryptic population structure and phenotype heterogeneity in large genomic cohorts. PLoS Genet. 15, e1008432 (2019). doi: 10.1371/journal.pgen.1008432; pmid: 31675358

68. T. Akiba, S. Sano, T. Yanase, T. Ohta, M. Koyama. Optuna: A next-generation hyperparameter optimization framework. Proceedings of the 25th ACM SIGKDD International Conference on Knowledge Discovery & Data Mining; 2623–2631 (2019).

69. J. Demšar et al., Orange: Data Mining Toolbox in Python. J. Mach. Learn. Res. 14, 2349−2353 (2013). https://jmlr.org/papers/v14/demsar13a.html

70. J. Kelleher, A. M. Etheridge, G. McVean, Efficient coalescent simulation and genealogical analysis for large sample sizes. PLoS Comput. Biol. 12, e1004842 (2016). doi: 10.1371/journal.pcbi.1004842; pmid: 27145223

71. F. Pedregosa et al., Scikit-learn: Machine Learning in Python. J. Mach. Learn. Res. 12, 2825−2830 (2011). https://jmlr.csail.mit.edu/papers/v12/pedregosa11a.html

72. S. Carmi et al., Sequencing an Ashkenazi reference panel supports population-targeted personal genomics and illuminates Jewish and European origins. Nat. Commun. 5, 4835 (2014). doi: 10.1038/ncomms5835; pmid: 25203624

73. J. Kim et al., KoVariome: Korean National Standard Reference Variome database of whole genomes with comprehensive SNV, indel, CNV, and SV analyses. Sci. Rep. 8, 5677 (2018). doi: 10.1038/s41598-018-23837-x; pmid: 29618732

74. E. Lowy-Gallego et al., Variant calling on the GRCh38 assembly with the data from phase three of the 1000 Genomes Project. Wellcome Open Res. 4, 50 (2019). doi: 10.12688/wellcomeopenres.15126.2; pmid: 32175479

75. S. Mallick et al., The Simons Genome Diversity Project: 300 genomes from 142 diverse populations. Nature 538, 201–206 (2016). doi: 10.1038/nature18964; pmid: 27654912

76. M. A. Almarri et al., The genomic history of the Middle East. Cell 184, 4612–4625.e14 (2021). doi: 10.1016/j.cell.2021.07.013; pmid: 34352227

77. Malaria Genomic Epidemiology Network, Insights into malaria susceptibility using genome-wide data on 17,000 individuals from Africa, Asia and Oceania. Nat. Commun. 10, 5732 (2019). doi: 10.1038/s41467-019-13480-z; pmid: 31844061

78. W. Zhang et al., Whole genome sequencing of 35 individuals provides insights into the genetic architecture of Korean population. BMC Bioinformatics. 15 Suppl 11(Suppl 11):S6 (2014). doi: 10.1186/1471-2105-15-S11-S6; pmid: 25350283

79. C. C. Wang et al., Genomic insights into the formation of human populations in East Asia. Nature 591, 413–419 (2021). doi: 10.1038/s41586-021-03336-2; pmid: 33618348

80. C. Jeong et al., The genetic history of admixture across inner Eurasia. Nat. Ecol. Evol. 3, 966–976 (2019). doi: 10.1038/s41559-019-0878-2; pmid: 31036896

81. S. A. Biagini et al., People from Ibiza: an unexpected isolate in the Western Mediterranean. Eur. J. Hum. Genet. 27, 941–951 (2019). doi: 10.1038/s41431-019-0361-1; pmid: 30765884

82. D. N. Vyas, A. Al-Meeri, C. J. Mulligan, Testing support for the northern and southern dispersal routes out of Africa: an analysis of Levantine and southern Arabian populations. Am. J. Phys. Anthropol. 164, 736–749 (2017). doi: 10.1002/ajpa.23312; pmid: 28913852

83. P. Skoglund et al., Reconstructing Prehistoric African Population Structure. Cell 171, 59–71.e21 (2017) doi: 10.1016/j.cell.2017.08.049; pmid: 28938123

84. P. Skoglund et al., Genomic insights into the peopling of the Southwest Pacific. Nature 538, 510–513 (2016). doi: 10.1038/nature19844; pmid: 27698418

85. I. Lazaridis et al., Genomic insights into the origin of farming in the ancient Near East. Nature 536, 419–424 (2016). doi: 10.1038/nature19310; pmid: 27459054

86. I. Lazaridis et al., Ancient human genomes suggest three ancestral populations for present-day Europeans. Nature 513, 409–413 (2014). doi: 10.1038/nature13673; pmid: 25230663

87. J. K. Pickrell et al., The genetic prehistory of southern Africa. Nat. Commun. 3, 1143 (2012). doi: 10.1038/ncomms2140; pmid: 23072811

88. P. Anagnostou et al., Berbers and Arabs: Tracing the genetic diversity and history of Southern Tunisia through genome wide analysis. Am. J. Phys. Anthropol. 173, 697–708 (2020). doi: 10.1002/ajpa.24139; pmid: 32936953

89. B. M. Henn et al., Genomic ancestry of North Africans supports back-to-Africa migrations. PLoS Genet. 8, e1002397 (2012). doi: 10.1371/journal.pgen.1002397; pmid: 22253600

90. L. R. Arauna et al., Recent Historical Migrations Have Shaped the Gene Pool of Arabs and Berbers in North Africa. Mol. Biol. Evol. 34, 318–329 (2017). doi:10.1093/molbev/msw218; pmid: 27744413

91. N. Hollfelder et al., Northeast African genomic variation shaped by the continuity of indigenous groups and Eurasian migrations. PLoS Genet. 13, e1006976 (2017). doi: 10.1371/journal.pgen.1006976; pmid: 28837655

92. Dobon B et al., The genetics of East African populations: a Nilo-Saharan component in the African genetic landscape. Sci Rep. 5, 9996 (2015). doi: 10.1038/srep09996; pmid: 26017457

93. D. M. Behar et al., No evidence from genome-wide data of a Khazar origin for the Ashkenazi Jews. Hum. Biol. 85, 859–900 (2013). doi: 10.3378/027.085.0604; pmid: 25079123

94. B. Yunusbayev et al., The Caucasus as an asymmetric semipermeable barrier to ancient human migrations. Mol. Biol. Evol. 29, 359–365 (2012). doi: 10.1093/molbev/msr221 pmid: 21917723

95. B. Yunusbayev et al., The genetic legacy of the expansion of Turkic-speaking nomads across Eurasia. PLoS Genet. 11, e1005068 (2015). doi: 10.1371/journal.pgen.1005068; pmid: 25898006

96. L. R. Botigué et al., Gene flow from North Africa contributes to differential human genetic diversity in southern Europe. Proc. Natl. Acad. Sci. U. S. A. 110, 11791–11796 (2013). doi: 10.1073/pnas.1306223110; pmid: 23733930

97. A. Flores-Bello et al., Genetic origins, singularity, and heterogeneity of Basques. Curr. Biol. 31, 2167–2177.e4 (2021). doi: 10.1016/j.cub.2021.03.010; pmid: 33770488

98. A. K. Pathak et al., The Genetic Ancestry of Modern Indus Valley Populations from Northwest India. Am. J. Hum. Genet. 103, 918–929 (2018). doi: 10.1016/j.ajhg.2018.10.022; pmid: 30526867

99. M. R. Nelson et al., The Population Reference Sample, POPRES: a resource for population, disease, and pharmacological genetics research. Am. J. Hum. Genet. 83, 347–358 (2008). doi: 10.1016/j.ajhg.2008.08.005; pmid: 18760391

100. P. Changmai et al., Indian genetic heritage in Southeast Asian populations. PLoS Genet. 18, e1010036 (2022). doi: 10.1371/journal.pgen.1010036; pmid: 35176016

101. K. Tätte et al., The genetic legacy of continental scale admixture in Indian Austroasiatic speakers. Sci. Rep. 9, 3818 (2019). doi: 10.1038/s41598-019-40399-8; pmid: 30846778

102. A. Mörseburg et al., Multi-layered population structure in Island Southeast Asians. Eur. J. Hum. Genet. 24, 1605–1611 (2016). doi: 10.1038/ejhg.2016.60; pmid: 27302840

103. D. Pierron et al., Strong selection during the last millennium for African ancestry in the admixed population of Madagascar. Nat. Commun. 9, 932 (2018). doi: 10.1038/s41467-018-03342-5; pmid: 29500350

104. G. Hudjashov et al., Complex Patterns of Admixture across the Indonesian Archipelago. Mol. Biol. Evol. 34, 2439–2452 (2017). doi: 10.1093/molbev/msx196; pmid: 28957506

105. A. Moreno-Estrada et al., Human genetics. The genetics of Mexico recapitulates Native American substructure and affects biomedical traits. Science 344, 1280–1285 (2014). doi: 10.1126/science.1251688; pmid: 24926019

106. M. Vicente et al., Population history and genetic adaptation of the Fulani nomads: inferences from genome-wide data and the lactase persistence trait. BMC Genomics 20, 915 (2019). doi: 10.1186/s12864-019-6296-7; pmid: 31791255

107. R. Laso-Jadart et al., The Genetic Legacy of the Indian Ocean Slave Trade: Recent Admixture and Post-admixture Selection in the Makranis of Pakistan. Am. J. Hum. Genet. 101, 977–984 (2017). doi: 10.1016/j.ajhg.2017.09.025; pmid: 29129317

