## Supplementary Figures and Tables for "Retracing Human Genetic Histories and Natural Selection Using Precise Local Ancestry Inference"

Figs. S1 to S64
Tables S1 to S3
References (*50 - 107*)

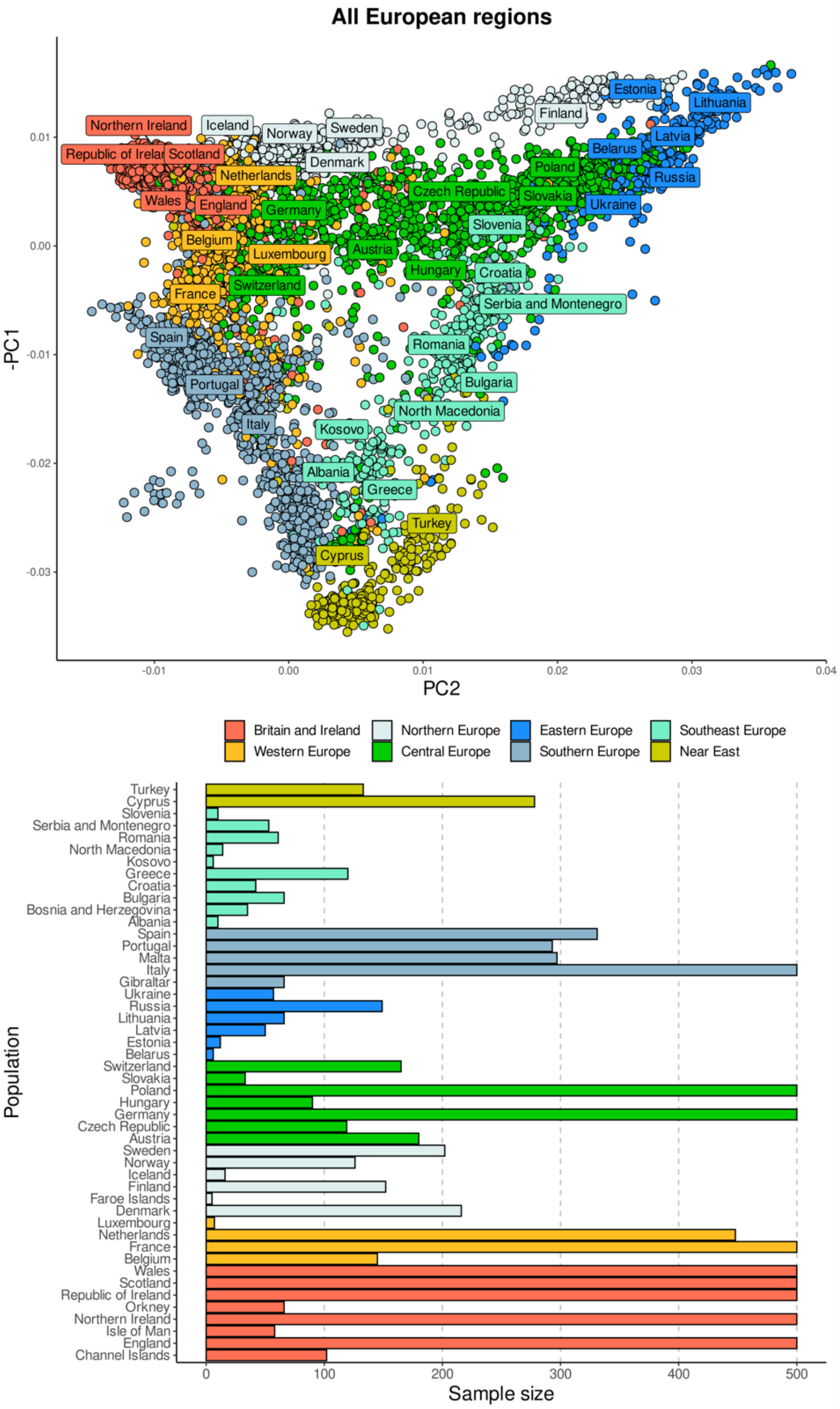


**Fig. S1. PCA of European UKBB participants used in the reference panel (A) and the number of samples per population (B)**. Countries and regions were randomly down-sampled to 500 individuals if their sample size was larger, in order to retain comparable numbers between populations for illustration purposes. Some country labels were not shown and rearranged around the sample centroids so they do not overlap. Note that this means that those populations capped at 500 have more participants from these regions.

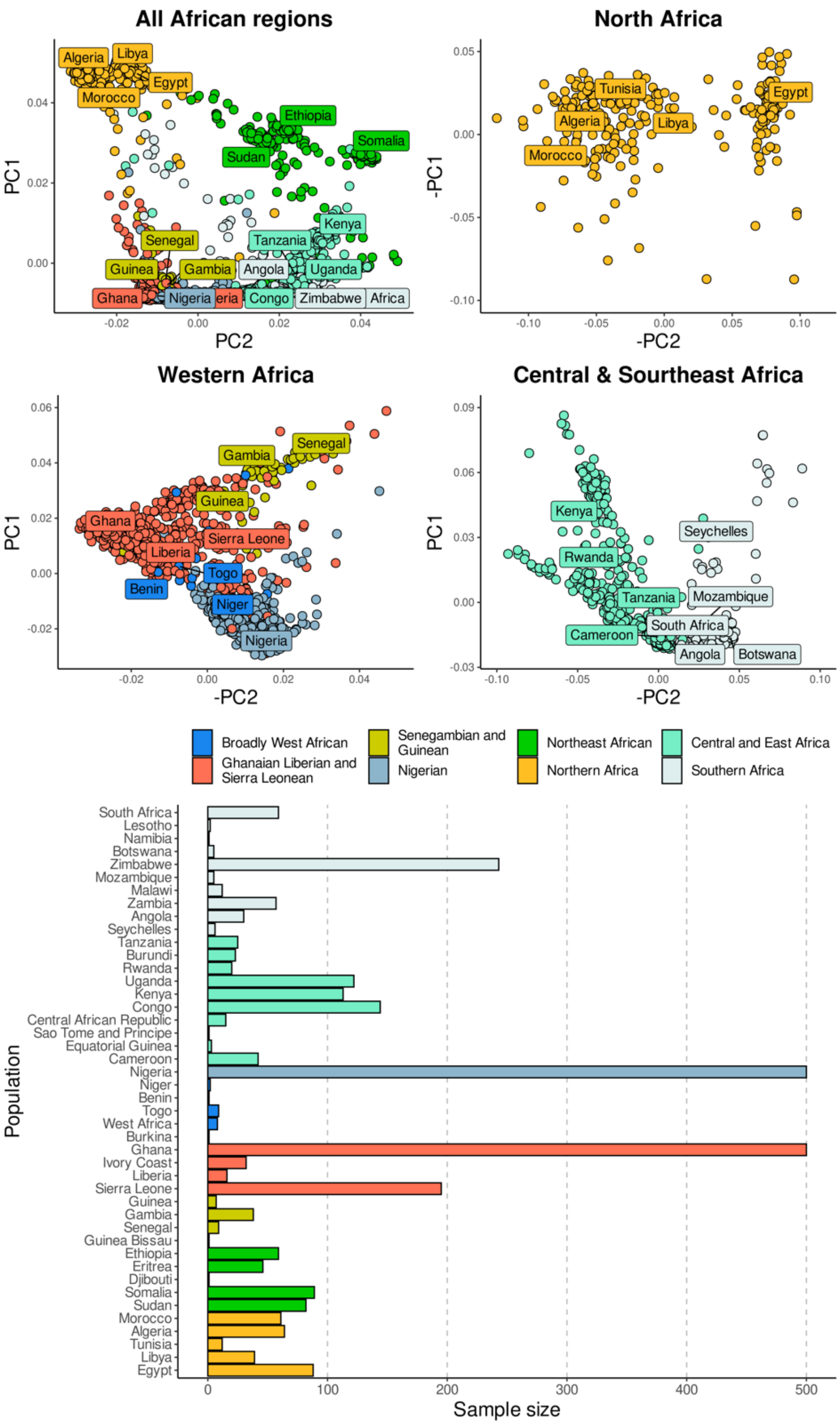


**Fig. S2. PCA of African UKBB participants used in the reference panel (A) and the number of samples per population (B).** Countries and regions were randomly down-sampled to 500 individuals if their sample size was larger, in order to retain comparable numbers between populations for illustration purposes. Some country labels were not shown and rearranged around the sample centroids so they do not overlap. Note that this means that those populations capped at 500 have more participants from these regions.

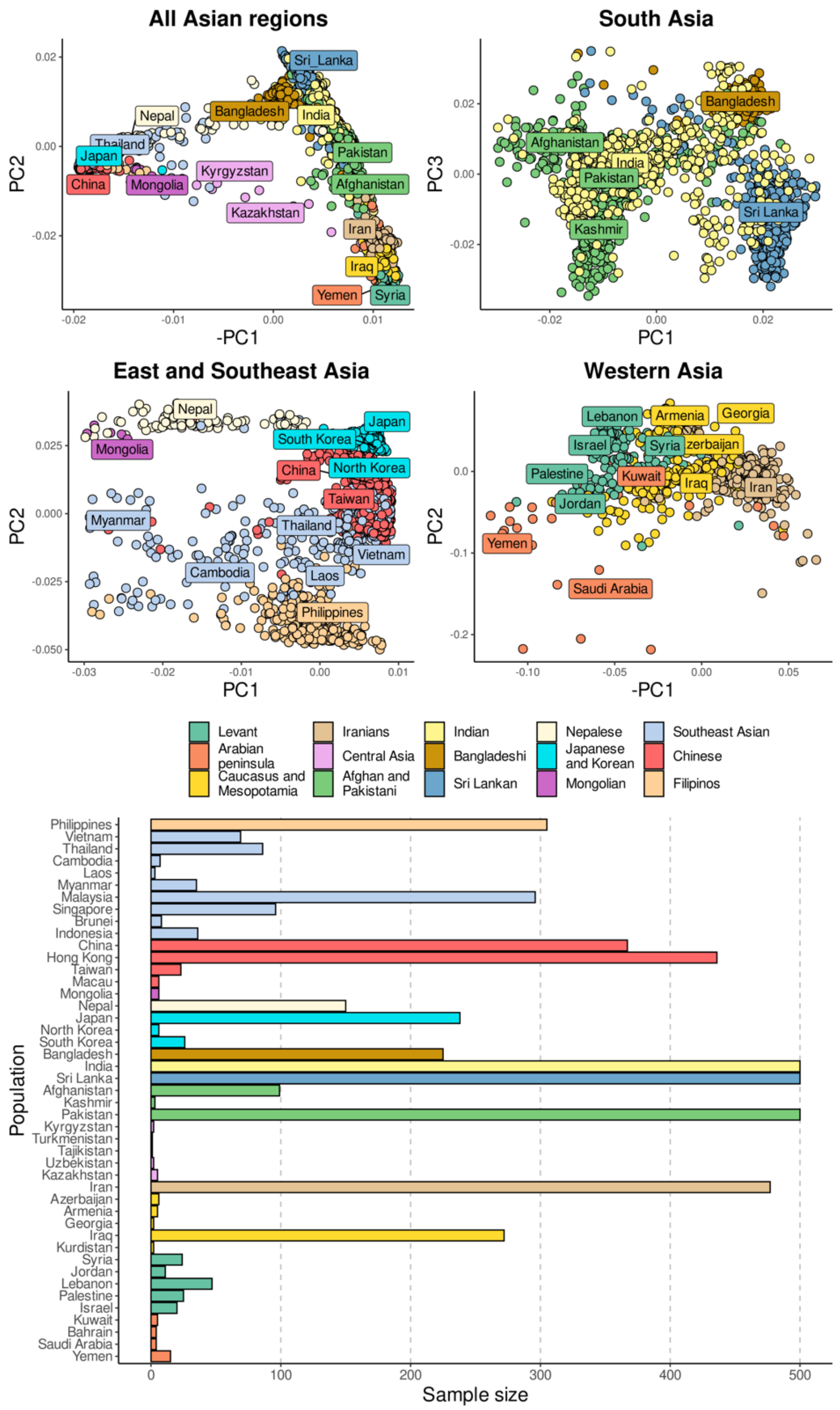


**Fig. S3. PCA of Asian UKBB participants used in the reference panel (A) and the number of samples per population (B).** Countries and regions were randomly down-sampled to 500 individuals if their sample size was larger, in order to retain comparable numbers between populations for illustration purposes. Some country labels were not shown and rearranged around the sample centroids so they do not overlap. Note that this means that those populations capped at 500 have more participants from these regions.

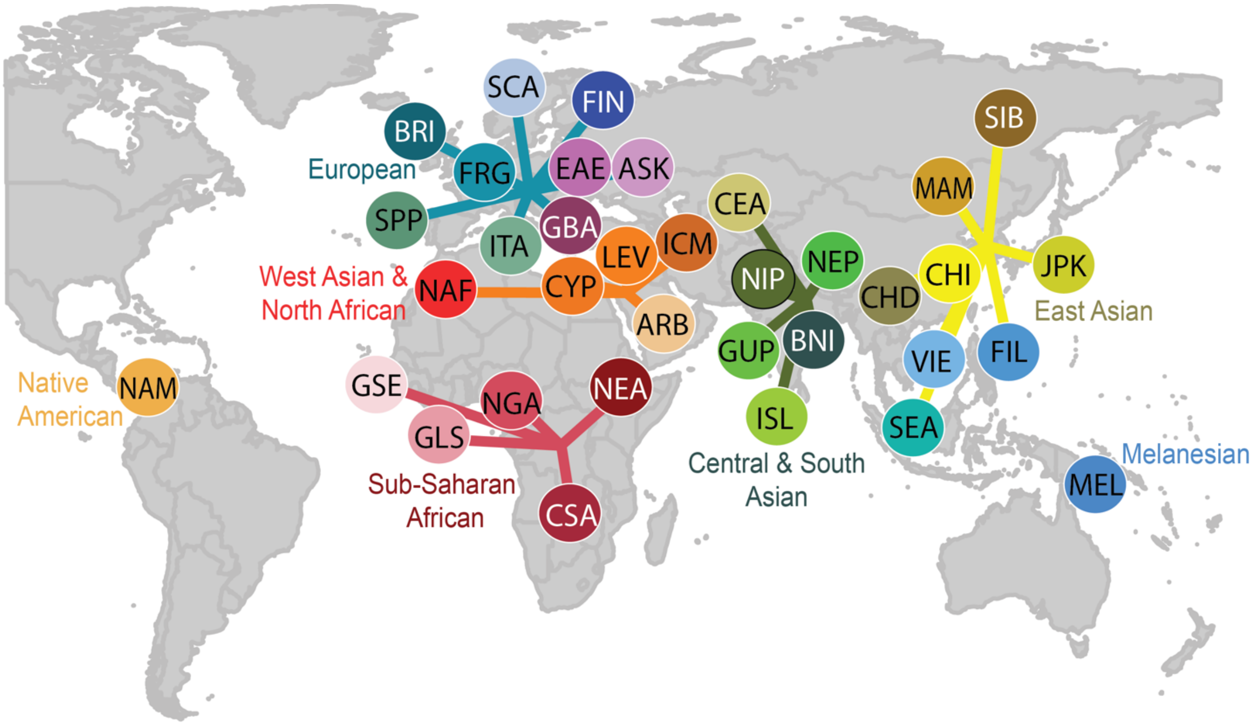


**Fig. S4. Map of 35 reference populations.** ARB = Arab, ASK = Ashkenazi Jewish, BRI = British & Irish, BNI = Bengali & East Indian, CEA = Central Asian, CHD = Chinese Dai, CHI = Han Chinese, CSA = Central, South & Southeast African, CYP = Cypriot, EAE = Eastern European, FIL = Filipino, FIN = Finish, FRG = French & German, GBA = Greek & Balkan, GLS = Ghanaian, Ivorian, Liberian & Sierra Leonean, GSE = Gambian & Senegalese, GUP = Gujarati Patels, ICM = Turkish, Iraqi, Iranian & Caucasian, ISL = Southern Indian & Sri Lankan, ITA = Italian, JPK = Japanese & Korean, LEV = Levantine, MAM = Manchurian & Mongolian, MEL = Melanesian & Aboriginal Australian, NAF = North African, NAM = Native American, NEA = Northeast African, NEP = Nepalese, NGA = Nigerian, NIP = North Indian & Pakistani, SCA = Scandinavian, SEA = Southeast Asian, SIB = Siberian, SPP = Spanish & Portuguese, VIE = Vietnamese.

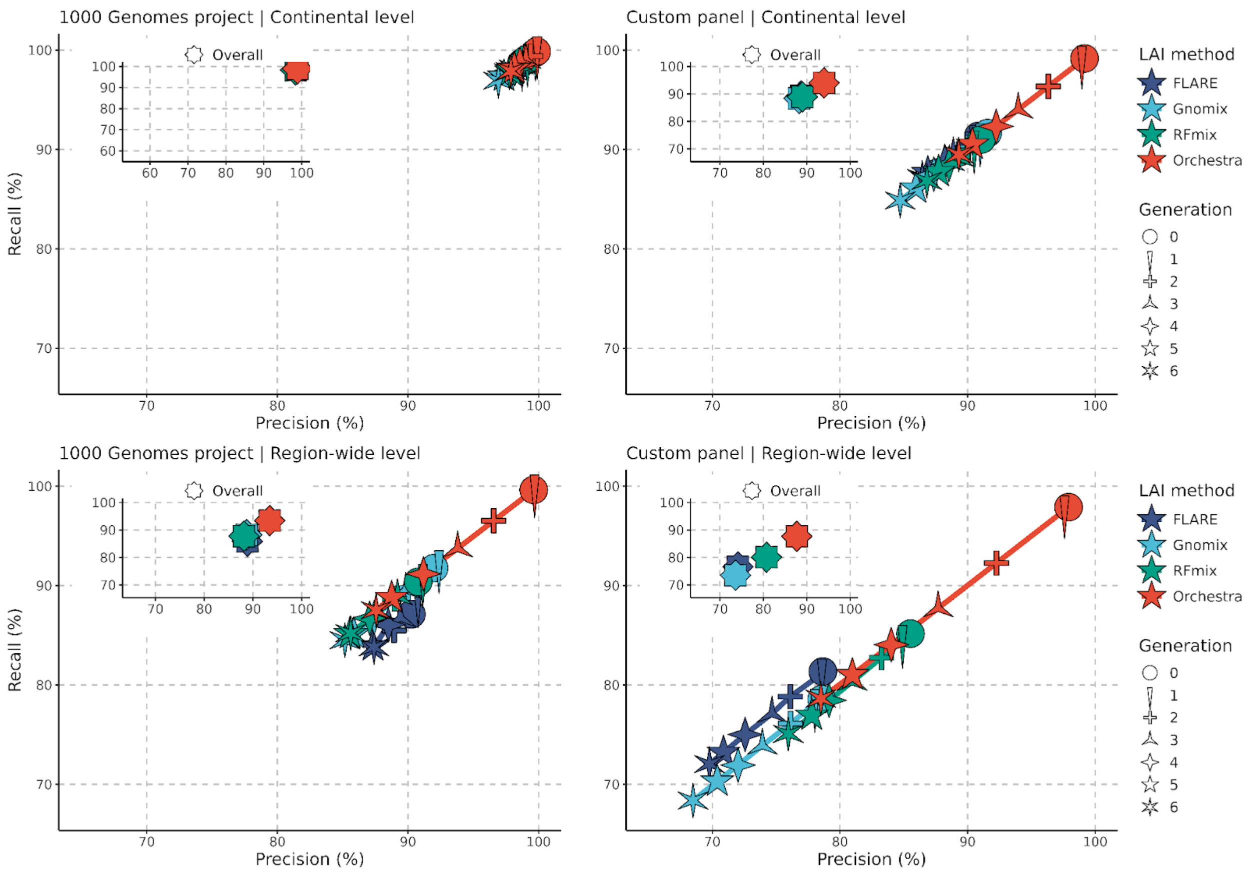


**Fig. S5. Performance of Orchestra and other LAI methods at continental and region-wide levels.** Recall and precision of Orchestra, RFmix, Gnomix and FLARE in ancestry deconvolution across 6 generations for 1KGP-16pops (left column) and custom-35pops (right column) reference panels. The generation is shown with star shapes and refers to the number of generations of simulated admixture (the more points the star has, the higher the generation). Performance at continental (upper row) and region-wide ( bottom row) level is shown. For the continental level, samples were grouped into 7 broad groups. (1) Sub-Saharan African 1KGP, (2) West Asian & Northern African, (3) European 1KGP, (4) Central & South Asian 1KGP, (5) East Asian 1KGP, (6) Melanesian and (7) Native American *. For the region-wide level, samples were grouped into 18 broad groups: (1) West African 1KGP, (2) Northern East African, (3) Central, South & Southeast African 1KGP, (4) North African, (5) Arab & Levantine, (6) Northern West Asian, (7) Northwest European 1KGP, (8) East European, (9) Southern European 1KGP, (10) Ashkenazi Jewish, (11) Central Asian, (12) Northern South Asian 1KGP, (13) Southern South Asian 1KGP, (14) Chinese & Southeast Asian 1KGP, (15) Japanese & Korean 1KGP, (16) Northern Asian, (17) Native American * and (18) Melanesian. 1KGP = populations present in the 1000 genomes project. Sub-regional or population level with 35 populations is shown in Figure 1. * Artificially created from 1KGP. Results were calculated using from chr17 to chr22.

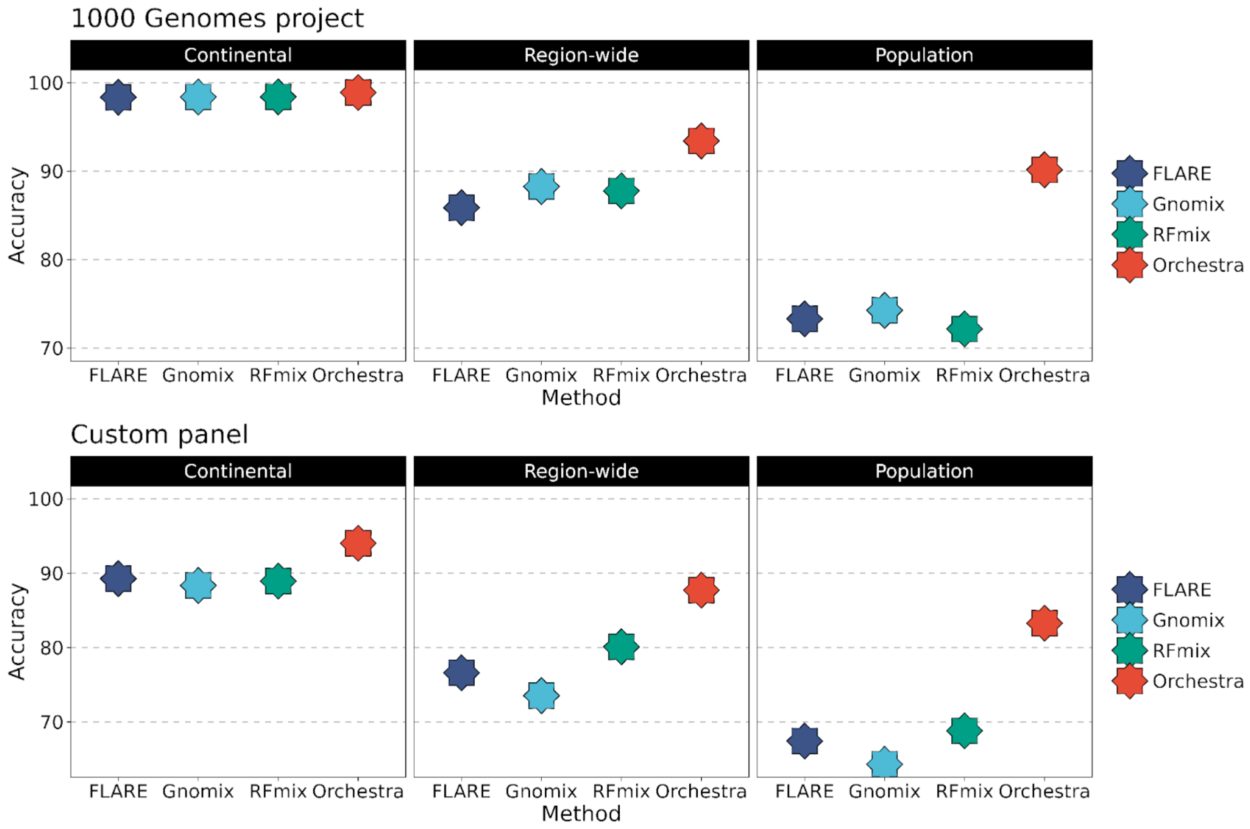


**Fig. S6. Performance across continental, region-wide and population level for all methods tested.** Accuracy (%) per panel for the 1KGP dataset and the larger custom dataset with 35 populations, across continental, region-wide and population level for all methods tested. For the continental level, samples were grouped into 7 broad groups. (1) Sub-Saharan African 1KGP, (2) West Asian & Northern African, (3) European 1KGP, (4) Central & South Asian 1KGP, (5) East Asian 1KGP, (6) Melanesian and (7) Native American *. For the region-wide level, samples were grouped into 18 broad groups: (1) West African 1KGP, (2) Northern East African, (3) Central, South & Southeast African 1KGP, (4) North African, (5) Arab & Levantine, (6) Northern West Asian, (7) Northwest European 1KGP, (8) East European, (9) Southern European 1KGP, (10) Ashkenazi Jewish, (11) Central Asian, (12) Northern South Asian 1KGP, (13) Southern South Asian 1KGP, (14) Chinese & Southeast Asian 1KGP, (15) Japanese & Korean 1KGP, (16) Northern Asian, (17) Native American * and (18) Melanesian. 1KGP = populations present in the 1000 genomes project. Sub-regional or population level with 35 populations is shown in Figure 1. * Artificially created from 1KGP. Results were calculated using from chr17 to chr22.

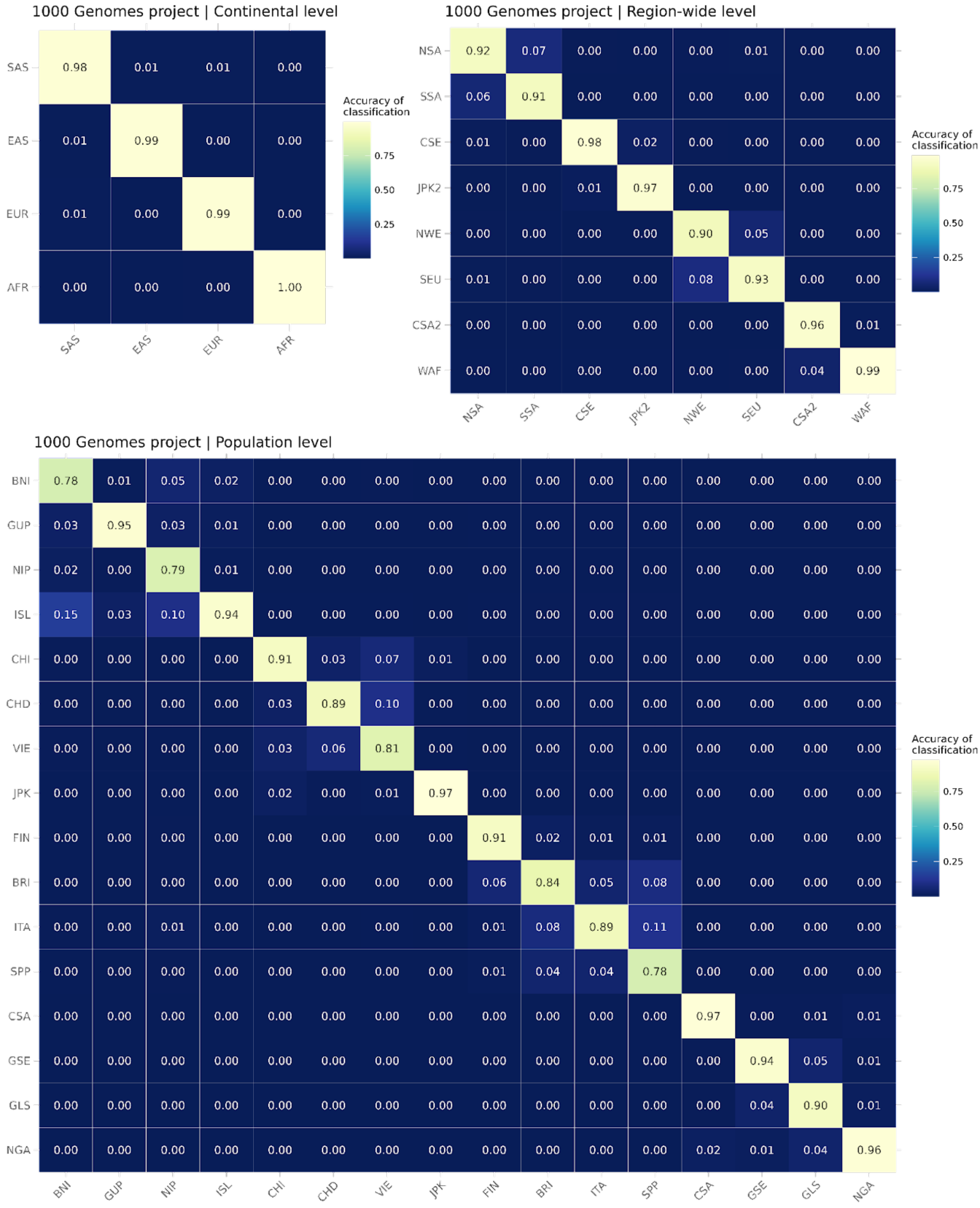


**Fig. S7. Confusion matrices of misclassifications obtained with Orchestra using the 1KGP-16pops panel.** Accuracy of ancestry classification and misclassifications at continental, region-wide and population levels.

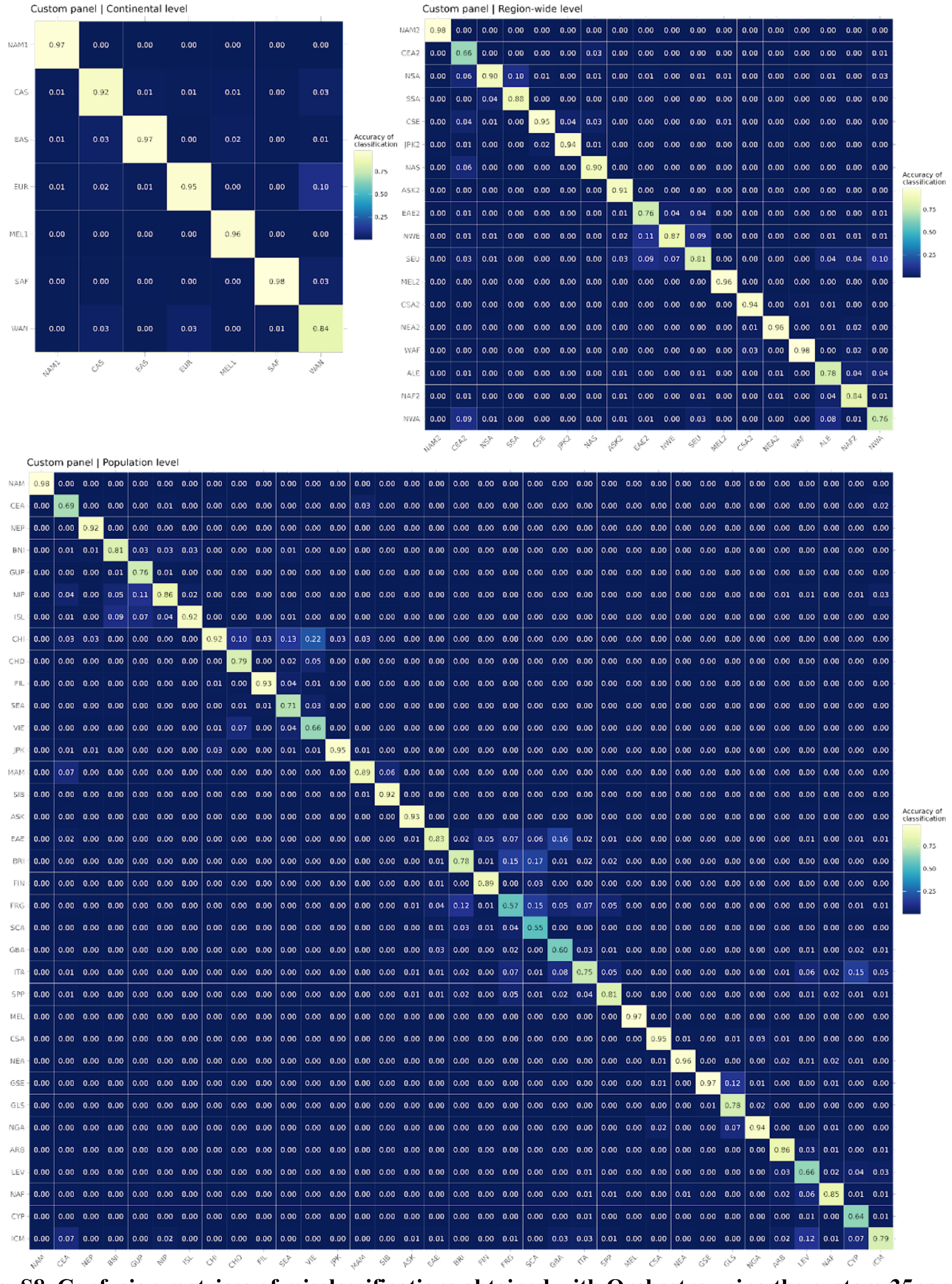


**Fig. S8. Confusion matrices of misclassifications obtained with Orchestra using the custom-35pops panel.** Accuracy of ancestry classification and misclassifications at continental, region-wide and population levels.

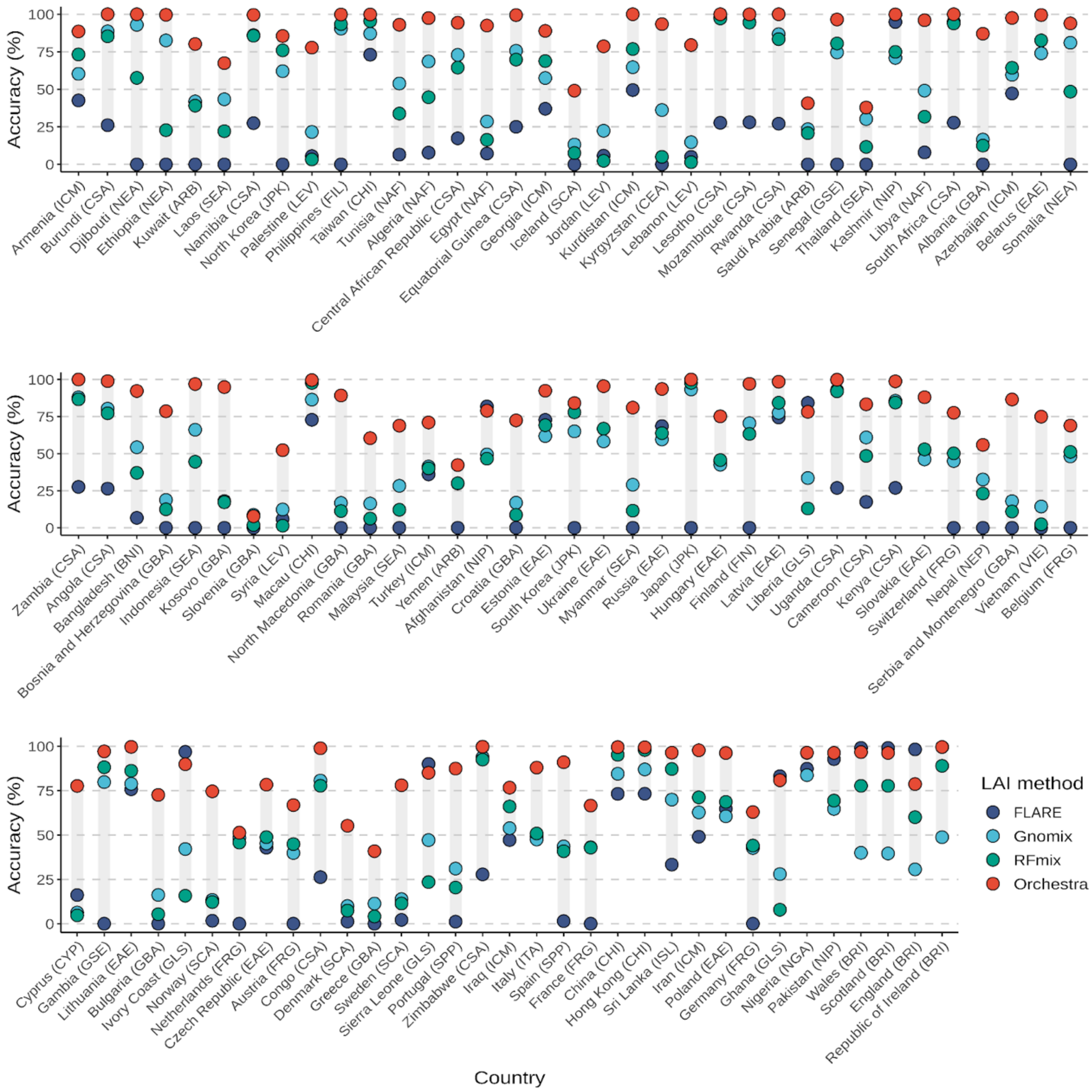


**Fig. S9. SuppFigure Benchmarking. ExternalUKBBpanel.** Accuracy (%) per country in the UKBB external panel with inferred single-origin individuals based on dimensionality reduction techniques. Countries are ordered by sample-size. The evaluated ancestry for each country is denoted in parentheses on the x-axis.

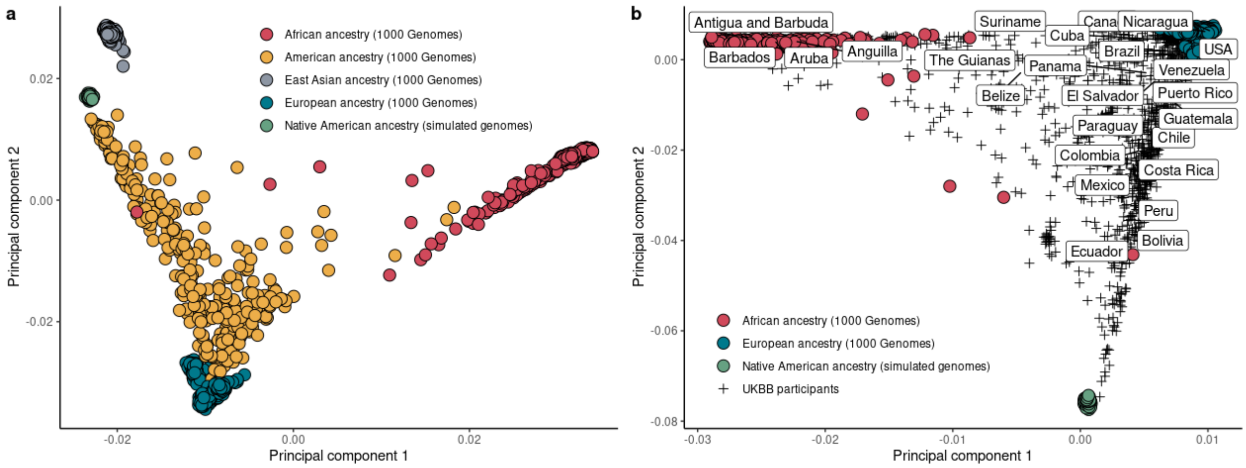


**Fig. S10. Population genetic structure of artificially-constructed Native American genomes.** (**A**) Principal component analysis (PCA) of generated pure Amerindian genomes compared to 1KGP samples as a reference. A gradient from European to American poles can be seen in those 1KGP admixed genomes that were sampled in the Americas. (**B**) UKBB participants that were born in American countries were included together with our set of pure Native American genomes and African and European individuals from 1KGP. Countries were indicated in the samples' centroid PC locations according to the UKBB participant labels. The same Europe-America gradient can be seen for admixed American UKBB participants.

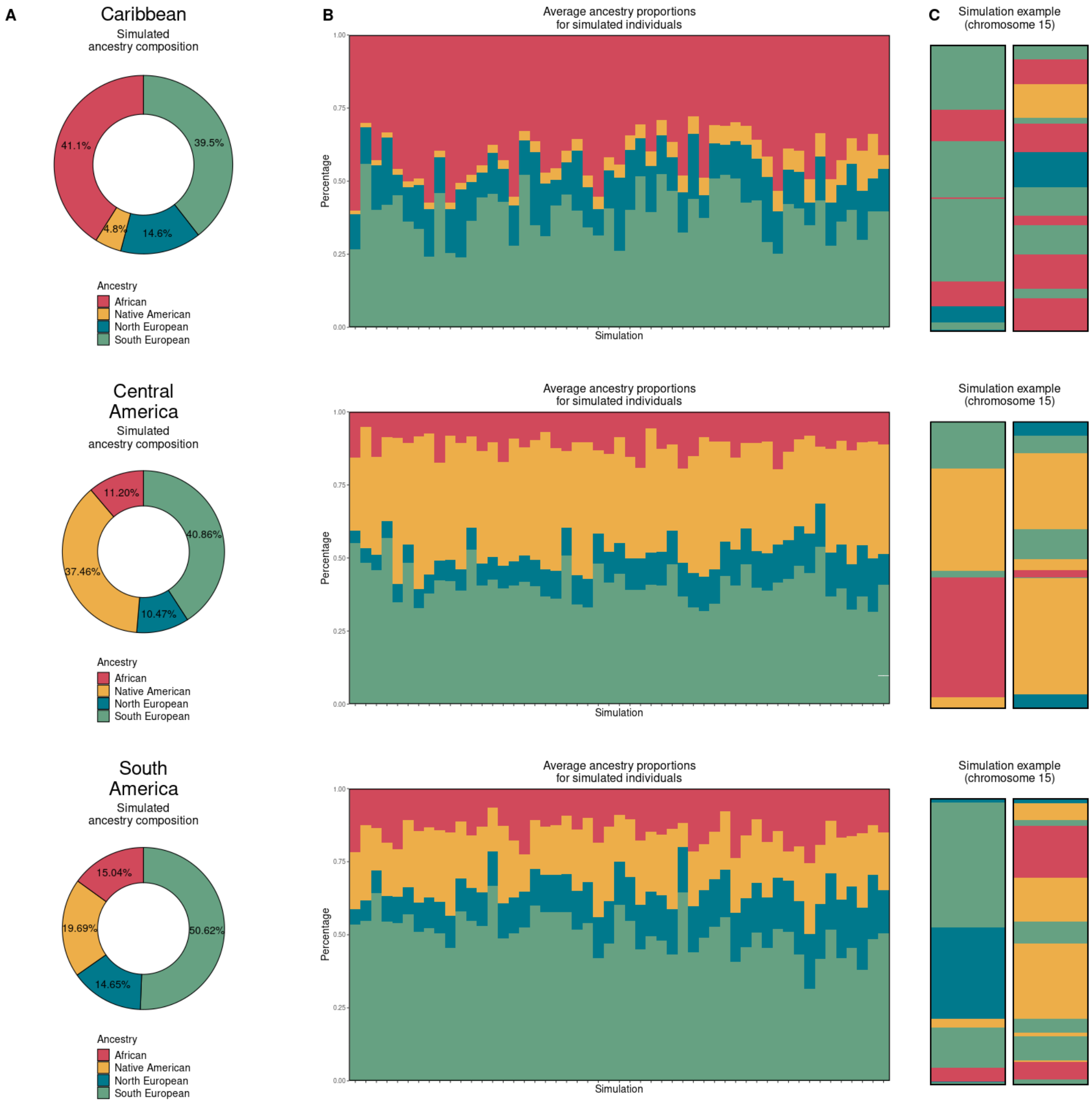


**Fig. S11. Simulated Latin American genomes for training and benchmarking purposes.** The simulated Latino genomes were designed to reflect the genetic patterns observed in contemporary Latinos from three major American regions: Caribbean (upper row), Central American (middle) and South America (lower row). We conducted intermixing for 12 generations using SLiM (A). Percentage of each ancestry per simulated sample in a subset of the simulations (B). Illustration of a simulated chromosome (15) showing the lengths of each segment derived from each ancestral origin (C).

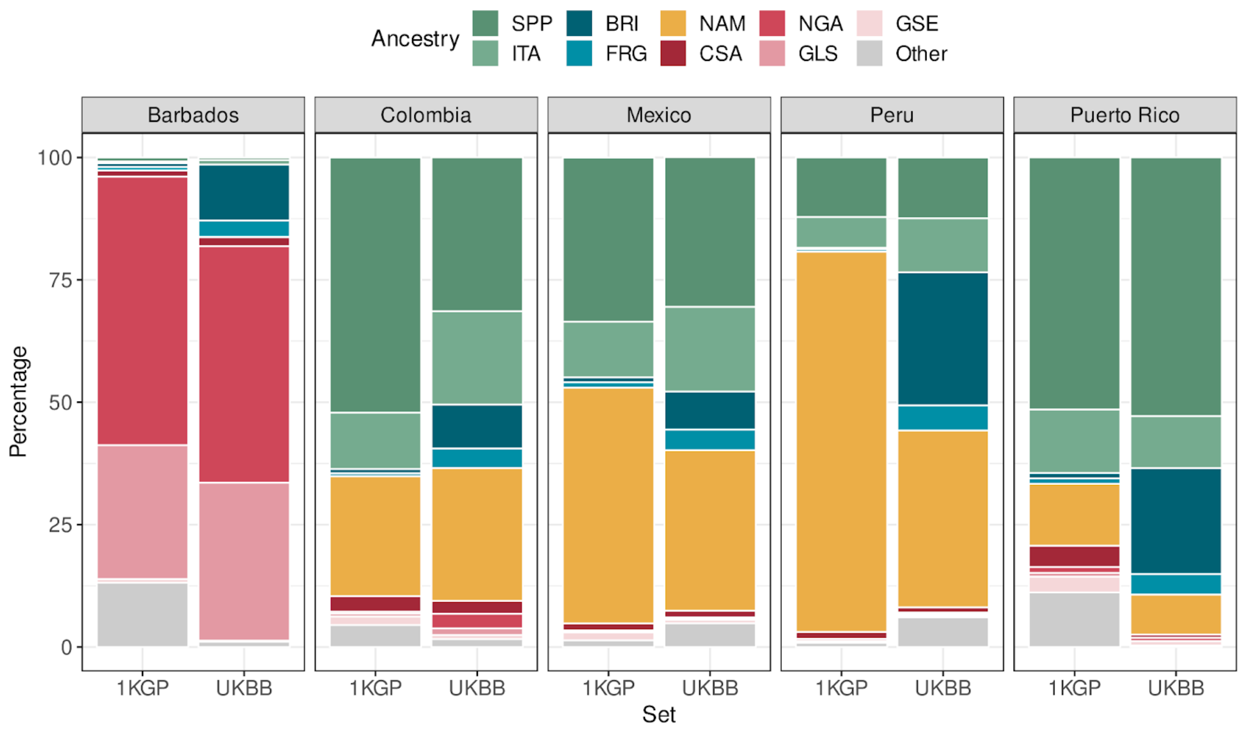


**Fig. S12. Comparison of ancestral composition between samples from 1KGP and UKBB.** UKBB participants that were born in the Americas show a higher proportion of British ancestry compared to 1KGP. BRI = British & Irish, CSA = Central, South & Southeast African, FRG = French & German, GLS = Ghanaian, Ivorian, Liberian & Sierra Leonean, GSE = Gambian & Senegalese, ITA = Italian, NAM = Native American, NGA = Nigerian, SPP = Spanish & Portuguese.

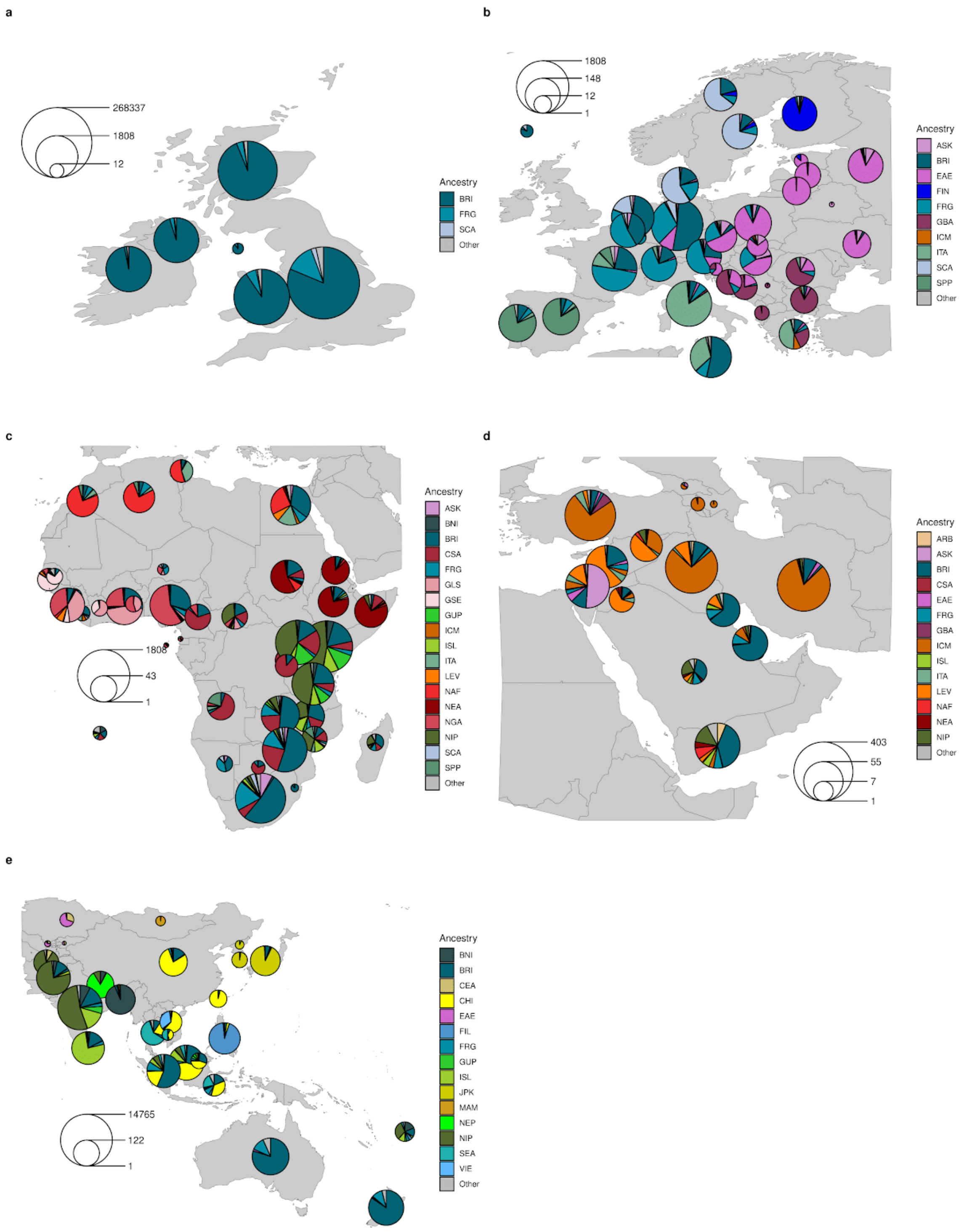


**Fig. S13. Inferred ancestry for UKBB samples mapped by country of birth.** Samples that were added to our reference panel were excluded. A bias towards excess British & Irish (BRI) ancestry is noted across all regions.

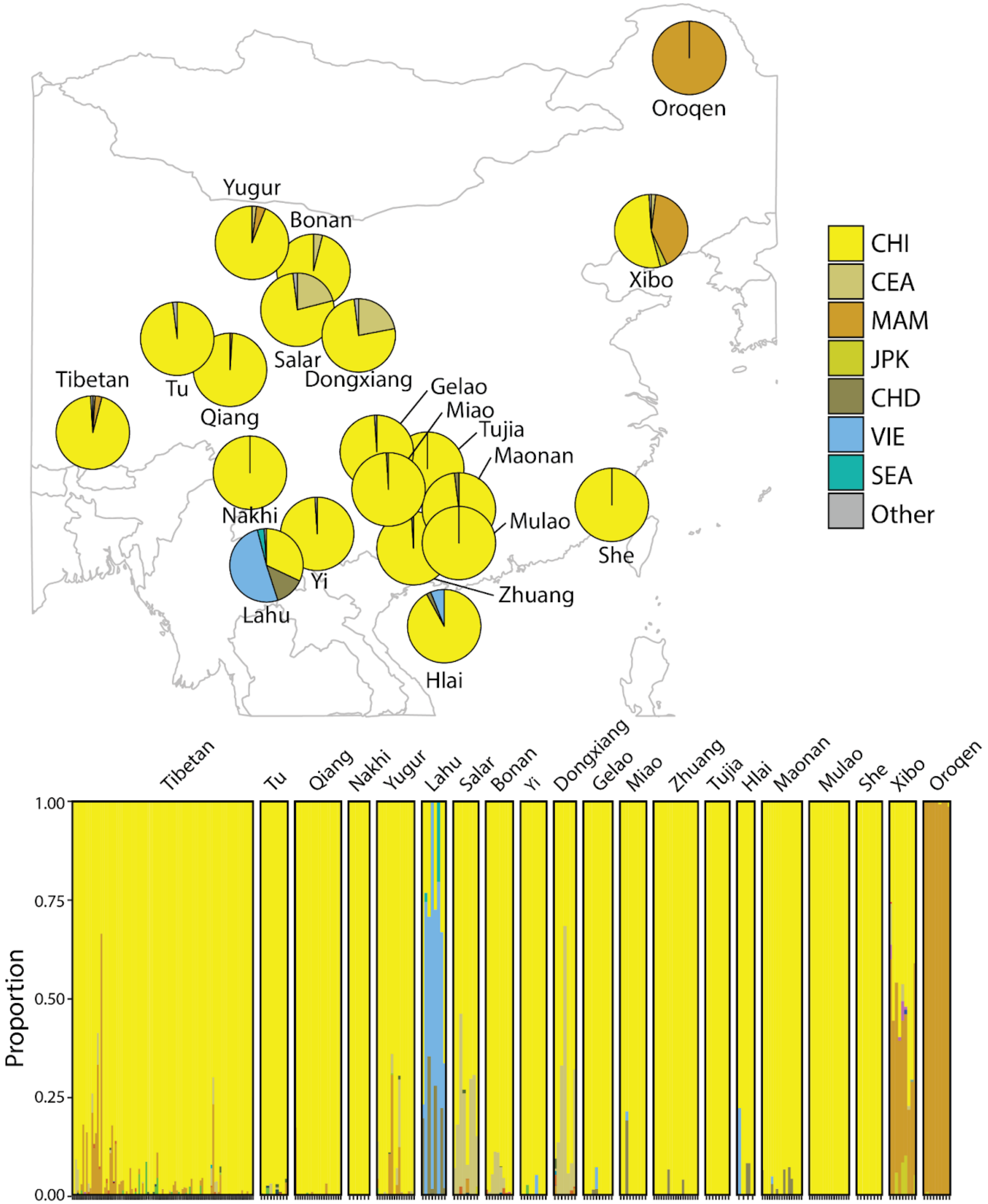


**Fig. S14.** Ancestry prediction by Orchestra for ethnic groups and minorities in China not included in our 35 reference populations.

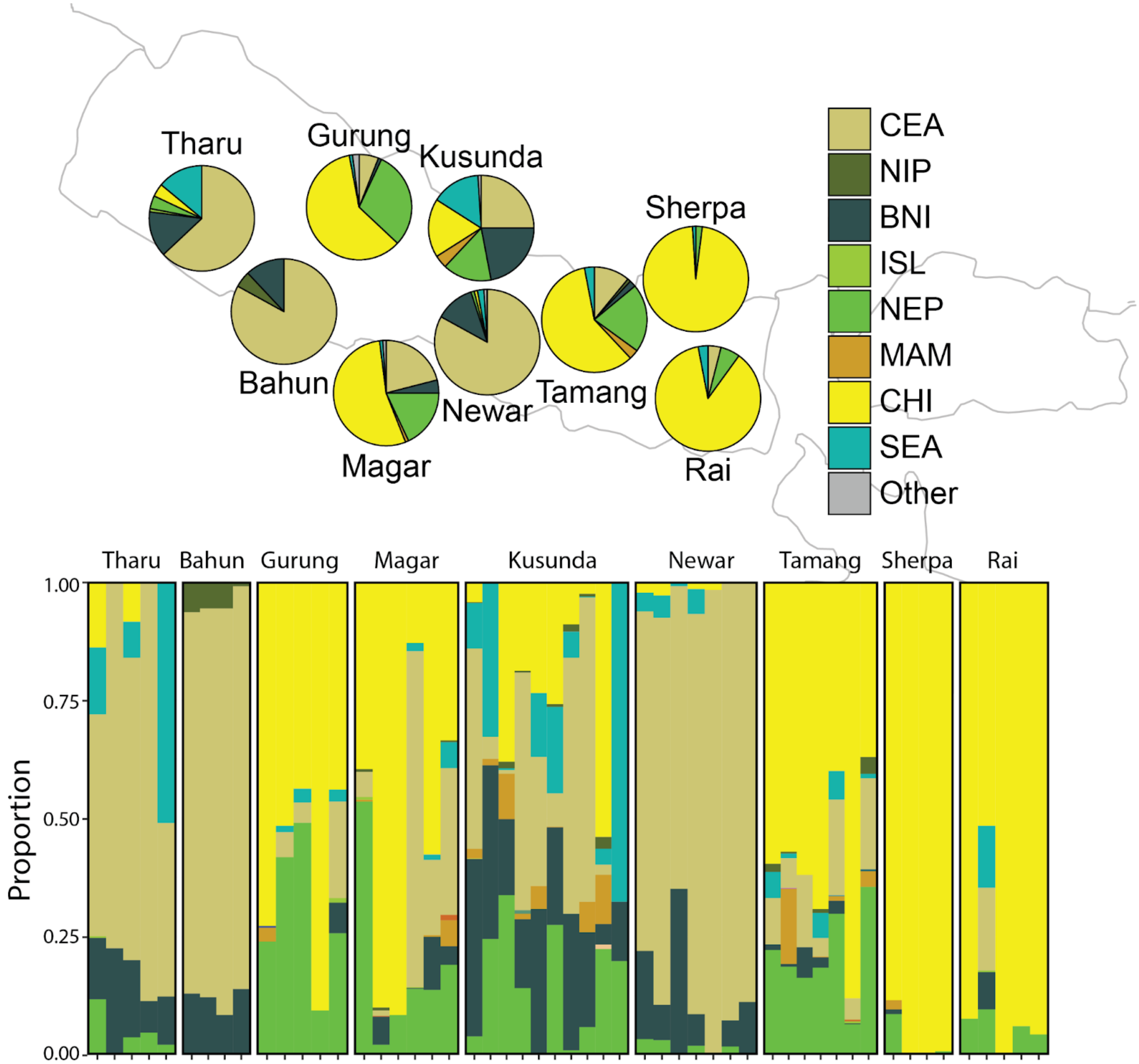


**Fig. S15.** Ancestry prediction by Orchestra for ethnic groups and minorities in Nepal not included in our 35 reference populations.

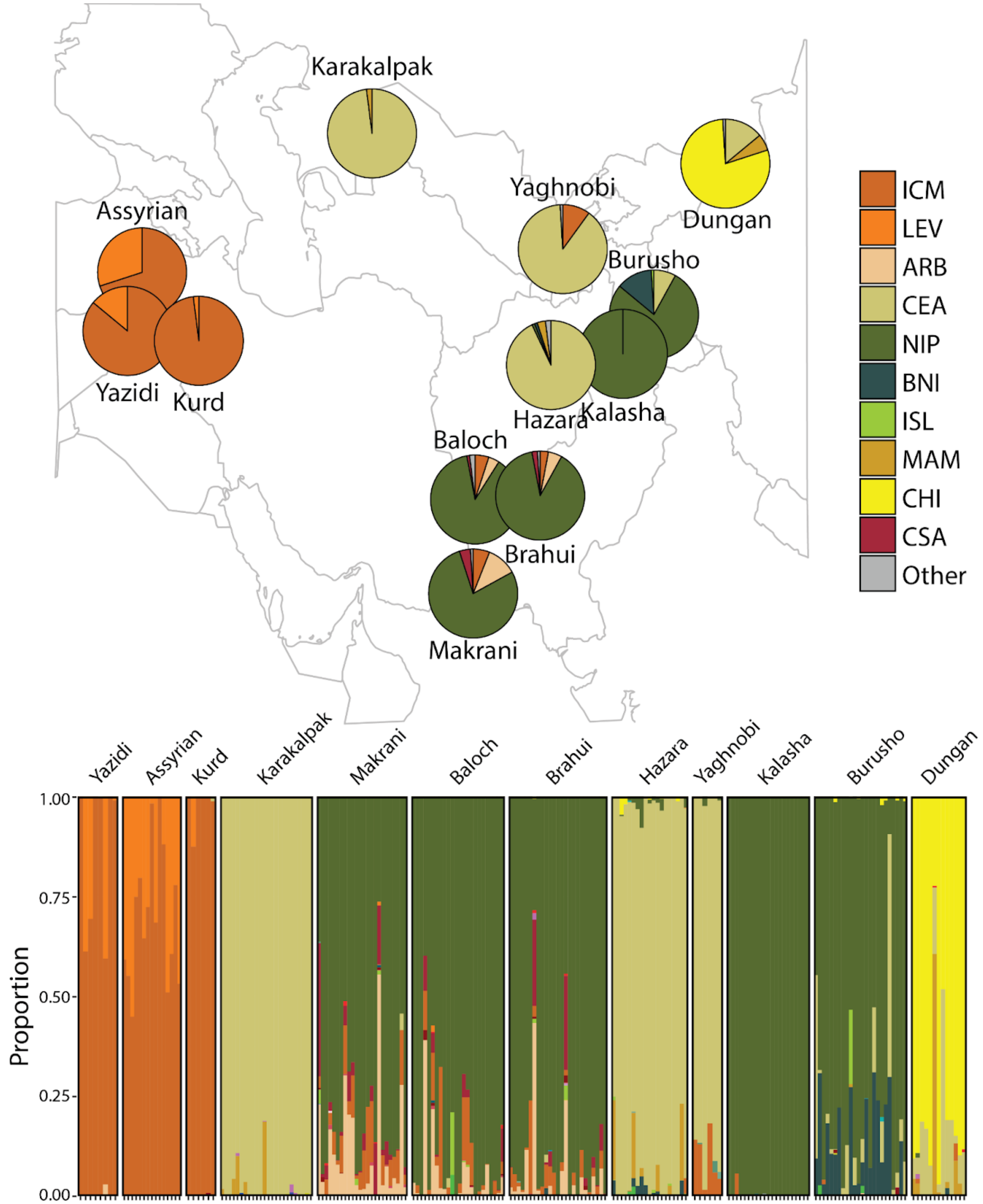


**Fig. S16.** Ancestry prediction by Orchestra for ethnic groups and minorities in Western and South Asia not included in our 35 reference populations.

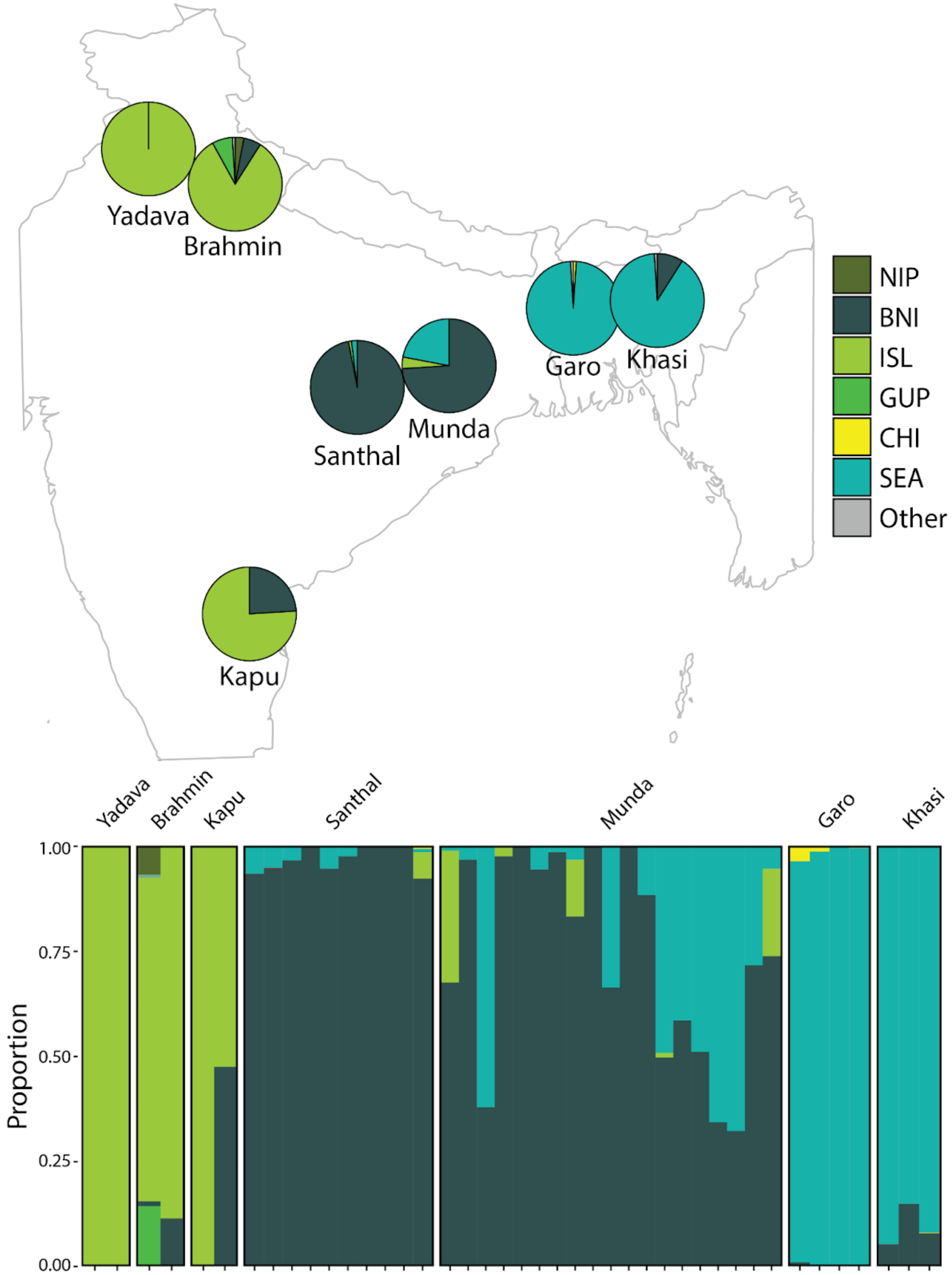


**Fig. S17.** Ancestry prediction by Orchestra for ethnic groups and minorities in South Asia not included in our 35 reference populations.

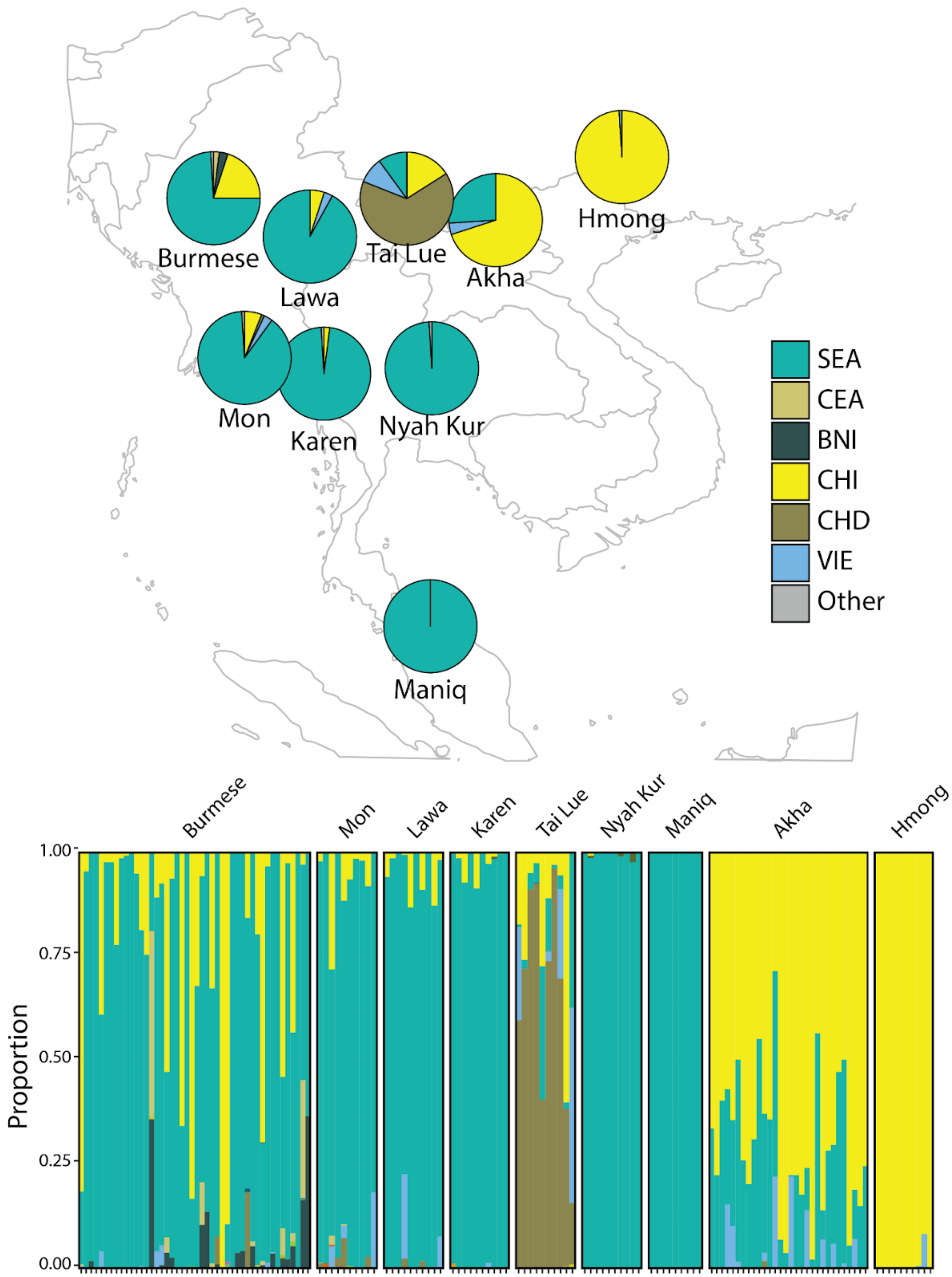


**Fig. S18.** Ancestry prediction by Orchestra for ethnic groups and minorities in Southeast Asia not included in our 35 reference populations.

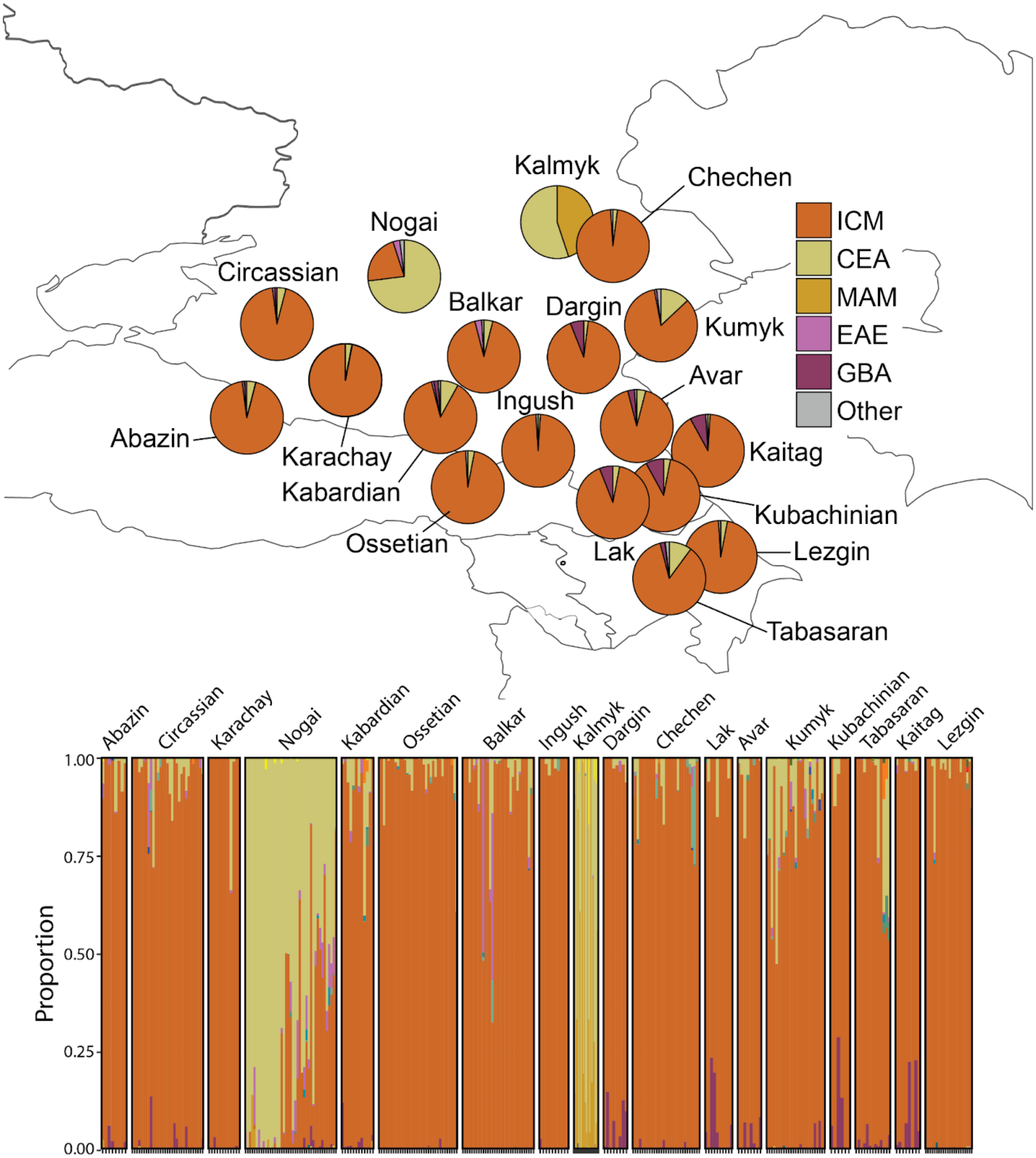


**Fig. S19.** Ancestry prediction by Orchestra for ethnic groups and minorities in the Caucassus not included in our 35 reference populations.

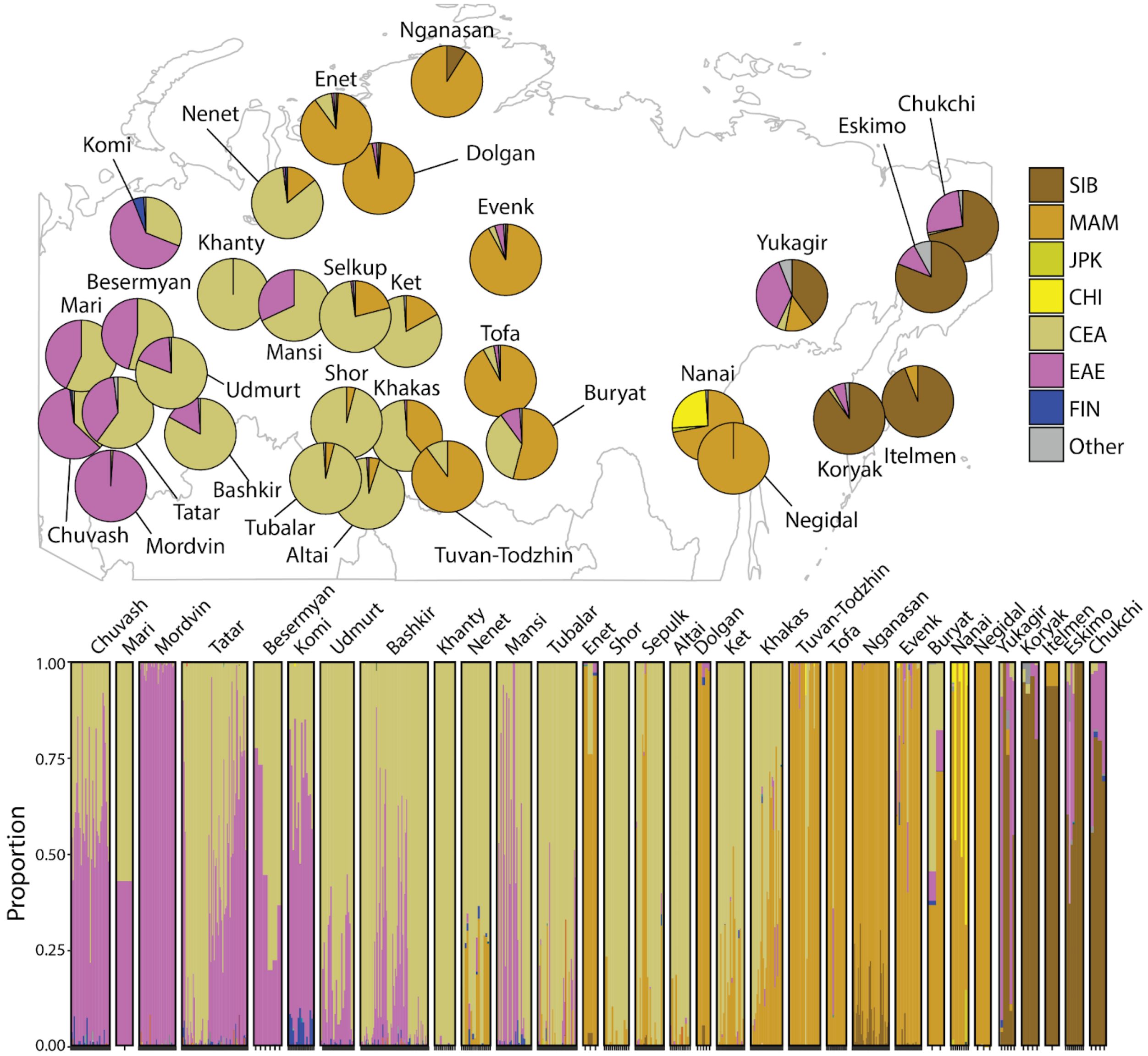


**Fig. S20.** Ancestry prediction by Orchestra for ethnic groups and minorities in Siberia not included in our 35 reference populations.

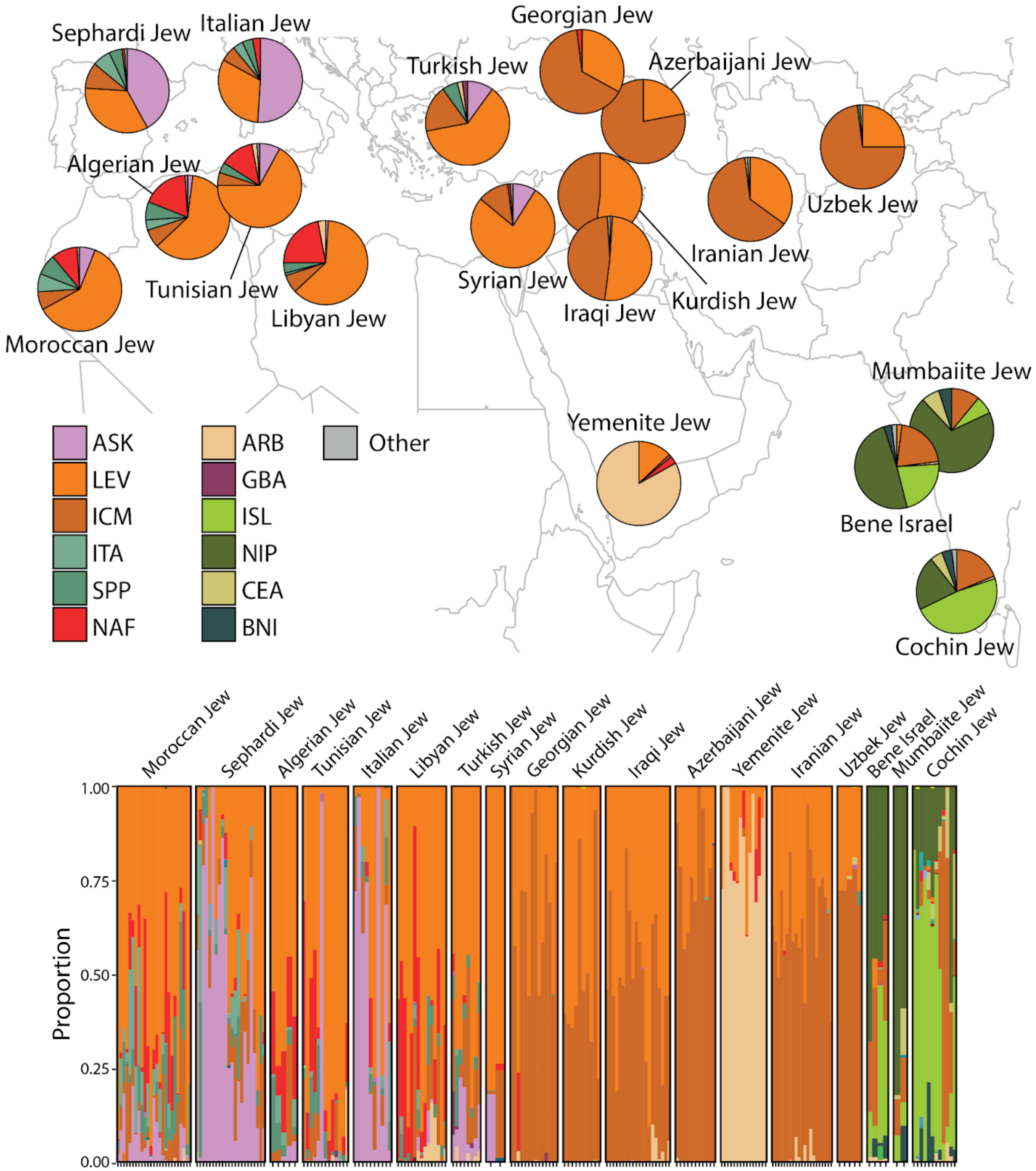


**Fig. S21.** Ancestry prediction by Orchestra for Jewish groups not included in our 35 reference populations.

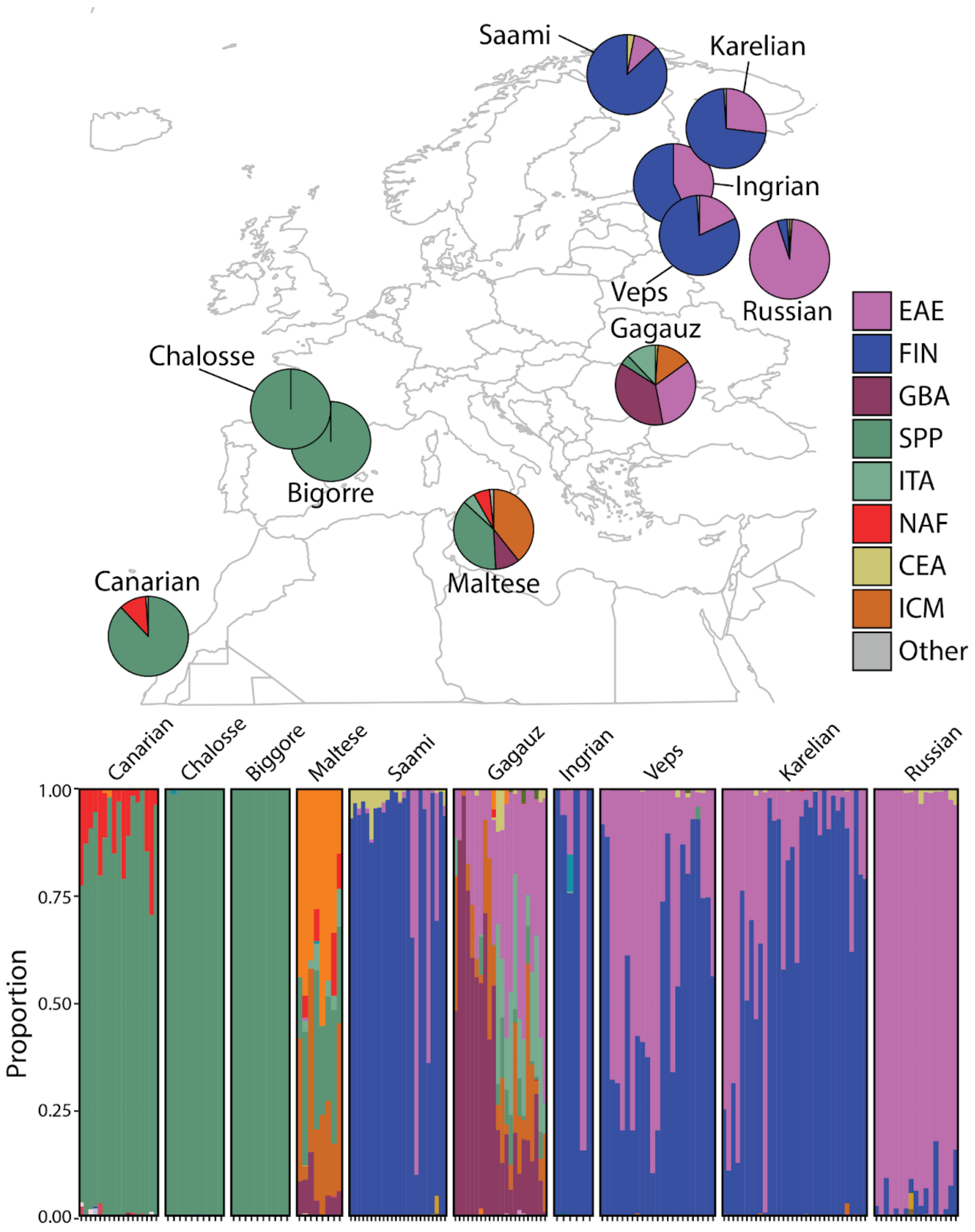


**Fig. S22.** Ancestry prediction by Orchestra for ethnic groups and minorities in Europe not included in our 35 reference populations.

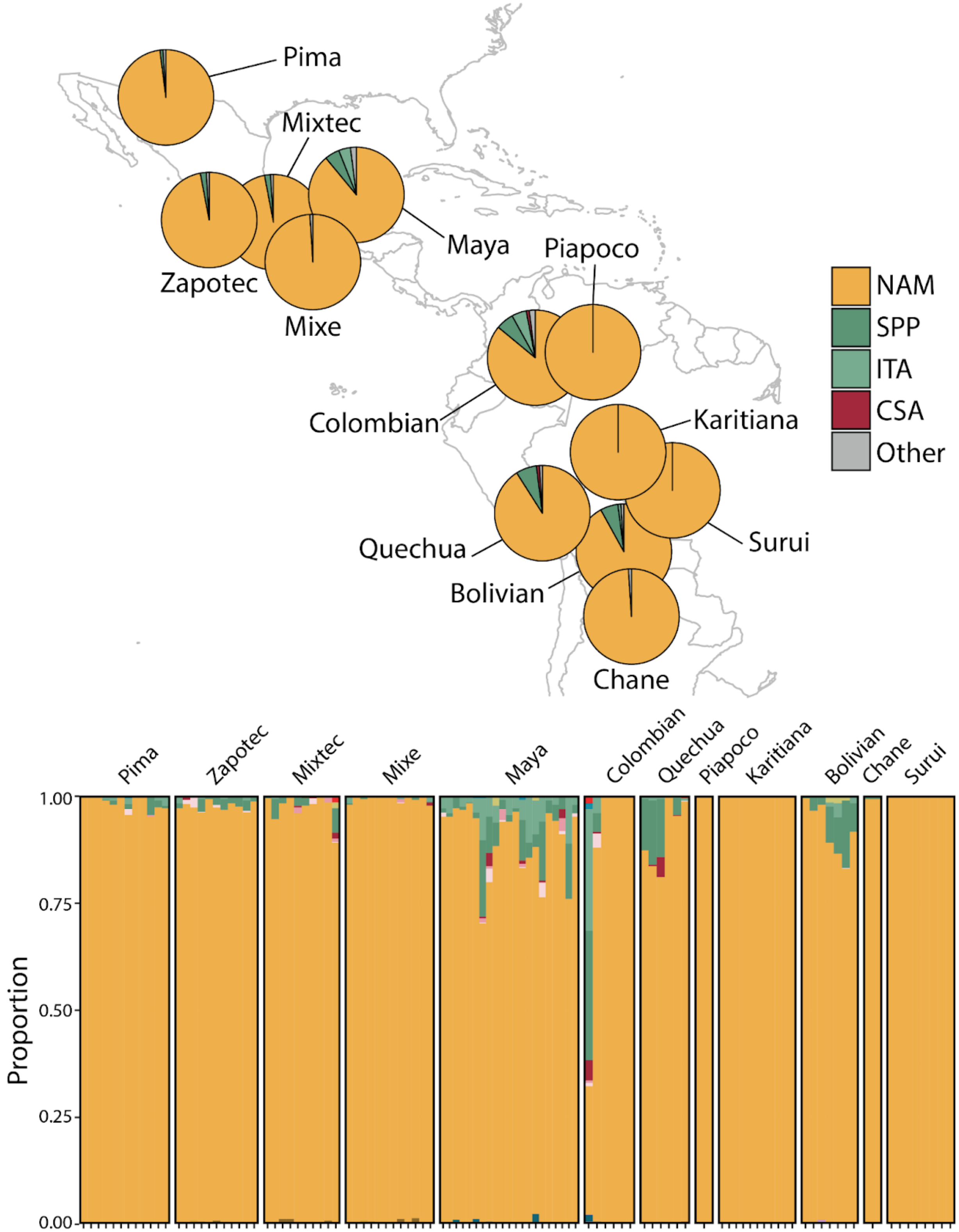


**Fig. S23.** Ancestry prediction by Orchestra for ethnic groups and minorities in South America.

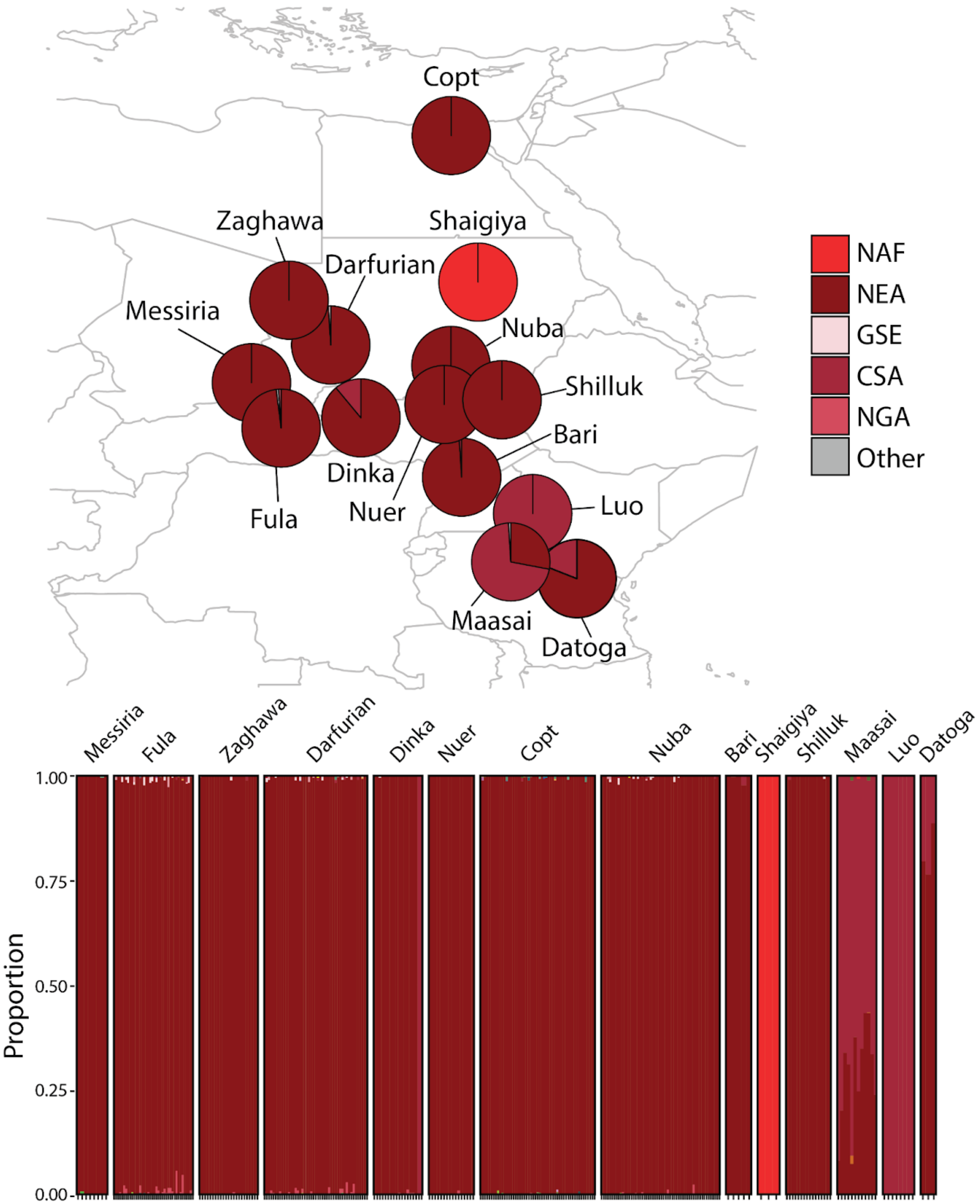


**Fig. S24.** Ancestry prediction by Orchestra for African ethnic groups and minorities not included in our 35 reference populations.

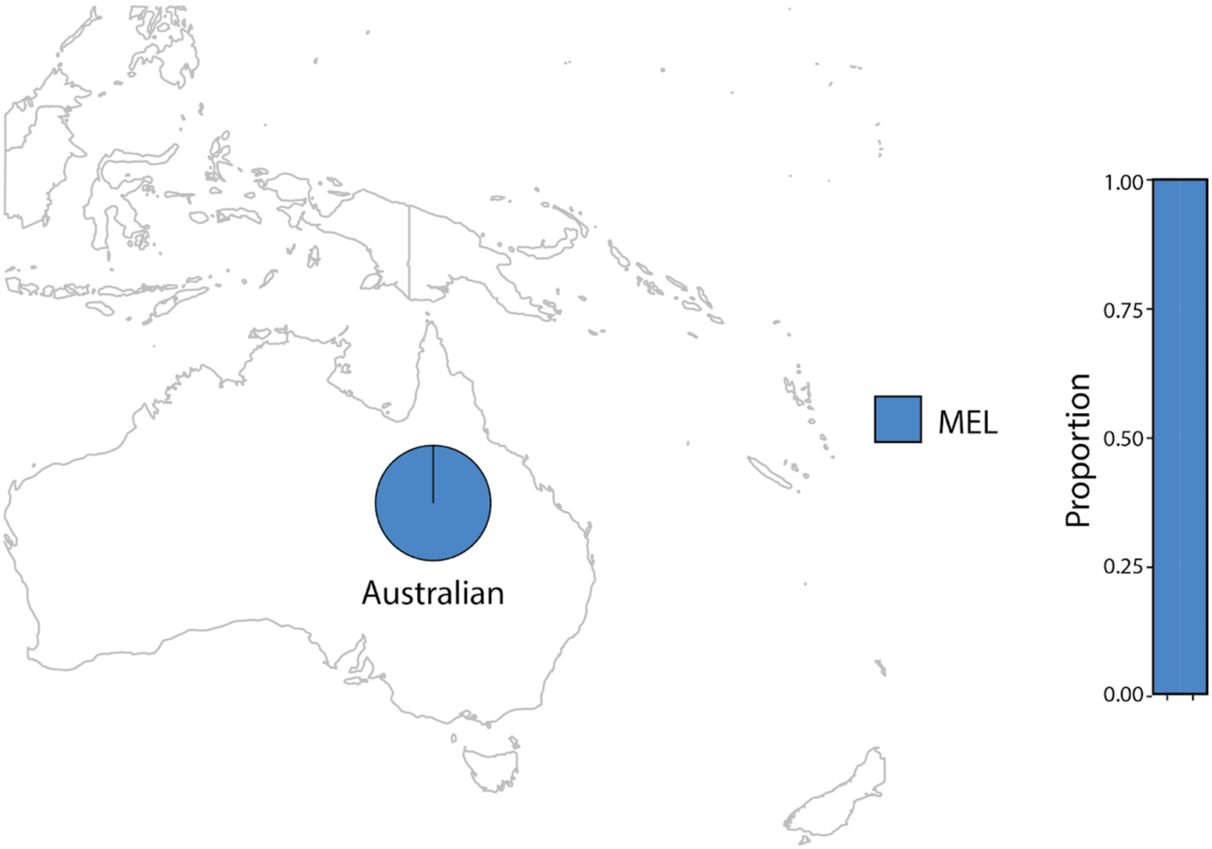


**Fig. S25.** Ancestry prediction by Orchestra for Aboriginal Australians.

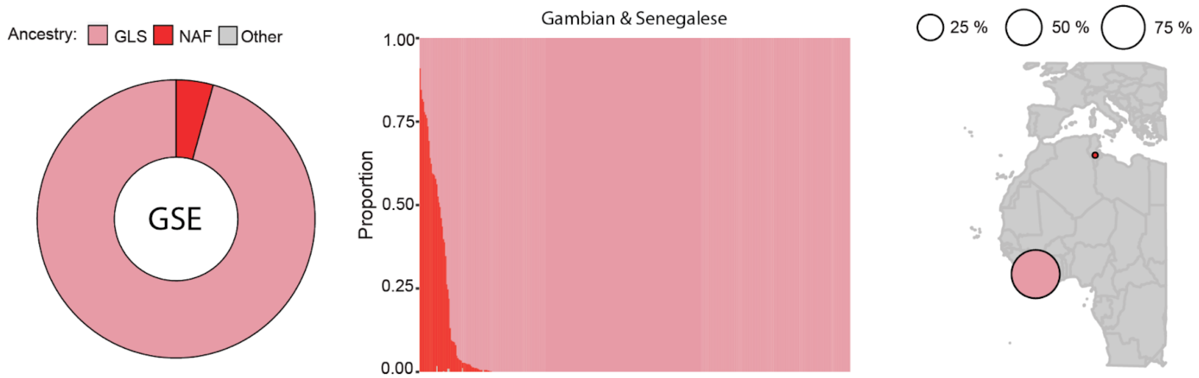


**Fig. S26. Ancestral mapping for the Gambian & Senegalese (GSE) population.** Overall ancestry proportions (left); individual ancestry proportions with each bar representing a single individual (middle); ancestry proportions shown on a map (right).

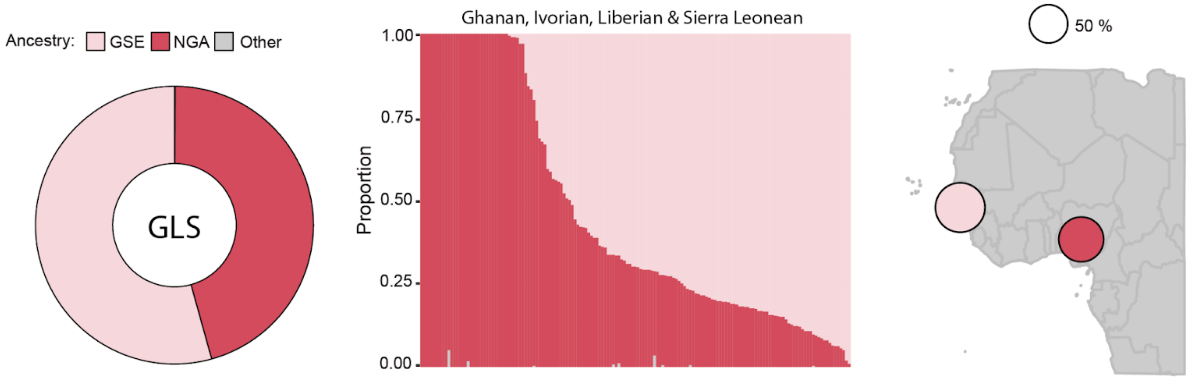


**Fig. S27. Ancestral mapping for the Ghanaian, Ivorian, Liberian & Sierra Leonean (GLS) population.** Overall ancestry proportions (left); individual ancestry proportions with each bar representing a single individual (middle); ancestry proportions shown on a map (right).

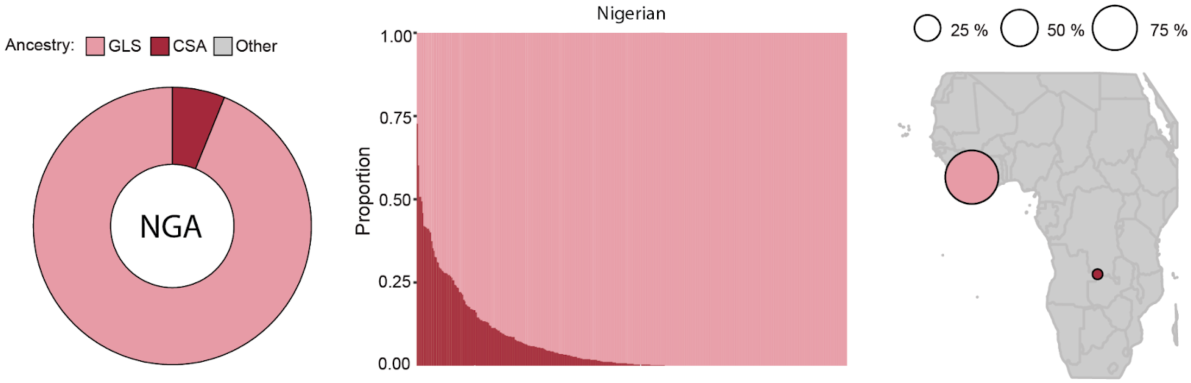


**Fig. S28. Ancestral mapping for the Nigerian (NGA) population.** Overall ancestry proportions (left); individual ancestry proportions with each bar representing a single individual (middle); ancestry proportions shown on a map (right).

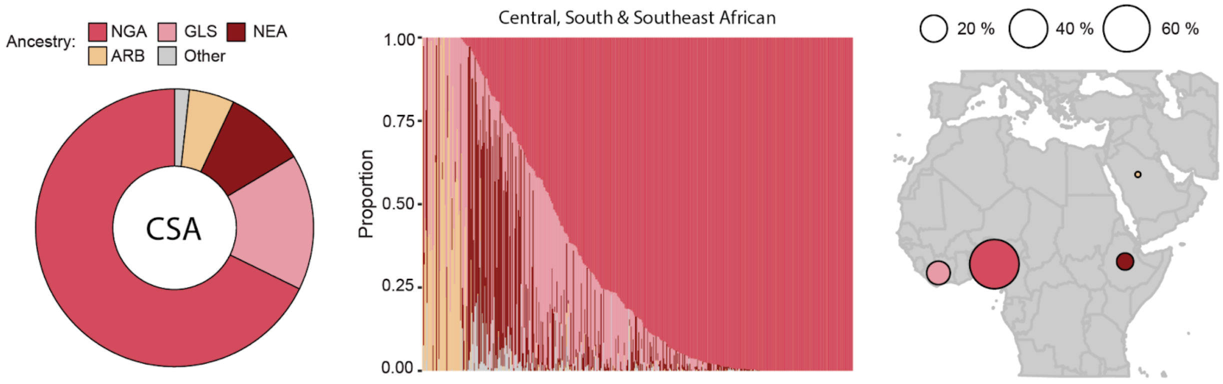


**Fig. S29. Ancestral mapping for the Central, South & Southeast African (CSA) population.** Overall ancestry proportions (left); individual ancestry proportions with each bar representing a single individual (middle); ancestry proportions shown on a map (right).

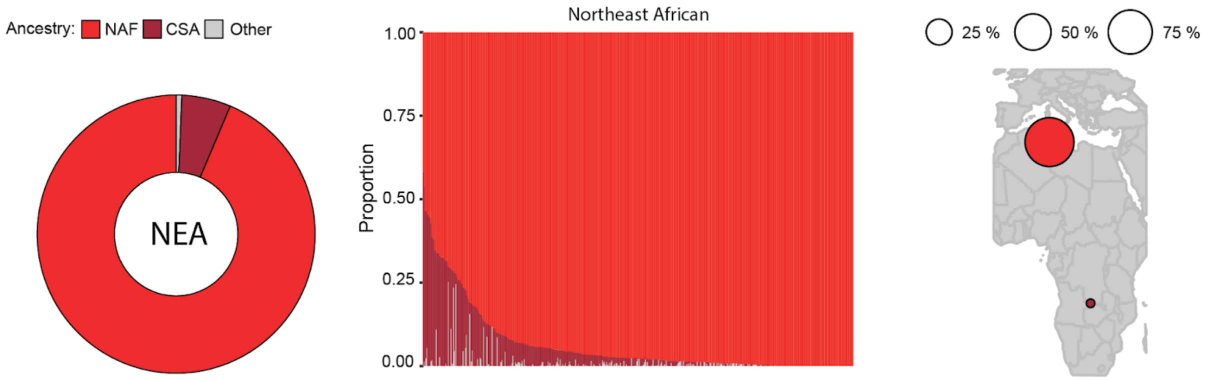


**Fig. S30. Ancestral mapping for the Northeast African (NEA) population.** Overall ancestry proportions (left); individual ancestry proportions with each bar representing a single individual (middle); ancestry proportions shown on a map (right).

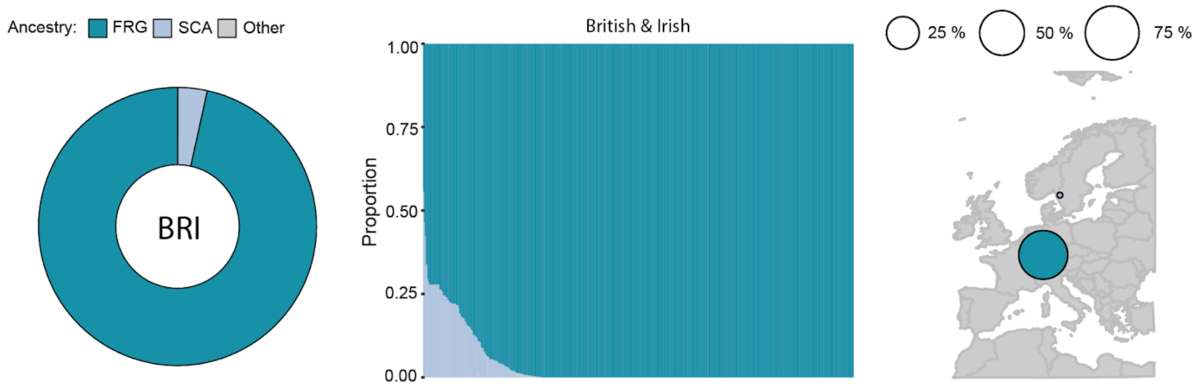


**Fig. S31. Ancestral mapping for the British & Irish (BRI) population.** Overall ancestry proportions (left); individual ancestry proportions with each bar representing a single individual (middle); ancestry proportions shown on a map (right).

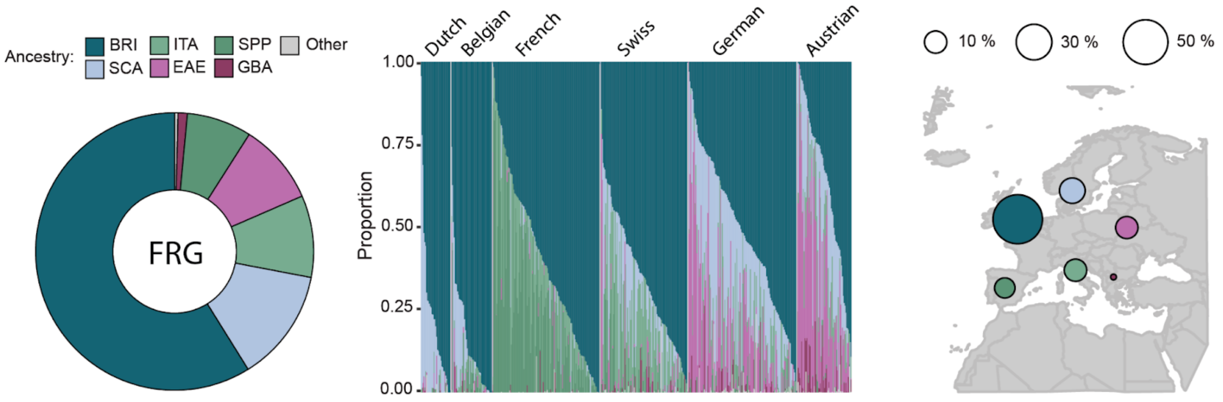


**Fig. S32. Ancestral mapping for the French & German (FRG) population.** Overall ancestry proportions (left); individual ancestry proportions shown grouped by ethnicity/subregion, with each bar representing a single individual (middle); ancestry proportions shown on a map (right).

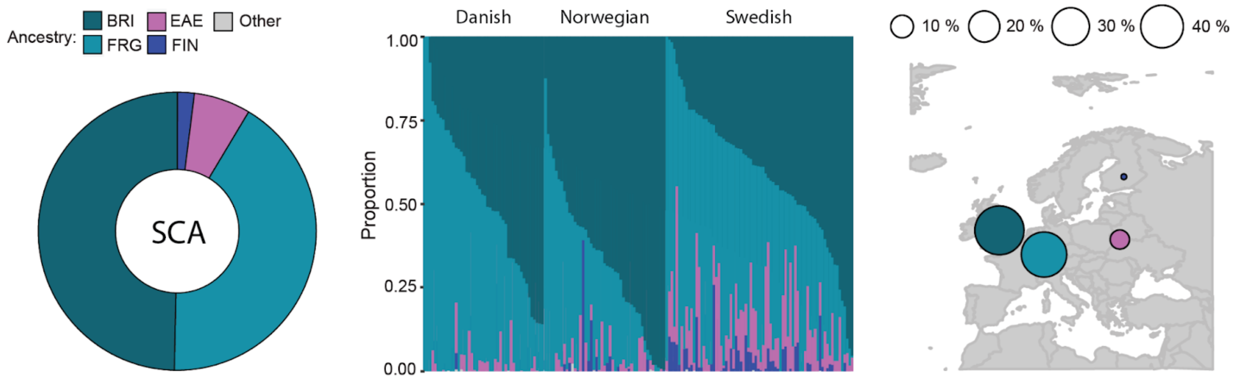


**Fig. S33. Ancestral mapping for the Scandinavian (SCA) population.** Overall ancestry proportions (left); individual ancestry proportions shown grouped by ethnicity/subregion, with each bar representing a single individual (middle); ancestry proportions shown on a map (right).

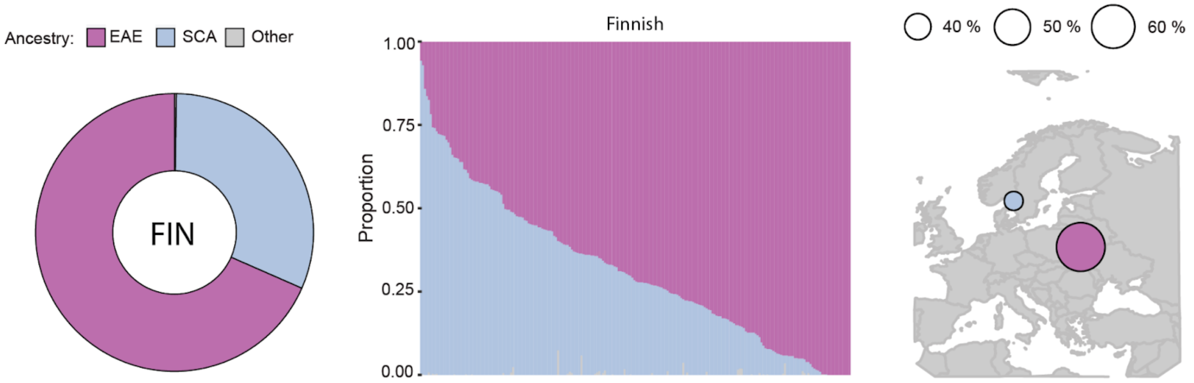


**Fig. S34. Ancestral mapping for the Finnish (FIN) population.** Overall ancestry proportions (left); individual ancestry proportions with each bar representing a single individual (middle); ancestry proportions shown on a map (right).

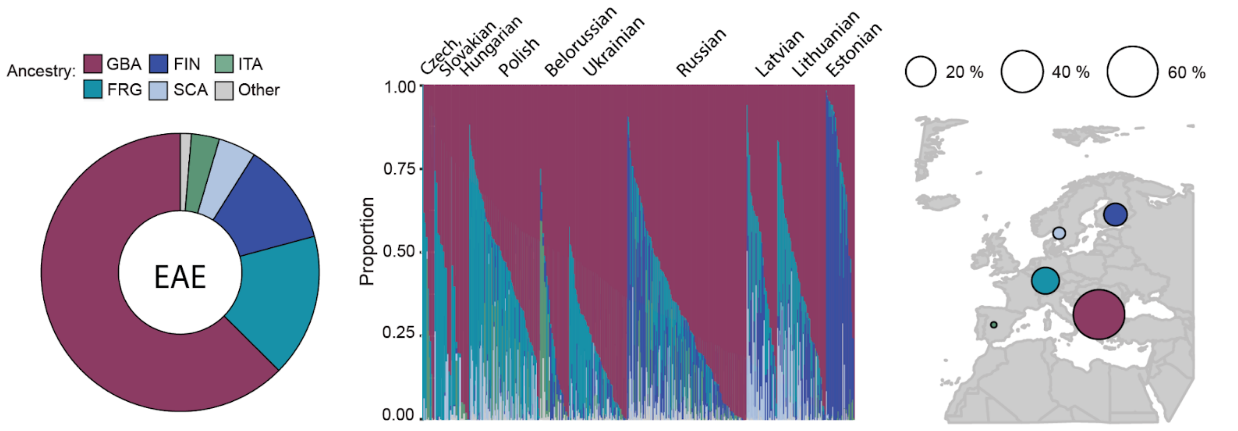


**Fig. S35. Ancestral mapping for the Eastern European (EAE) population.** Overall ancestry proportions (left); individual ancestry proportions shown grouped by ethnicity/subregion, with each bar representing a single individual (middle); ancestry proportions shown on a map (right).

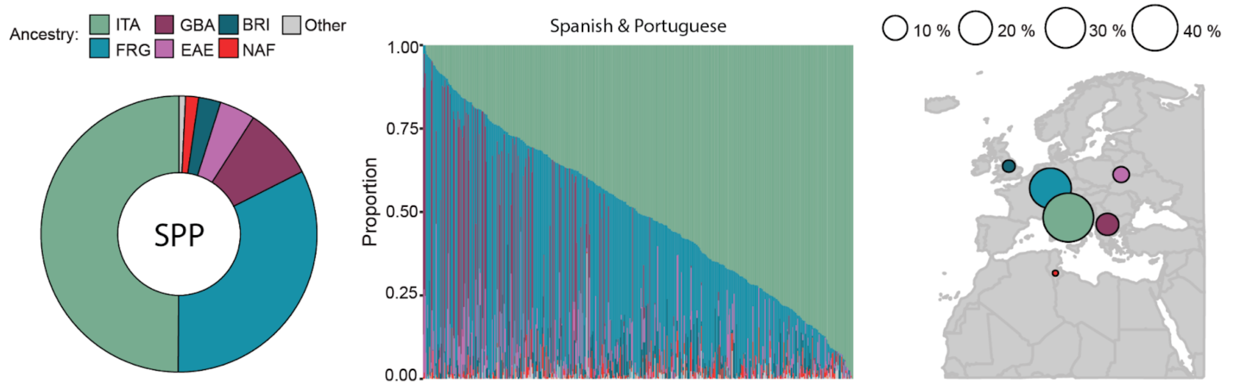


**Fig. S36. Ancestral mapping for the Spanish & Portuguese (SPP) population.** Overall ancestry proportions (left); individual ancestry proportions with each bar representing a single individual (middle); ancestry proportions shown on a map (right).

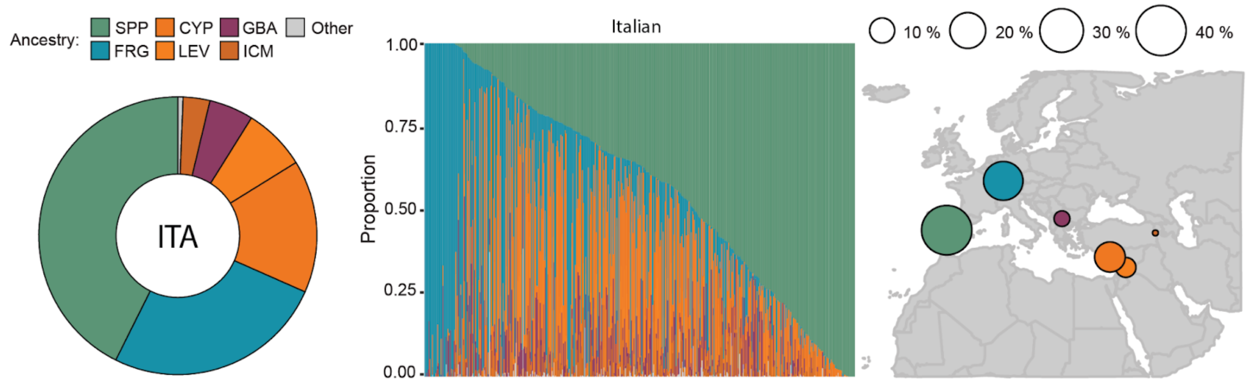


**Fig. S37. Ancestral mapping for the Italian (ITA) population.** Overall ancestry proportions (left); individual ancestry proportions with each bar representing a single individual (middle); ancestry proportions shown on a map (right).

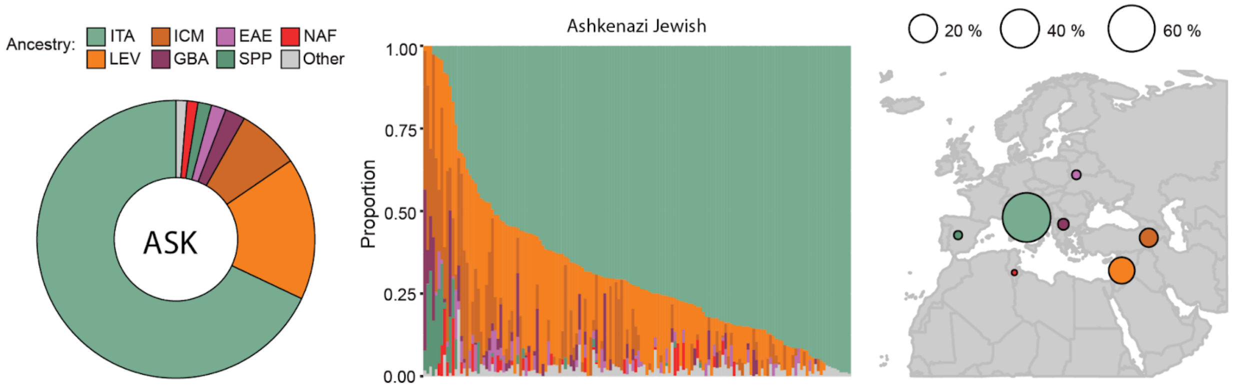


**Fig. S38. Ancestral mapping for the Ashkenazi Jewish (ASK) population.** Overall ancestry proportions (left); individual ancestry proportions with each bar representing a single individual (middle); ancestry proportions shown on a map (right).

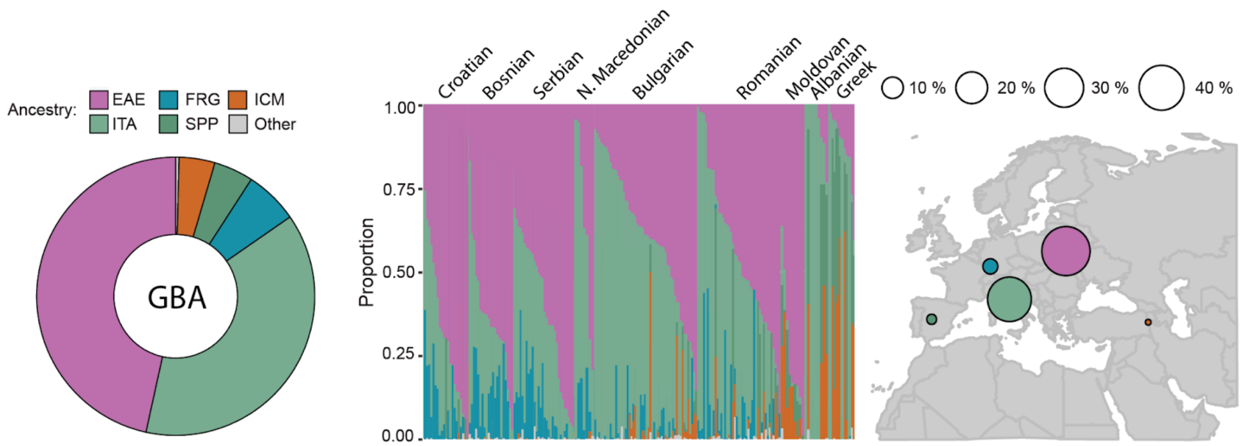


**Fig. S39. Ancestral mapping for the Greek & Balkan (GBA) population.** Overall ancestry proportions (left); individual ancestry proportions shown grouped by ethnicity/subregion, with each bar representing a single individual (middle); ancestry proportions shown on a map (right).

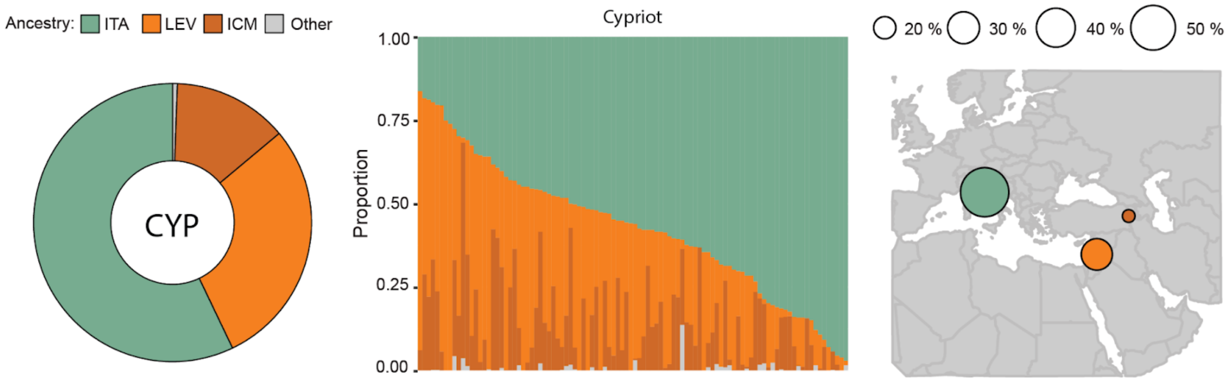


**Fig. S40. Ancestral mapping for the Cypriot (CYP) population.** Overall ancestry proportions (left); individual ancestry proportions with each bar representing a single individual (middle); ancestry proportions shown on a map (right).

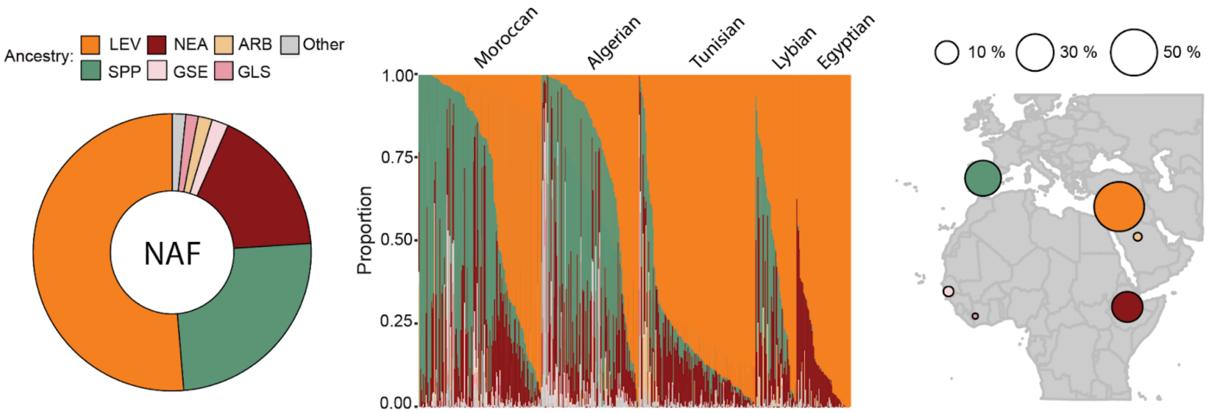


**Fig. S41. Ancestral mapping for the North African (NAF) population.** Overall ancestry proportions (left); individual ancestry proportions shown grouped by ethnicity/subregion, with each bar representing a single individual (middle); ancestry proportions shown on a map (right).

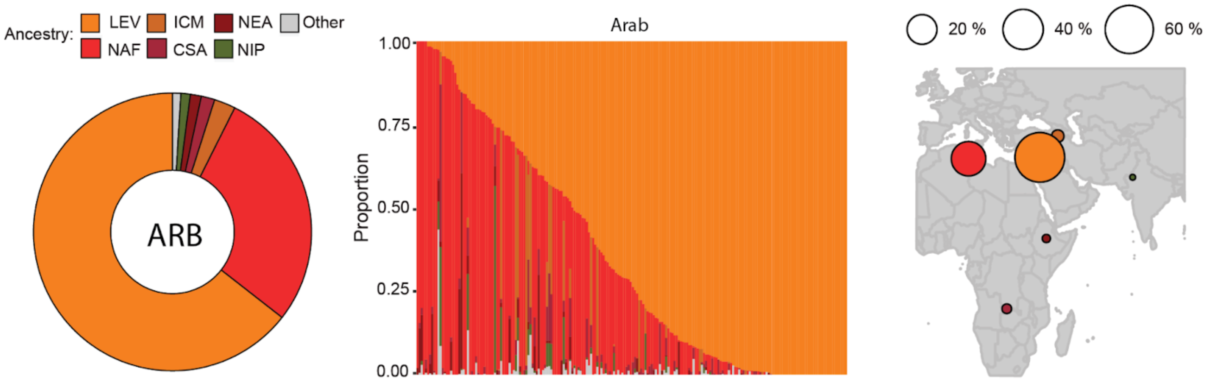


**Fig. S42. Ancestral mapping for the Arab (ARB) population.** Overall ancestry proportions (left); individual ancestry proportions with each bar representing a single individual (middle); ancestry proportions shown on a map (right).

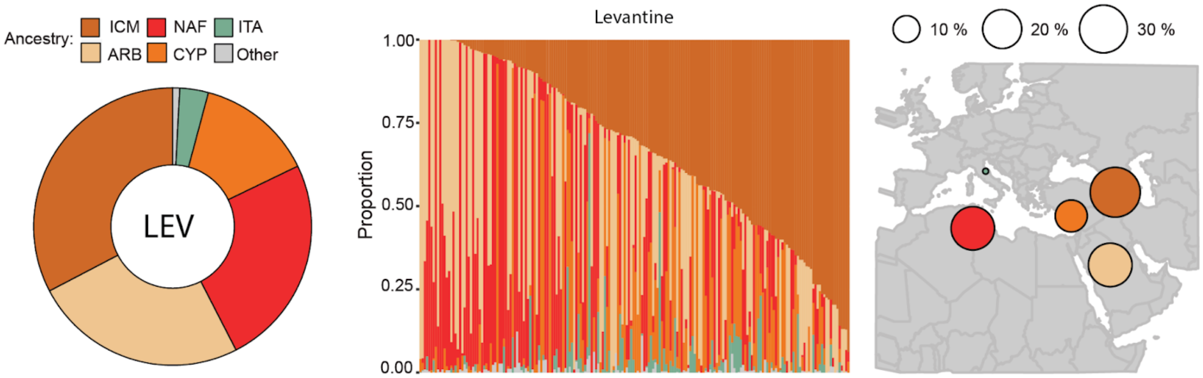


**Fig. S43. Ancestral mapping for the Levantine (LEV) population.** Overall ancestry proportions (left); individual ancestry proportions with each bar representing a single individual (middle); ancestry proportions shown on a map (right).

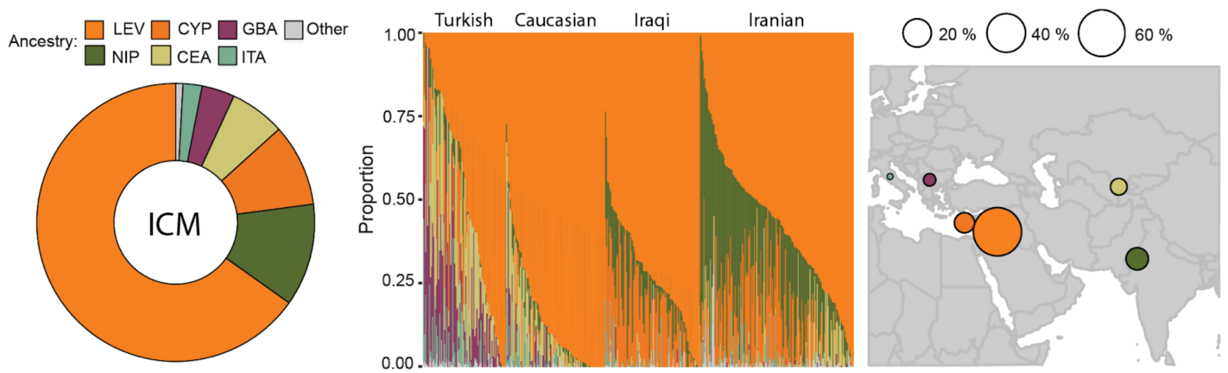


**Fig. S44. Ancestral mapping for the Turkish, Iraqi, Iranian & Caucasian (ICM) population.** Overall ancestry proportions (left); individual ancestry proportions shown grouped by ethnicity/subregion, with each bar representing a single individual (middle); ancestry proportions shown on a map (right).

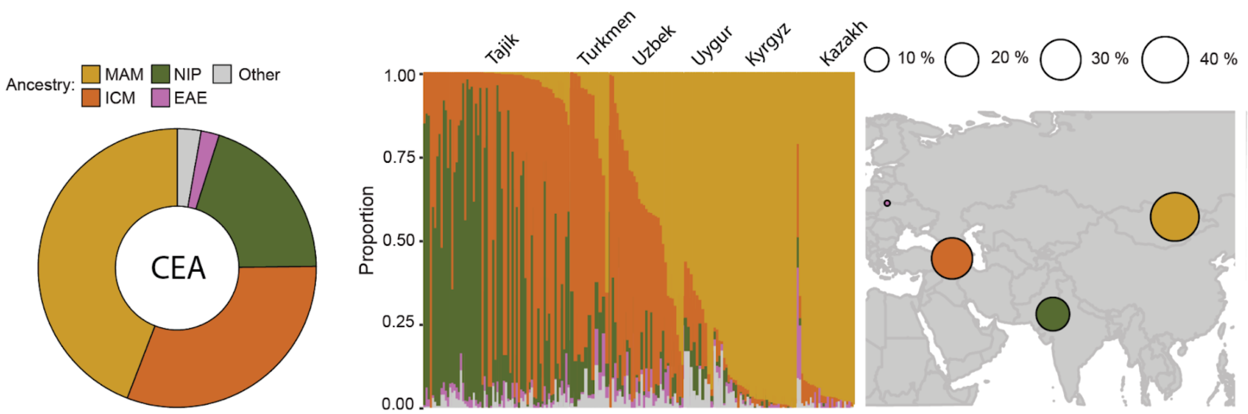


**Fig. S45. Ancestral mapping for the Central Asian (CEA) population.** Overall ancestry proportions (left); individual ancestry proportions shown grouped by ethnicity/subregion, with each bar representing a single individual (middle); ancestry proportions shown on a map (right).

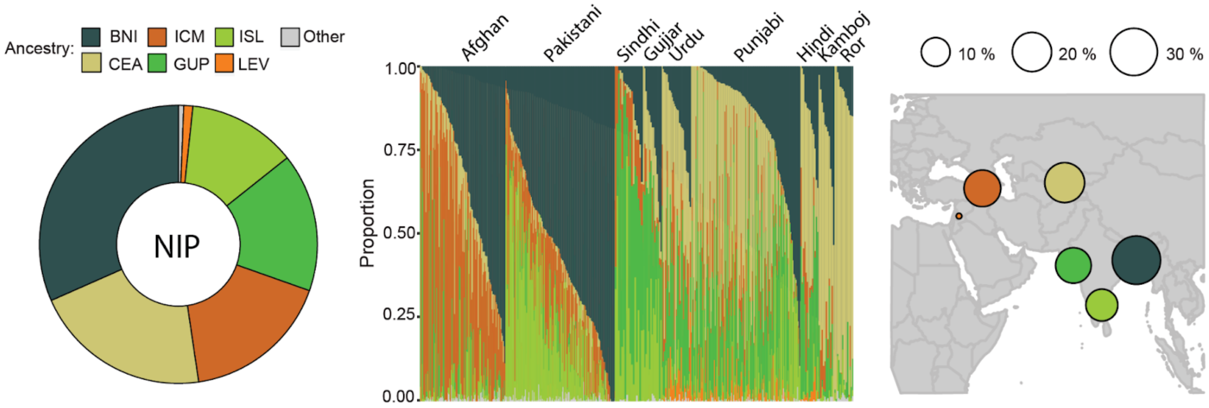


**Fig. S46. Ancestral mapping for the Northern Indian & Pakistani (NIP) population.** Overall ancestry proportions (left); individual ancestry proportions shown grouped by ethnicity/subregion, with each bar representing a single individual (middle); ancestry proportions shown on a map (right).

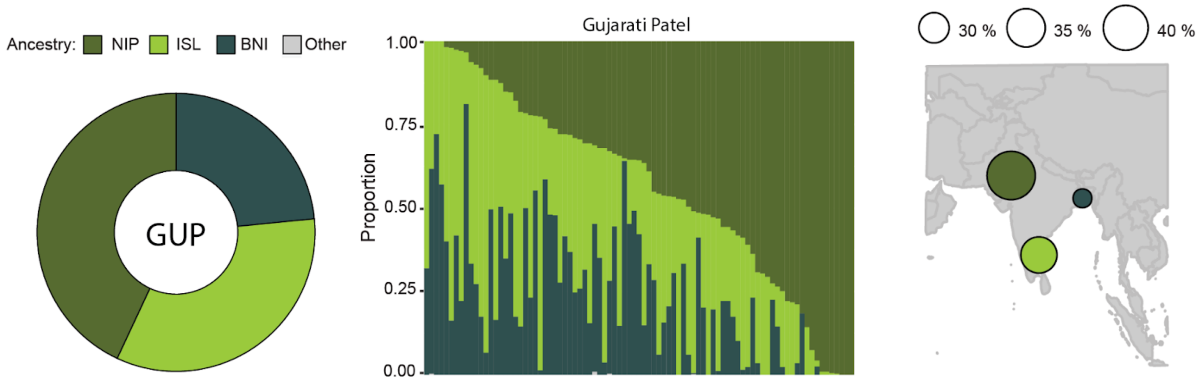


**Fig. S47. Ancestral mapping for the Gujarati Patel (GUP) population.** Overall ancestry proportions (left); individual ancestry proportions with each bar representing a single individual (middle); ancestry proportions shown on a map (right).

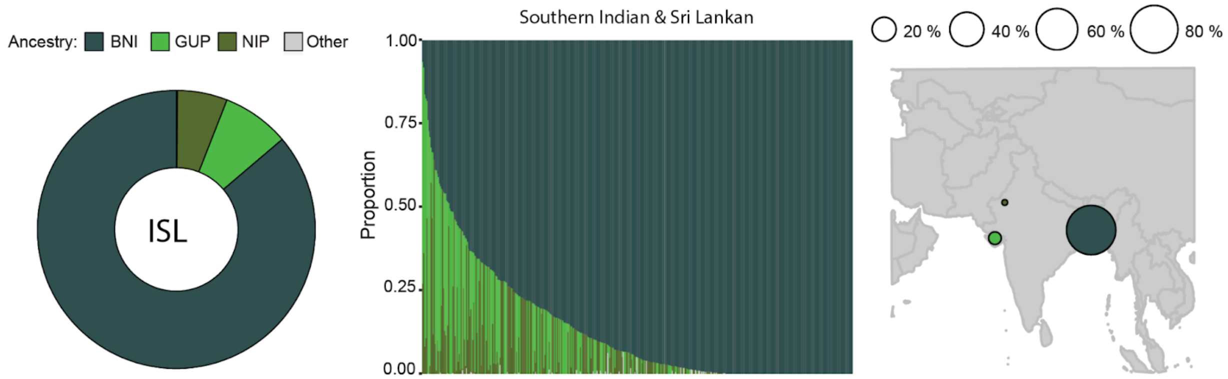


**Fig. S48. Ancestral mapping for the Southern Indian & Sri Lankan (ISL) population.** Overall ancestry proportions (left); individual ancestry proportions with each bar representing a single individual (middle); ancestry proportions shown on a map (right).

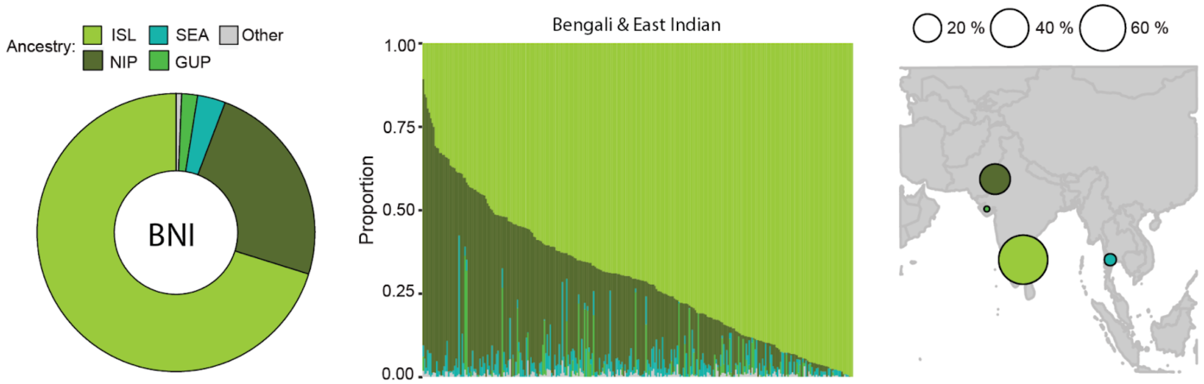


**Fig. S49. Ancestral mapping for the Bengali & East Indian (BNI) population.** Overall ancestry proportions (left); individual ancestry proportions with each bar representing a single individual (middle); ancestry proportions shown on a map (right).

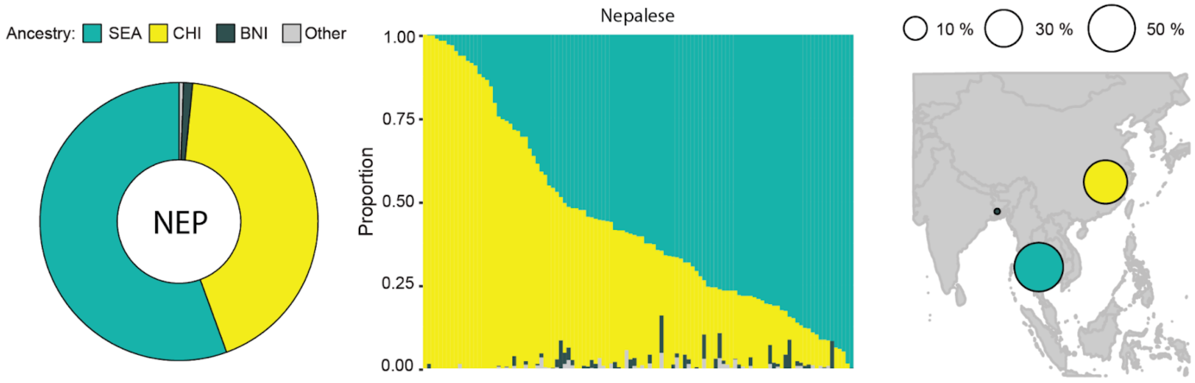


**Fig. S50. Ancestral mapping for the Nepalese (NEP) population.** Overall ancestry proportions (left); individual ancestry proportions with each bar representing a single individual (middle); ancestry proportions shown on a map (right).

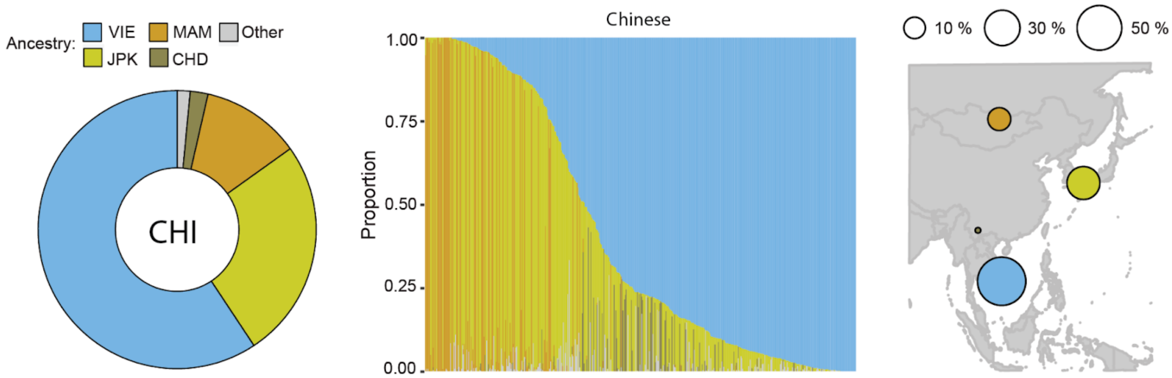


**Fig. S51. Ancestral mapping for the Chinese (CHI) population.** Overall ancestry proportions (left); individual ancestry proportions with each bar representing a single individual (middle); ancestry proportions shown on a map (right).

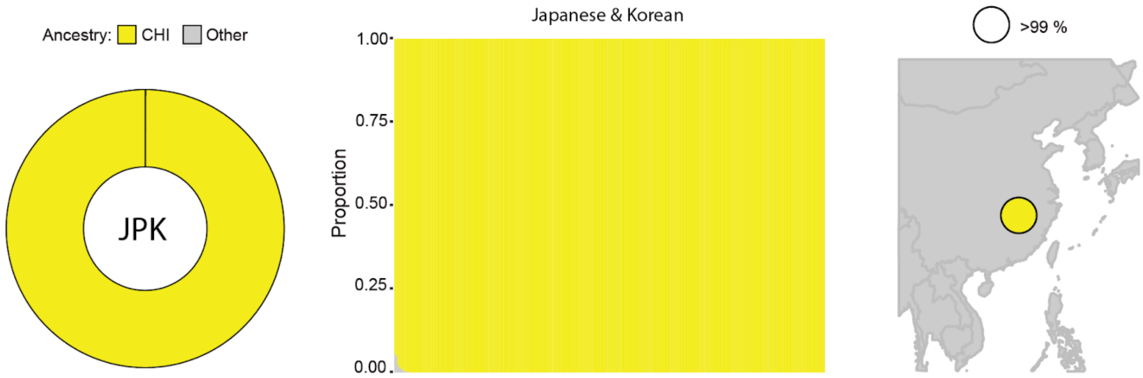


**Fig. S52. Ancestral mapping for the Japanese & Korean (JPK) population.** Overall ancestry proportions (left); individual ancestry proportions with each bar representing a single individual (middle); ancestry proportions shown on a map (right).

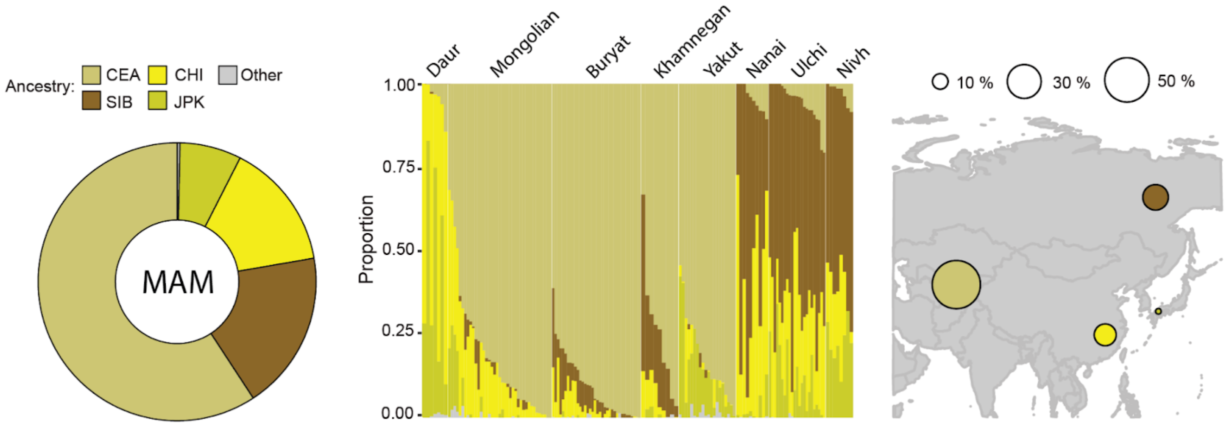


**Fig. S53. Ancestral mapping for the Manchurian & Mongolian (MAM) population.** Overall ancestry proportions (left); individual ancestry proportions shown grouped by ethnicity/subregion, with each bar representing a single individual (middle); ancestry proportions shown on a map (right).

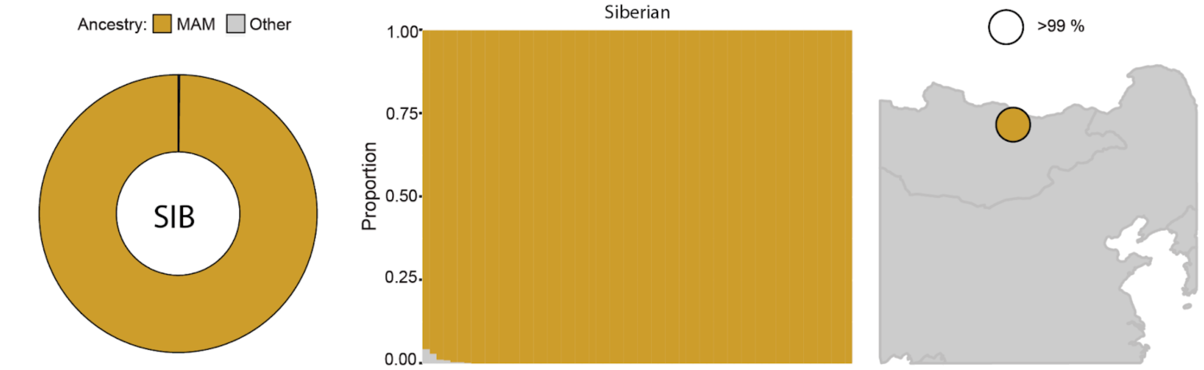


**Fig. S54. Ancestral mapping for the Siberian (SIB) population.** Overall ancestry proportions (left); individual ancestry proportions with each bar representing a single individual (middle); ancestry proportions shown on a map (right).

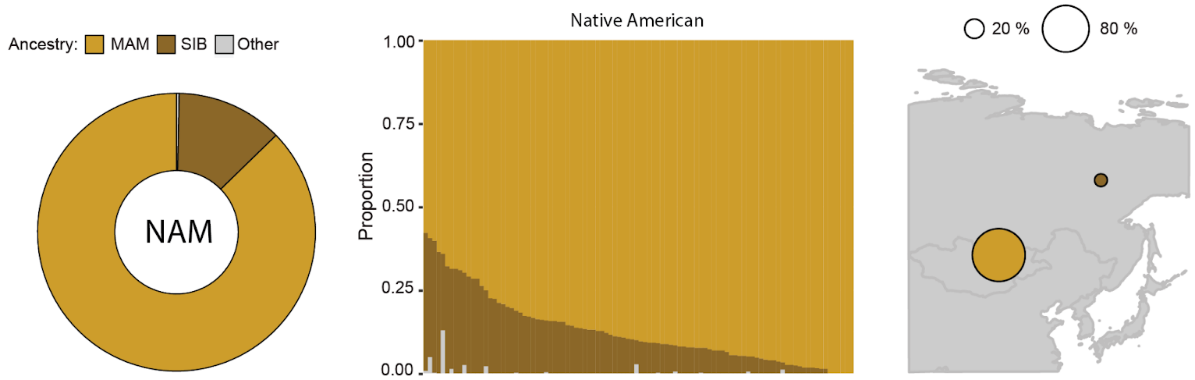


**Fig. S55. Ancestral mapping for the Native American (NAM) population.** Overall ancestry proportions (left); individual ancestry proportions with each bar representing a single individual (middle); ancestry proportions shown on a map (right).

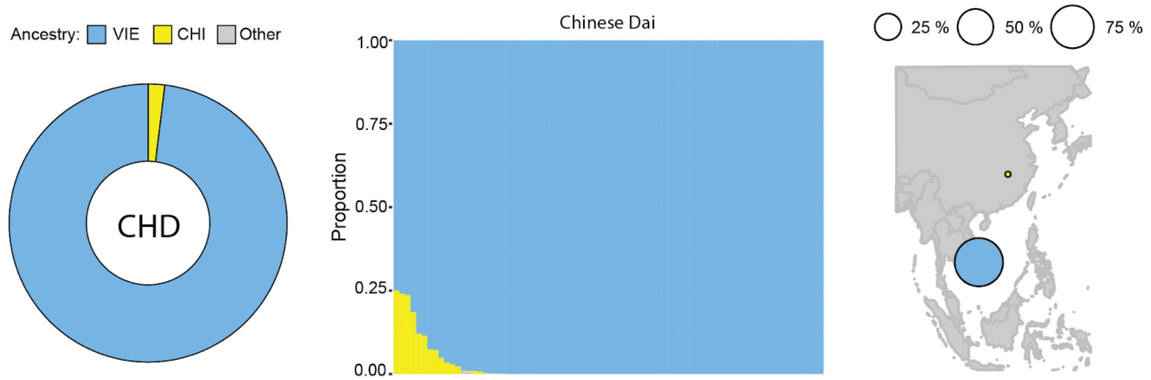


**Fig. S56. Ancestral mapping for the Chinese Dai (CHD) population.** Overall ancestry proportions (left); individual ancestry proportions with each bar representing a single individual (middle); ancestry proportions shown on a map (right).

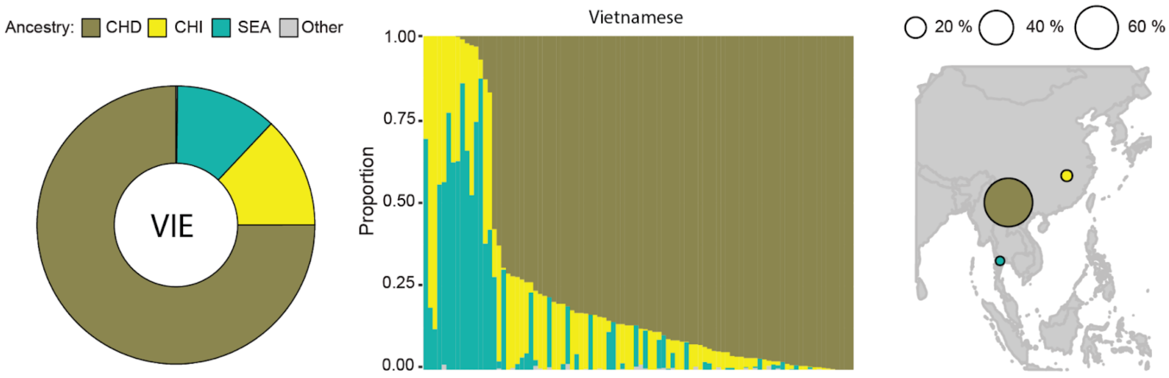


**Fig. S57. Ancestral mapping for the Vietnamese (VIE) population.** Overall ancestry proportions (left); individual ancestry proportions with each bar representing a single individual (middle); ancestry proportions shown on a map (right).

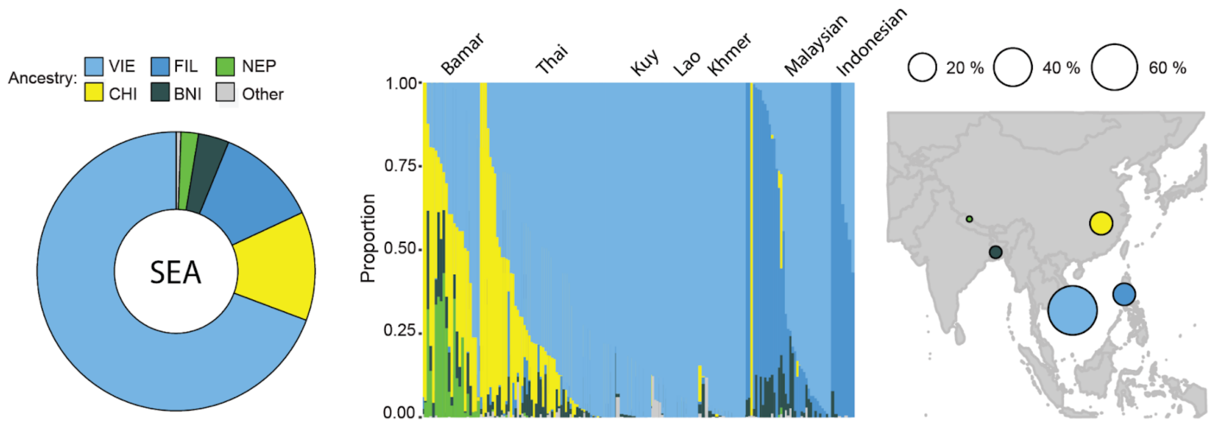


**Fig. S58. Ancestral mapping for the Southeast Asian (SEA) population.** Overall ancestry proportions (left); individual ancestry proportions shown grouped by ethnicity/subregion, with each bar representing a single individual (middle); ancestry proportions shown on a map (right).

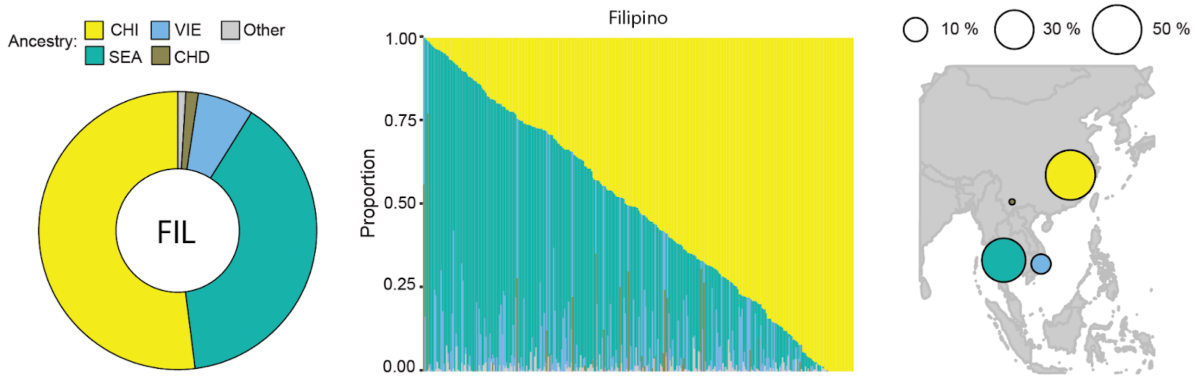


**Fig. S59. Ancestral mapping for the Filipino (FIL) population.** Overall ancestry proportions (left); individual ancestry proportions with each bar representing a single individual (middle); ancestry proportions shown on a map (right).

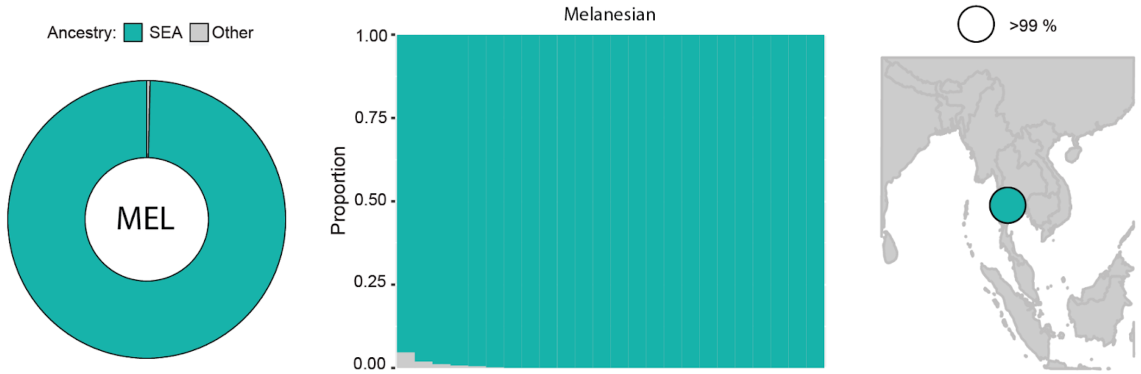


**Fig. S60. Ancestral mapping for the Melanesian (MEL) population.** Overall ancestry proportions (left); individual ancestry proportions with each bar representing a single individual (middle); ancestry proportions shown on a map (right).

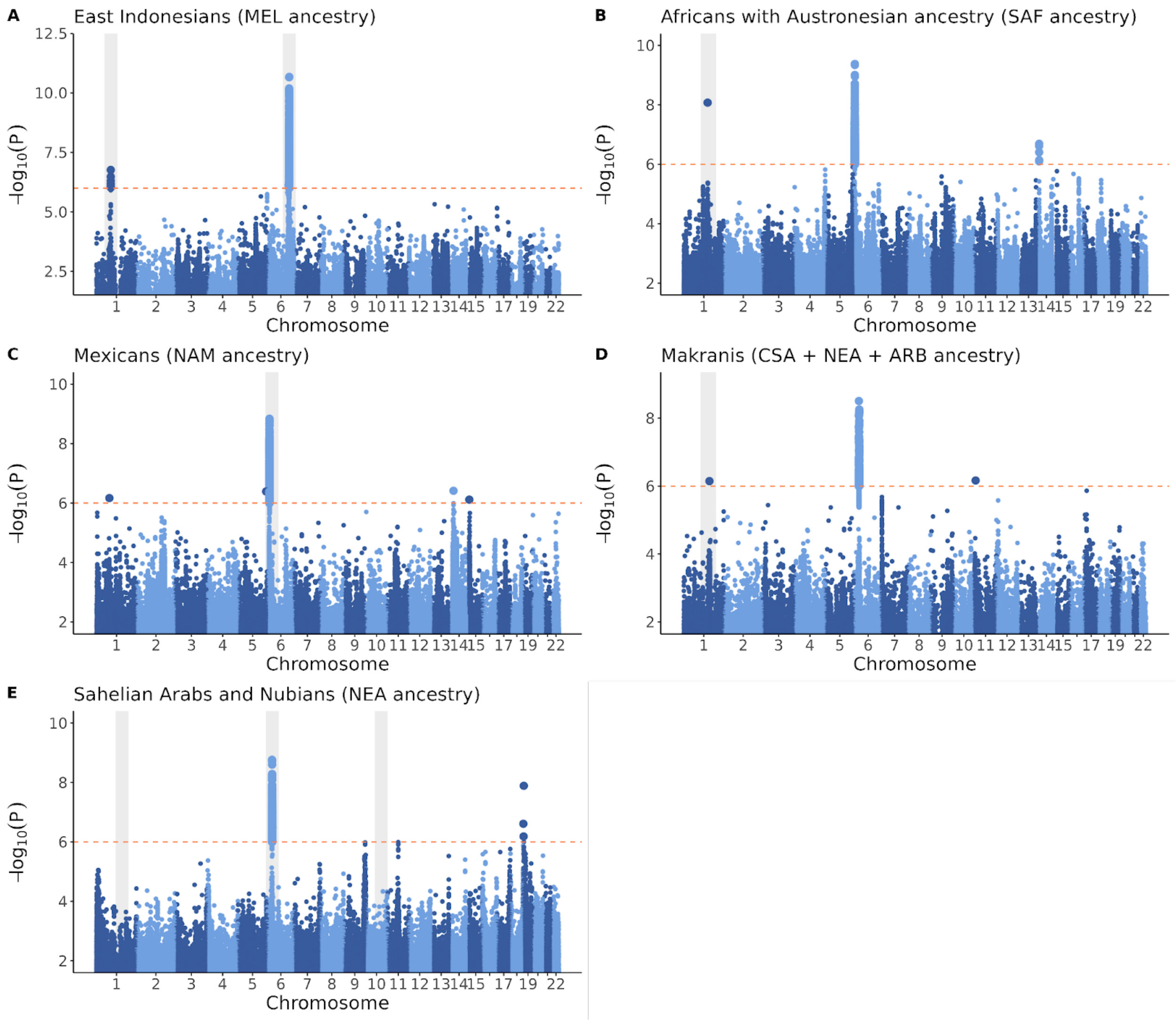


**Fig. S61. Genomic signals of adaptive admixture form Cuadros-Espinoza et al (2022) replicated in this study.** Regions shaded in gray indicate local signatures of adaptation described by Cuadros-Espinoza et al (2022). Admixed populations tested include East Indonesians (A), Africans with Austronesian ancestry as a proxy of Malagasy (B), Mexicans (C), Makranis (D) and Sahelian Arabs and Nubians (E). The assessed ancestry for each population is indicated in parentheses within the title of each panel. Larger points indicate variants that pass the established significance threshold (shown by a horizontal dotted line). See table S3 for more details.

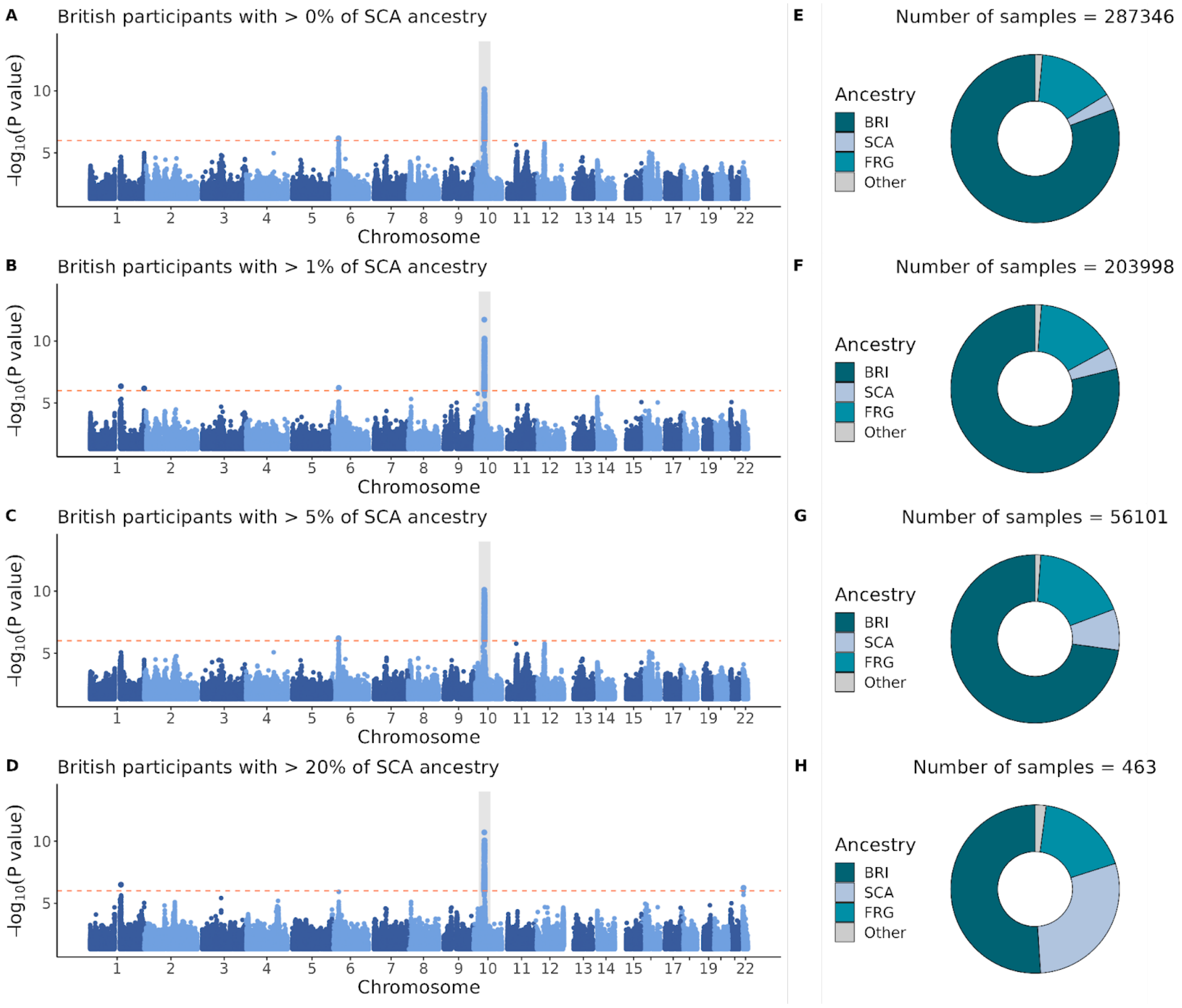


**Fig. S62. Genome-wide adaptive admixture signal in British UKBB participants with varying levels of Scandinavian ancestry.** Larger points indicate variants that pass the established significance threshold (shown by a horizontal dotted line). *P* values were derived from a combined analysis of Fadm and LAD scores as outlined in Cuadros-Espinoza et al (2022). The highlighted area in gray pinpoints the chr10 signal discovered. Genome-wide signals are presented for British sample sets with increased Scandinavian heritage (A,B,C,D), alongside their corresponding ancestry proportions and the total number of individuals selected (E,F,G,H).

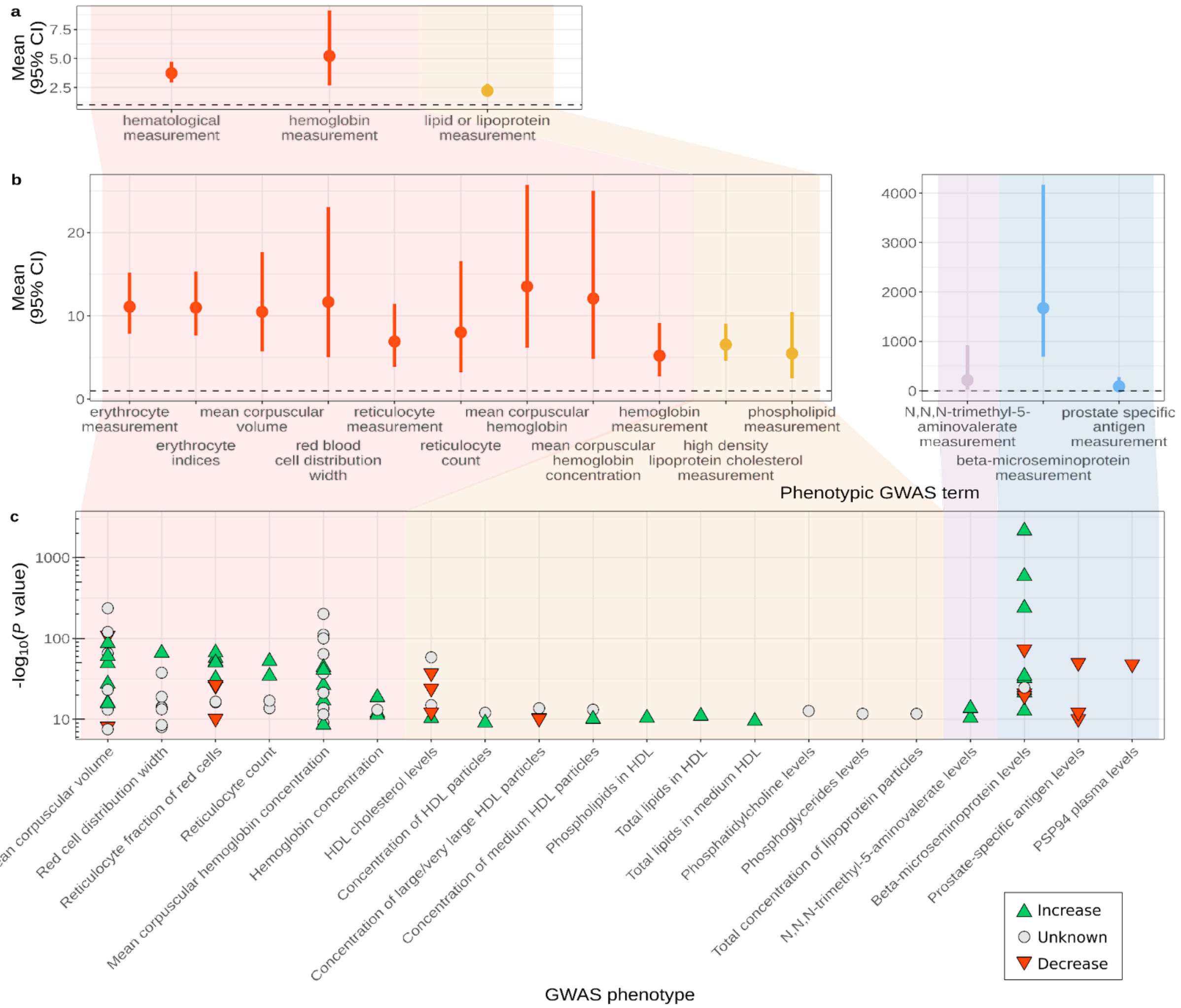


**Fig. S63. Excess of GWAS hits located in the chromosome 10 region corresponding to the identified adaptive signal.** Enrichment of parent categories (a) and GWAS phenotypes (b). The direction of the GWAS effects resulting from the presence of SCA instead of BRI ancestry was inferred from SNP frequencies (c). Discrepancies may be related to SNP frequency inaccuracies.

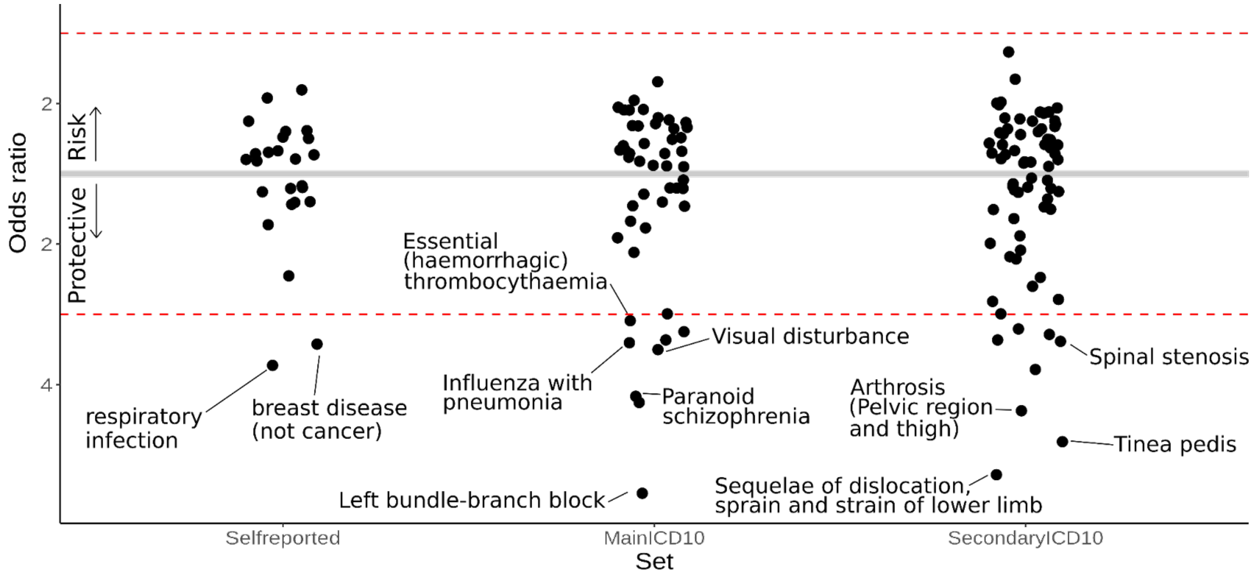


**Fig. S64. Admixture mapping**. Nominally significant UKBB phenotypes (encompassing self-reported illness codes and primary/secondary ICD10 codes) are ranked by odds ratio. 'Protective' indicates an excess of BRI ancestry among cases or SCA among controls, while 'risk' points to a higher proportion of SCA ancestry in cases.

**Table S1. Datasets included in the reference panel.** Data source, genotyping technology and corresponding population in our study.

| Data source | Description | Genotyping technology | Populations |
|---|---|---|---|
| Byrska-Bishop et al, 2022; Lowy-Gallego et al, 2019 | 1000 Genomes Project | WGS | CSA, GSE, GLS, NGA, CHD, CHI, VIE, JPK, FIN, ITA, SPP, BNI, GUP, NIP, ISL |
| Bergström et al, 2020 | Human Genome Diversity Project | WGS | CSA, GSE, NGA, CHI, SEA, JPK, MAM, MEL, ITA, SPP, ARB, LEV, NAF, CEA, NIP |
| Mallick et al, 2016 | Simons Genome Diversity Project | WGS | CSA, GSE, NGA, FIL, SIB, MEL, EAE, ITA, SPP, ARB, LEV, NAF, ICM, CEA |
| This study | Artificially constructed Native Americans | WGS (from 1KGP, HGDP and SGDP genomes) | NAM |
| Almarri et al, 2021 | Middle Eastern populations | WGS | ARB |
| Malaria Genomic Epidemiology Network, 2019 | Gambian Genome Variation Project (Fula, Jola, Mandinka and Wolof ethnic groups) | WGS | GSE |
| Zhang et al, 2014; Kim et al, 2018 | Korean Personal Genome Project | WGS (Illumina Hiseq) | JPK |
| Carmi et al, 2014 | The Ashkenazi Genome Consortium (healthy individuals of Ashkenazi Jewish descent) | WGS | ASK |
| Bycroft et al, 2018 | UK Biobank project (participants from across the United Kingdom) | UK BiLEVE Axiom and UK Biobank Axiom arrays (imputed with the Haplotype Reference Consortium, UK10K and 1000 Genomes reference panels) | CSA, NEA, GSE, GLS, NGA, CHI, FIL, SEA, VIE, JPK, MAM, EAE, BRI, FIN, FRG, SCA, GBA, ITA, SPP, LEV, NAF, CYP, ICM, CEA, NEP, BNI, NIP, ISL |
| Wang et al, 2021; Jeong et al, 2019; Biagini et al, 2019; Vyas et al, 2017; Skoglund et al, 2017; Skoglund et al, 2016; Lazaridis et al, 2016; Lazaridis et al, 2014; Pickrell et al, 2012 | Modern samples from the '1240K+HO' dataset provided by Dr David Reich laboratory | Affymetrix Human Origins array | CSA, NEA, CHI, FIL, MAM, SIB, ASK, EAE, GBA, SPP, ARB, LEV, NAF, ICM, CEA |
| Anagnostou et al, 2020 | Berbers and Arabs from Southern Tunisia | Illumina Human OmniExpressExome v 8.1 array | NAF |
| Henn et al, 2012; Arauna et al, 2017 | Berbers and Arabs from North Africa and Syria | Affymetrix 6.0 array | LEV, NAF |
| Hollfelder et al. 2017 | Sudanese and South Sudanese populations | Illumina Human Omni5MExome array | NEA |
| Dobon et al, 2015 | Populations (Arabs, Beja, Ethiopian and Nubian) from Sudanese region | Illumina Infinium Immunochip | NEA |

| | | | |
|---|---|---|---|
| Behar et al, 2013 | Ashkenazi Jews and non-Ashkenazi populations from Eastern Europe, Northeast Africa and the Caucasus | Illumina Human610-Quad, Human660W-Quad, HumanOmniExpress-12v1 730K and HumanOmni1-Quad array | NEA, ASK, EAE, ICM |
| Yunusbayev et al, 2012 | Caucasians and geographically nearby populations (Central Asia, Eastern Europe and Balkans) | Illumina 610K array | EAE, GBA, ICM, CEA |
| Yunusbayev et al, 2015 | Turkic-speaking populations from regions across Eurasia and their geographic neighbors | Illumina 550k, 610k, 650k and Human1M-Duo BeadChips | MAM, EAE, ICM, CEA |
| Behar et al, 2010 | Jewish Diaspora communities and non-Jewish neighbor populations from Europe, Asia and Africa | Illumina Human610-Quad and Human660W-Quad bead arrays | NEA, ASK, EAE, SPP, ARB, LEV, NAF, ICM, CEA |
| Tambets et al, 2018 | Uralic-speaking populations and local geographic neighbors | Illumina Human610-Quad, HumanHap650Y and Human660W-Quad BeadChip | EAE, FIN |
| Botigué et al, 2013 | Spanish from South (Andalusian) and Northwest (Galician) Spain | Affymetrix 6.0 array | SPP |
| Flores-Bello et al, 2021 | Basques (from France and Spain) and Spanish Peri-Basques | Axiom Genome-Wide Human Origins 1 Array | SPP |
| Henn et al, 2012 | Basques from Spanish Basque country | Affymetrix 6.0 array | SPP |
| Pathak et al, 2018 | Northwest Indian populations from Rajasthan and Haryana states (Gujjar, Kamboj, Ror) | Illumina HumanOmniExpress -24 BeadChip | NIP |
| Nelson et al, 2008 | Indian Asians (Gujarati, Hindi, Punjabi, Pushto and Urdu) from the Population Reference Sample (POPRES) project | Affymetrix GeneChip 500K array | GUP, NIP |
| Changmai et al, 2022 | Khmer and Kuy from mainland Southeast Asia (Thailand) | Affymetrix Human Origins SNP array | SEA |
| Tätte et al, 2019 | Lao from Laos | Illumina OmniExpress BeadChips for 650k, 710k and 730k | SEA |
| Mörseburg et al, 2016 | Island Southeast Asian populations (Malay, Igorot, Luz) | Illumina OmniExpress BeadChip for 730k | FIL, SEA |

**Table S2. 35 reference populations and associated codes.** Hierarchy is shown in three levels, corresponding to the continental or subcontinental, broader regional and sub-regional or population level.

| Continent | Region | Population/Subregion | Code | N | HEX |
|---|---|---|---|---|---|
| Sub-Saharan African | Central, South & Southeast African | Central, South & Southeast African | **CSA** | **500** | #a4293a |
| | West African | Ghanaian, Ivorian, Liberian & Sierra Leonean | **GLS** | **157** | #e59aa5 |
| | | Gambian & Senegalese | **GSE** | **388** | #f5d7db |
| | | Nigerian | **NGA** | **387** | #d1495b |
| | Northern East African | Northeast African | **NEA** | **500** | #8B0000 |
| Western Asian & North African | Arab & Levantine | Arab | **ARB** | **206** | #EEC591 |
| | | Levantine | **LEV** | **196** | #FF7F00 |
| | Northern West Asian | Turkish, Iraqi, Iranian & Caucasian | **ICM** | **451** | #CD6600 |
| | | Cypriot | **CYP** | **100** | #EE7621 |
| | North African | North African | **NAF** | **500** | #EE2C2C |
| Central & South Asian | Central Asian | Central Asian | **CEA** | **182** | #CDC673 |
| | Northern South Asian | North Indian & Pakistani | **NIP** | **500** | #556B2F |
| | | Gujarati Patel | **GUP** | **87** | #32CD32 |
| | | Bengali & East Indian | **BNI** | **288** | #2F4F4F |
| | | Nepalese | **NEP** | **111** | #00FF00 |
| | Southern South Asian | Southern Indian & Sri Lankan | **ISL** | **412** | #9ACD32 |
| European | Northwest European | British & Irish | **BRI** | **500** | #006373 |
| | | French & German | **FRG** | **500** | #008fa6 |
| | | Scandinavian | **SCA** | **167** | #B0C4DE |
| | | Finnish | **FIN** | **220** | #0000EE |
| | East European | Eastern European | **EAE** | **500** | #CD69C9 |
| | South European | Greek & Balkan | **GBA** | **209** | #8B3A62 |
| | | Italian | **ITA** | **500** | #76ab8f |
| | | Spain & Portugal | **SPP** | **500** | #5a9375 |
| | Ashkenazi Jewish | Ashkenazi Jewish | **ASK** | **152** | #CD96CD |
| East Asian | Chinese & Southeast Asian | Han Chinese | **CHI** | **500** | #FFFF00 |
| | | Chinese Dai | **CHD** | **77** | #8B864E |
| | | Filipino | **FIL** | **318** | #4F94CD |
| | | Southeast Asian | **SEA** | **167** | #20B2AA |
| | | Vietnamese | **VIE** | **94** | #63B8FF |
| | Japanese & Korean | Japanese & Korean | **JPK** | **462** | #CDCD00 |
| | Northern Asian | Siberian | **SIB** | **62** | #8B6508 |
| | | Manchurian & Mongolian | **MAM** | **152** | #CD9B1D |
| Native American | Native American | Native American | **NAM** | **100** | #edae49 |
| Melanesian | Melanesian & Aboriginal Australian | Melanesian & Aboriginal Australian | **MEL** | **24** | #1E90FF |

**Table S3. Replication of adaptive admixture signals from Cuadros-Espinoza et al (2022)**. Admixed population, datasets used, sample size (N), inferred ancestry and signal replication.

| Population (Country) | Study | Dataset | N | Source populations vs. main inferred ancestries [admixture proportions] | Signals replicated? |
|---|---|---|---|---|---|
| **Malagasy (Madagascar)** | **Cuadros-Espinosa et al. 2022** | EGAS00001002549 (Pierron et al., 2017) | 700 | East and south Bantu speakers (Luhya, Sotho, Zulu) [0.60] / Mandarese [0.40] | **YES** |
| | **This study** | Southeast African UKBB participants with >1% Austronesian heritage | 221 | BNI [0.31] / ISL [0.25] / CSA [0.17] / NIP [0.08] /FRG [0.03] / BRI [0.03] / SEA [0.03] | |
| **East Indonesians: Flores Bena, Flores Bama, Sumba, Timor, Lembata, Alor, Pantar (Indonesia)** | **Cuadros-Espinosa et al. 2022** | GSE80534 (Hudjashov et al., 2017) | 201 | Filipinos [0.58] / Papuans [0.42] | **YES** |
| | **This study** | GSE80534 (Hudjashov et al., 2017) | 201 | SEA [0.92] / FIL [0.04] / MEL [0.03] | |
| **Sahelian Arabs: Bataheen, Gaalien, Shaigia, Messiria & Nubians: Danagla, Mahas, Halfawieen (Sudan)** | **Cuadros-Espinosa et al. 2022** | http://jakobssonlab.iob.uu.se/data/ (Hollfelder et al., 2017) | 97 | East Africans (Gumuz, Somali) and Nile peoples (Baria, Dinka, Nuer, Shilluk) [0.50] / South Europeans (Tuscans, Sardinians) and Druze and Bedouins [0.50] | **Partially** |
| | **This study** | http://jakobssonlab.iob.uu.se/data/ (Hollfelder et al., 2017) | 97 | NEA [0.55] / ARB [0.34] / CSA [0.11] | |
| **Cosmopolitan Mexicans (Mexico)** | **Cuadros-Espinosa et al. 2022** | Moreno-Estrada et al., 2014 | 418 | Mexican Native Americans (Maya Campeche, Tepehuano, north Zapotec) [0.54] / Iberian populations from Spain (IBS) [0.42] / Sub-Saharan Africans (YRI) [0.04] | **YES** |
| | **This study** | Mexicans from UKBB | 75 | NAM [0.35] / SPP [0.34] / ITA [0.16] / BRI [0.05] / FRG [0.04] | |
| **Peruvians from Lima (Peru)** | **Cuadros-Espinosa et al. 2022** | 1KGP (The 1000 Genomes Project Consortium, Nature 2015) | 85 | Peruvian Native Americans (Quechua, Aymara) [0.77] / Iberian populations from Spain (IBS) [0.20] / Sub-Saharan Africans (GWD, LWK, YRI) [0.03] | **No** |
| | **This study** | Peruvians from UKBB | 60 | NAM [0.47] / SPP [0.019] / ITA [0.13] / BRI [0.07] /FRG [0.04] | |
| **Fulani (Burkina Faso)** | **Cuadros-Espinosa et al. 2022** | E-MTAB-8434 (Vicente et al., 2019) | 53 | west Africans (Jola, Igbo) [0.77] / north Africans (Mozabite) + Europeans (CEU, Czech) [0.23] | **No** |
| | **This study** | E-MTAB-8434 | 53 | GSE [0.92] / NGA [0.05] / NAF [0.02] | |
| **Makranis, Makrani Baluch (Pakistan)** | **Cuadros-Espinosa et al. 2022** | EGAS00001002558 (Laso-Jadart et al. 2017) | 46 | Baluch [0.85] /east and south Bantu speakers (Luhya, Sotho) [0.15] | **YES** |
| | **This study** | Makrani samples from Reich lab | 24 | NIP [0.73] / ICM [0.12] / ARB [0.09] / CSA [0.04] /ISL [0.01] | |
